# COMET: A toolkit for composing customizable genetic programs in mammalian cells

**DOI:** 10.1101/769794

**Authors:** Patrick S. Donahue, Joseph W. Draut, Joseph J. Muldoon, Hailey I. Edelstein, Neda Bagheri, Joshua N. Leonard

## Abstract

Engineering mammalian cells to carry out sophisticated and customizable genetic programs requires a toolkit of multiple orthogonal and well-characterized transcription factors (TFs). To address this need, we developed the COmposable Mammalian Elements of Transcription (COMET)—an ensemble of TFs and promoters that enable the design and tuning of gene expression to an extent not previously possible. COMET currently comprises 44 activating and 12 inhibitory zinc-finger TFs and 83 cognate promoters, combined in a framework that readily accommodates new parts. This system can tune gene expression over three orders of magnitude, provides chemically inducible control of TF activity, and enables single-layer Boolean logic. We also develop a mathematical model that provides mechanistic insights into COMET performance characteristics. Altogether, COMET enables the design and construction of customizable genetic programs in mammalian cells.

## INTRODUCTION

The construction of synthetic genetic programs has emerged as a powerful approach for investigating signaling and regulatory networks^1^ and for engineering cell-based therapeutic and diagnostic devices^2, 3^. Applications in mammalian cells often involve designing new ways for cells to sense and respond to internal states or environmental cues. To achieve these functions, most programs utilize transcriptional regulation. However, while large libraries of components such as transcription factors (TFs) and promoters have been developed for prokaryotes^4^, a dearth of analogous parts for mammalian systems currently limits both fundamental research and applications in medicine.

Early synthetic TFs that have been implemented in eukaryotic cells are based on the bacterial tetracycline-responsive repressor, TetR^5, 6^, and the yeast transcriptional regulator, Gal4^7^, and these proteins remain workhorses for mammalian cell biology research. However, since only one TetR-responsive promoter and one Gal4-responsive promoter can be used simultaneously, they are not scalable for building large genetic programs. Recently, several new families of TFs have been developed to overcome this limitation through programmable DNA sequence recognition. They include zinc finger (ZF)-TFs^8, 9^, transcription activator-like effectors (TALEs)^10–12^, dCas9-TFs^13^, and TetR family regulators^14^. Proteins that bind DNA via ZF motifs are especially attractive TFs for the foundation of a comprehensive toolkit for transcriptional control in mammalian cells, as they have a well-characterized structure-function relationship, can be modified to bind specific DNA sequences, and are the smallest of these new TFs. This last property is particularly useful because gene delivery vehicles such as lentivirus and transposon vectors are constrained by cargo size; smaller TFs afford space for more transcriptional units and more complex genetic programs.

From a cell engineering perspective, an ideal transcriptional toolkit would include well-characterized TFs and promoters; a physical understanding of how design choices impact performance characteristics; and a quantitative framework that describes how such biological parts may be combined to produce intended behaviors. Such a toolkit should include: (i) multiple activating and inhibitory TFs that are orthogonal to one another and to endogenous mechanisms of regulation; (ii) sets of TFs and promoters that enable one to experimentally “scan” through values of a given performance characteristic; and (iii) modularity in TF and promoter design to enable swapping and expansion of the toolkit and interfacing with other biological parts.

To address these needs, in this study, we developed and characterized the COmposable Mammalian Elements of Transcription (COMET)—an ensemble of engineered promoters and modular ZF-TFs with tunable properties. We incorporated into COMET a panel of 19 TFs that were originally developed to regulate gene expression in yeast^15^ based on ZF domains that were designed using the oligomerized pool engineering (OPEN) protocol^16^. We first employed these TFs to characterize new promoters, and then we appended new functional domains onto the ZFs. In doing so, we elucidated design rules for utilizing TFs and promoters to build gene expression programs exhibiting customizable activation, inhibition, small molecule responsiveness, and Boolean logic in mammalian cells, and we developed a mathematical model to describe the properties of these genetic parts and programs.

## RESULTS

### Identifying promoter design rules in mammalian cells

#### ZFa activate transcription in mammalian cells

In nature, TFs based on ZF domains coordinate a variety of functions.^17^ For example, the evolutionarily ancient and widely expressed TF SP1 contains three Cys2-His2-type ZF motifs (generally considered a minimal ZF domain), and SP1 binding sites appear as tandem arrays in genes regulating cell growth, apoptosis, and immune function, as well as in compact, dynamically regulated viral promoters such as the long terminal repeat of HIV^18^. To begin developing a toolkit for constructing a variety of transcriptional programs from basic parts, we first considered the use of five synthetic ZF domains characterized in yeast by Khalil et al.,^15^ and we investigated whether these tools could be adapted to regulate transcription in mammalian cells. In this mammalian library, each TF comprises a ZF DNA-binding domain fused to the VP16 activation domain (AD), forming a ZF activator (ZFa) that recruits RNA polymerase II (RNAPII) and induces transcription^19^. A new cognate promoter was generated for each ZFa by placing one ZF binding site upstream of the YB_TATA minimal promoter (**Fig. 1a**), which confers low background and inducible expression in several cell types^20^. Sequences for reporters and TFs are provided in **Supplementary Tables 1–8**. When transfected into HEK293FT cells, all five ZFa induced expression from their cognate reporters between 4 and 17-fold above background (ZFa-independent) expression (**Fig. 1b, Supplementary Fig. 1a**). Interestingly, the rank order of the magnitudes with which these ZFa induced their cognate reporters differed from that observed in a similar system in yeast^15^.

**Fig. 1.**
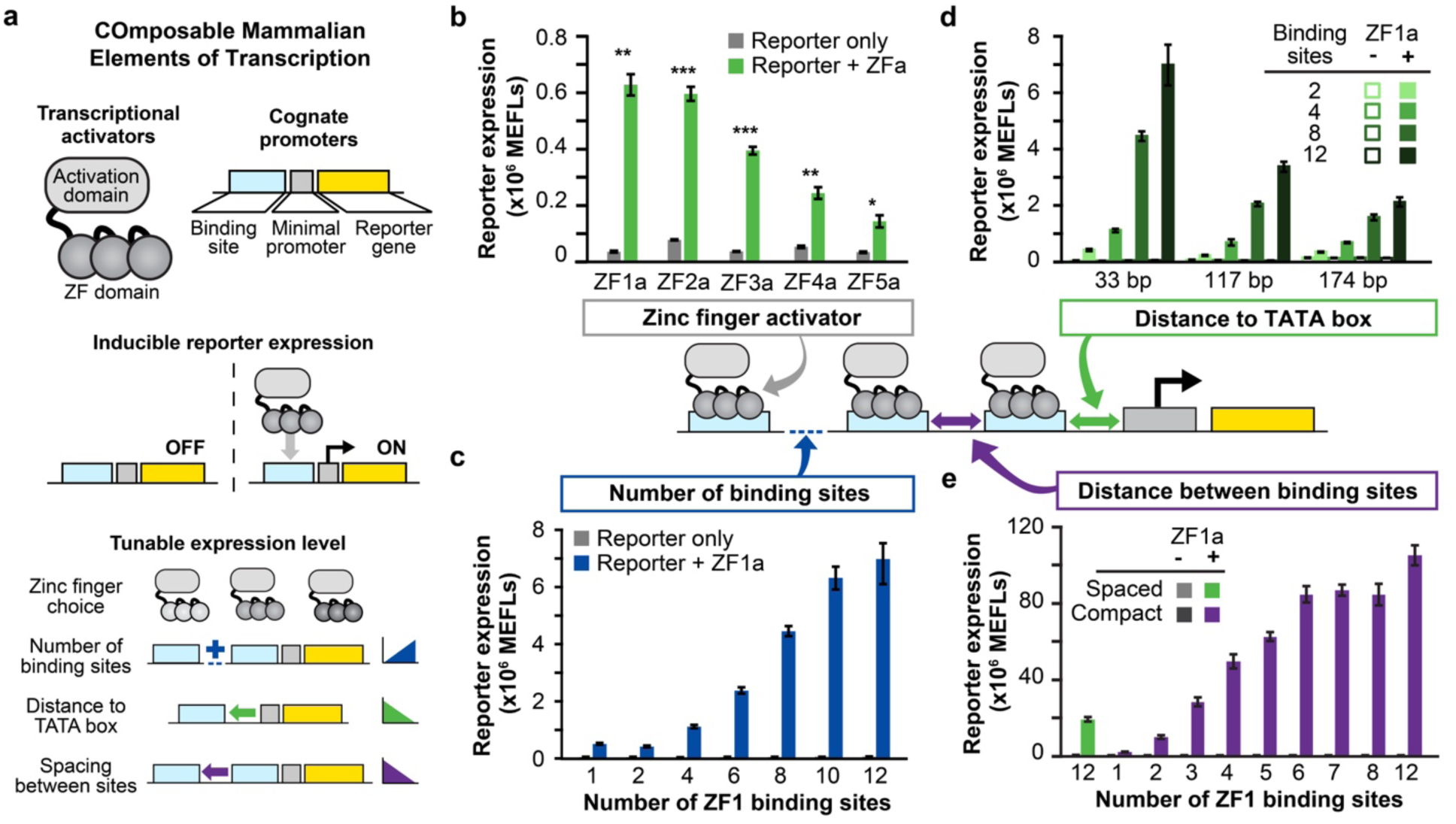
Investigation of COMET promoter design rules. **a** The schematic shows the modular, tunable features of COMET TFs and promoters. **b** Five ZFa with different ZF domains all induced reporter expression (one-tailed Welch’s *t*-test: \**p* < 0.05, \*\**p* < 0.01, \*\*\**p* < 0.001). **c** Increasing the number of ZF binding sites increased the level of gene expression in the presence of ZFa (ANOVA *p* < 0.001) but not without ZFa (ANOVA *p* = 0.24). Reporter expression increased significantly from 6 to 8 and from 8 to 10 binding sites but not on either side of this range (Tukey’s HSD test with α = 0.05). **d** Moving the ZF binding site array further upstream of the TATA box reduced reporter expression (two-factor ANOVA *p* < 0.001), and arrays with more binding sites showed more substantial decreases in reporter expression. **e** Compaction of ZFa binding sites enhanced ZFa-induced reporter expression, for an equivalent number of ZF binding sites (one-tailed Welch’s *t*-test, *p* = 0.002), and across compact promoters, ZFa-induced reporter expression increased with the number of binding sites (ANOVA *p* < 0.001). Reporter expression increased significantly from 2 to 3, 3 to 4, 5 to 6, and 8 to 12 binding sites (Tukey’s HSD test with α = 0.05). Experiments were conducted in biologic triplicate, and data were analyzed as described in **Online Methods**. Error bars represent the S.E.M.

#### The number and placement of binding sites affect maximal gene expression

Reporter fluorescence driven by the initial panel of promoters was relatively dim (on the order of 10^5^ Molecules of Equivalent Fluorescein (MEFLs, an absolute unit of fluorescence^21^) per cell), so we explored strategies for building stronger inducible promoters. An established principle is that inducible gene expression increases with the number of TF binding sites, so we tested a panel of ZF1a-responsive promoters containing multiple ZF1 sites in an array upstream from the minimal promoter (**Fig. 1c, Supplementary Fig. 1b**). In general, ZF1a-inducible reporter expression increased with the number of sites while background reporter expression was unaffected. The ZF1 promoter with 12 sites (ZF1×12) yielded 113-fold induction—approximately 12 times greater than the ZF1×1 promoter.

We hypothesized that inducible expression might also be influenced by the distance between the ZF binding site array and the TATA box, potentially promoting favorable interactions with RNAPII or conferring steric occlusion of RNAPII. To test for this effect, we constructed promoters in which the binding site array was moved from 33 bp upstream of the TATA box (the original position) to 117 bp or 174 bp upstream. Overall, increasing the spacing led to decreased expression, and this effect was especially pronounced for promoters with many sites (**Fig. 1d**). We conclude that when ZFa bind to sites that are farther from the promoter, the AD becomes too distant to contribute substantially to transcriptional activation. This explanation would also account for the diminishing returns observed with the addition of binding sites to large arrays such as 10-site and 12-site promoters (**Fig. 1c**). Thus, increasing the number of binding sites enhances gene expression, but only if the sites are near the TATA box.

#### Compacting the binding sites increases gene expression

Based upon the observation that promoters with binding sites near the TATA box were the most responsive to ZFa, we investigated whether compacting binding sites near the minimal promoter could potentiate transcriptional output. Our initial constructs had 16 to 38 bp spacers between each 9 bp binding site. To generate a more compact structure, constructs were generated with 6 bp spacers, such that ZFa would bind 15 bp apart in a rotating configuration around the DNA, as one turn of the double helix is 10.5 bp. We hypothesized that this configuration could avoid steric occlusion while increasing the local concentration of ZFa. A panel of “compact” promoters was generated, each containing 1–12 binding sites in an array beginning 33 bp upstream of the TATA box. These new promoters yielded a strong fluorescent readout of up to 10^8^ MEFLs and 360-fold induction over background (**Fig. 1e, Supplementary Fig. 1c**). The output of the strongest compact promoter (ZF1×12-C) was over five-times greater than that of the spaced promoter with the same number of binding sites (ZF1×12-S). Notably, no promoter modifications significantly altered the background level of transcription, which was low across all constructs. Also notable was that several of the strongest promoters exhibited mild squelching—a phenomenon in which inducing the expression of a TF (here, ZFa, which is expected to induce expression of the EYFP reporter) causes unexpected diminishment in the expression of a gene (here, the constitutively expressed EBFP2 transfection control) by competing for a limited pool of cellular resources that carry out transcription^19, 22^. In our data, these effects are apparent when populations of cells with high EYFP expression have lower EBFP2 expression than do populations of cells with lower EYFP expression (**Supplementary Fig. 1c**). Together, these observations indicate that COMET ZFa and promoters can be potentiated to the point where they begin to saturate the cellular capacity for transgene expression^23^, and there exist simple rules by which transcriptional output can be titrated up or down by selecting ZF binding site number, spacing, and compaction to tune promoter strength.

### Elucidating mechanisms of COMET gene expression

#### ZFa-induced gene expression is transcriptionally limited

When investigating how ZFa-responsive promoters respond to varying levels of TF, we observed that at high doses of the plasmid driving ZFa expression, reporter output plateaued at different levels depending on promoter architecture (**Fig. 2a, Supplementary Fig. 2a–c**). This plateau did not increase by switching to either a CMV or SV40 minimal promoter (**Supplementary Fig. 2d**). Although the CMV minimal promoter yielded higher gene expression at low ZFa doses, it did so at the cost of increased ZFa-independent background, which we deemed undesirable. To investigate whether the plateaus arose due to a bottleneck in transcription or translation, we increased the total plasmid dose while holding the ratio of the ZFa plasmid and reporter plasmid constant. For the ZF1×6-C promoter, reporter expression was linearly proportional to total plasmid dose (**Supplementary Fig. 2e**), which would not occur if translation were limiting, suggesting that in this system ZFa-driven reporter output is limited by the amount of plasmid template available for transcription.

**Fig. 2.**
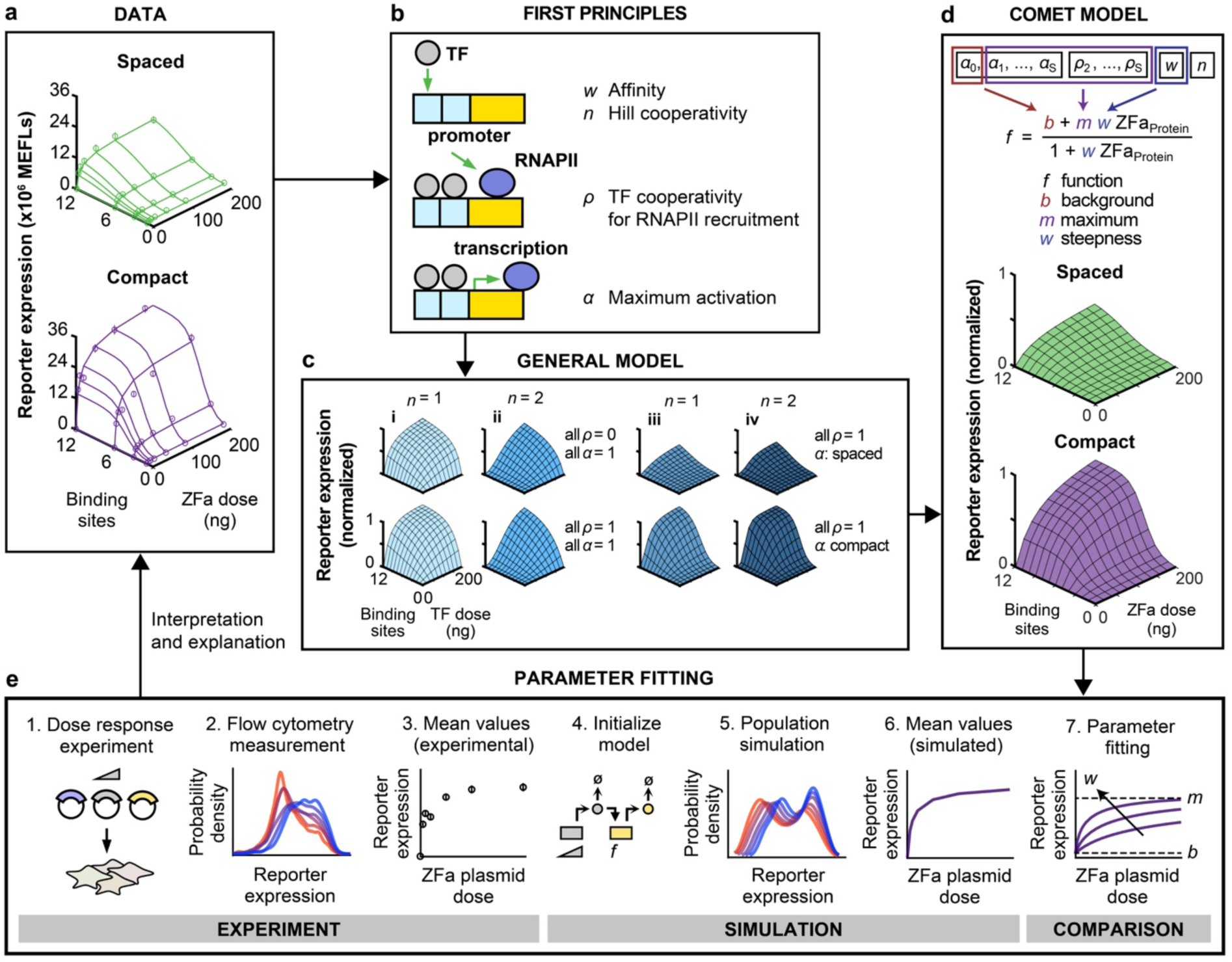
A model for COMET-mediated gene regulation. This figure summarizes the process of model development, refinement, and fitting. **a** The COMET model (model outputs are represented by the lines on each plot) explains experimentally observed trends (circles) for reporter expression as a function of ZFa dose and promoter features. This model uses a fitted transfer function for ZFa-induced gene expression (discussed in **b–e**) and simulates a cell population to account for variation in gene expression (**Supplementary Fig. 3**); lines depict the average outcome for the population. **b** We started with a detailed model of transcriptional activation in which reporter expression depends on TF concentration, a metric related to TF-DNA binding affinity (*w*), TF-DNA binding cooperativity (*n* = 1 for non-cooperative, *n* > 1 for cooperative), RNAPII recruitment cooperativity by each multiple-TF configuration at a promoter (*ρ* = 0 for non-cooperative, *ρ* > 0 for cooperative), and maximum promoter activation by each configuration (0 ≤ *α* ≤ 1). **c** This model yielded four types of landscapes (i–iv) under different assumptions, and two representative examples of each type are shown. COMET most closely resembles (iii). **d,e** A model that represents ZFa-induced reporter expression by a transfer function was used to fit the data in **a** (workflow for parameter estimation shown in **e**). The terms in this concise model can be related to terms in the mechanistic model. Landscapes in **c**,**d** are simulations of a single cell (homogenous model), and those in **a** are simulated mean values for a heterogeneous population. The outputs of this final fitted model are represented alongside experimental data in **a**.

#### COMET regulation can be explained by a concise mathematical model

To help elucidate the mechanisms by which COMET operates, we developed a mathematical model of this system. As summarized in **Figure 2** and detailed in **Online Methods**, our process involved first considering mechanistic steps involved in gene expression, writing equations that capture these steps (formally, writing a set of such equations is tantamount to formulating a hypothesis as to how gene expression operates), identifying a formulation that is consistent with experimentally observed behavior, simplifying this representation by removing details that are not required to describe the observed trends in order to generate a concise model, and finally fitting parameters of the concise model to data in order to quantitatively describe the experimental observations. We hypothesized that this process should generate a set of experimentally grounded parameters representing interpretable features of TF-promoter activity. Throughout, our goal was not to predict TF or promoter sequences *de novo*, but rather to describe and provide insight into observed trends. The explanatory value of such a model often exceeds insights that are accessible by intuition alone, and ultimately this framework could be used to design new genetic functions based upon COMET parts.

We initiated this process by using first principles to produce a detailed model with features of transcriptional control^24^ including physical and functional interactions between the promoter, TFs, and proteins like RNAPII (**Fig. 2b, Online Methods**). This detailed model relates transcriptional output to TF concentration, TF-DNA binding affinity, TF-DNA binding cooperativity, RNAPII recruitment cooperativity, and maximum promoter activation. We then generated a series of theoretical landscapes analogous to the experimental landscapes in **Fig. 2a,** varying parameters across a biologically reasonable range, and observed that the landscapes fell within one of four categories defined with respect to the concavity and sigmoidicity of cross-sections along each axis (**Fig. 2c**). The experimental data most closely resembled case (iii), indicating that TF-DNA binding is non-cooperative, but RNAPII recruitment is cooperative, and the maximum transcription rate (at a high ZFa dose) increases with both the number and compactness of binding sites.

Based upon the observed ZFa dose response profiles (**Fig. 2a)** and these insights, we proposed a concise transfer function to represent the rate of transcription (*f*) as a function of ZFa dose with three parameters: background (TF-independent) transcription (*b*), a steepness metric (*w*) related to TF-DNA-binding affinity, and a metric for maximum transcription (*m*) **(Fig. 2d, Online Methods**). As indicated, the three parameters in this concise transfer function can be related to the additional parameters in the original detailed representation. For a given ZFa-promoter combination, *m* is experimentally determined and is based upon the number and spacing of binding sites in the promoter, and *b* is determined based on reporter expression without ZFa; *w* can be fit to ZFa dose response data by our previously developed method that improves parameter estimation by accounting for variation in gene expression^25^ (**Fig. 2e, Supplementary Fig. 3a–c**; fitted parameters are listed in **Supplementary Tables 9–10**). Simulated data from the calibrated model provided close agreement with the experimental data, demonstrating that a concise representation can be used to analyze and describe COMET-mediated gene expression.

Comparison of the calibrated model and experimental data confirmed two trends that hold across conditions (**Supplementary Fig. 3d**). First, the dependence of relative reporter output on binding site number is independent of the dose of ZFa plasmid when the output is scaled to its maximum value in each binding site series. Second, the dependence of relative reporter output on ZFa dose is independent of the number of binding sites when the output is scaled to its maximum value in each dose series. Thus, inducible gene expression follows patterns that hold across various promoter designs and that are captured by a concise model.

### ZFa library characterization and orthogonality

#### Design rules are extendable to other ZFa

Building upon the characterization of the initial panel of five ZFa (**Fig. 1b**), we evaluated whether all nineteen previously characterized ZFa^15^ could activate gene expression in mammalian cells. We cloned the ZFa into mammalian expression vectors, constructed a new cognate x6-C promoter for each, and observed that all ZFa drove transcription from the promoters to varying extents (**Fig. 3a, Supplementary Fig. 4a**). The dose response profiles for the strongest twelve ZFa revealed a set of uncorrelated *m* and *w* values (**Supplementary Table 9, Supplementary Fig. 5a–c**). Additionally, the magnitude of induced reporter expression varied substantially between ZFa, which we hypothesized might be due to differential ZF affinity for binding cognate DNA sequences. One factor that can affect the strength of the ZF-DNA interaction is the nucleotide sequence between the canonical binding sites. In particular, the base pair upstream and the base pair downstream (“flanking nucleotides”) of each 9 bp binding site have been shown to affect ZF affinity^26^, and we did not account for these nucleotide preferences in the design of this initial panel of constructs. To investigate whether flanking nucleotide choices impact ZFa responsiveness, we revisited promoters for two ZFa with contrasting outcomes in **Fig. 3a** (ZF2a for high expression and ZF3a for low expression) and observed that the choice of flanking nucleotides significantly affected outcomes for both TFs (**Supplementary Fig. 5d**). In both cases, there existed choices guided by prior knowledge^15, 26^ that increased transcriptional activation by ZF2a or ZF3a. Therefore, although it seems that no specific flanking nucleotides are required for ZFa-mediated transcription, this choice can be used to tune transcriptional activation. To test whether the magnitude of reporter induction mediated by ZF2a and ZF3a depends on the number of binding sites in a manner similar to that observed for ZF1a (**Fig. 1e**), we varied the number of sites using compact promoters, and observed a similar trend for up to eight sites (**Fig. 3b**). Interestingly, there was a small decrease in reporter expression as the number of binding sites increased from 6 to 7 for both ZF2 and ZF3. It is possible that some promoter architectures, such as ZFx7-C, involve mechanisms that result in slight deviations from overall trends.

**Fig. 3.**
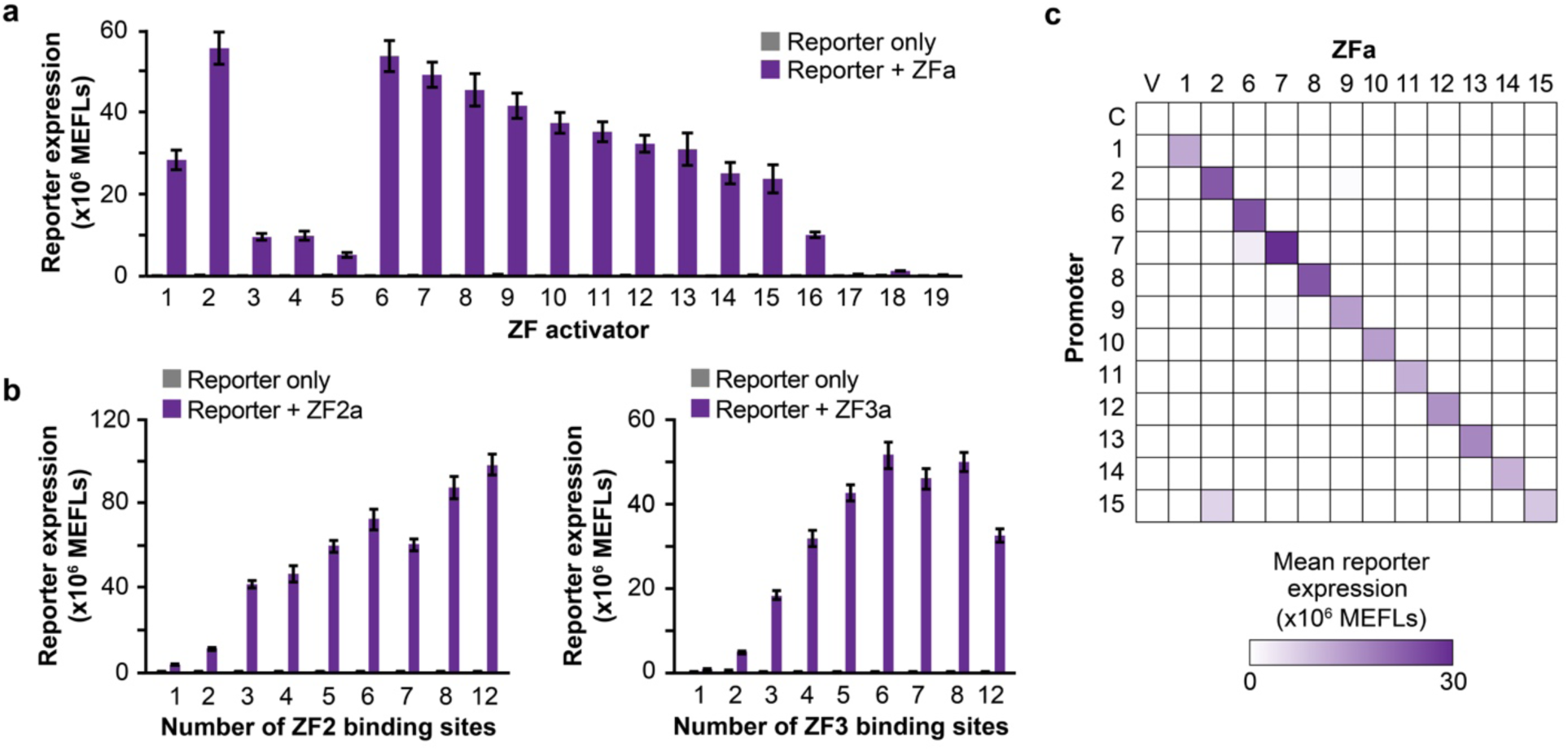
Characterizing an expanded panel of ZFa. **a** Nineteen ZFa were paired with cognate x6-C promoters, and all significantly induced gene expression (one-tailed Welch’s *t*-test all *p* < 0.02). **b** ZFa-induced gene expression increased with the number of binding sites, on compact promoters, for ZF2 (ANOVA *p* < 0.001) and ZF3 (ANOVA *p* < 0.001). **c** Investigating the orthogonality between the 12 strongest ZFa using x6-C promoters. Abbreviations: V (Vector control, no ZFa gene), C (Control reporter, no ZF binding sites). Experiments were conducted in biologic triplicate, and data were analyzed as described in **Online Methods**. Error bars represent the S.E.M.

#### ZFa are orthogonal

To test whether ZFa-mediated activation of cognate promoters is orthogonal across ZFa-promoter combinations, we performed a series of pairwise evaluations using the twelve strongest ZFa and x6-C reporters. The highest expression from each promoter was observed with its cognate ZFa (**Fig. 3c, Supplementary Fig. 5e**). Of the 132 pairs of ZFa and non-cognate promoters, 80% showed less than 1% of the maximal expression from that promoter (i.e., off-target activation), and 97% showed less than 3% off-target activation. The highest off-target activation of a ZFa/non-cognate promoter pair (ZF2a/ZF15×6-C at 75%) may be explained by the similarities in the binding site sequences for ZF2 and ZF15 (7 of 9 bp in common). However, such sequence similarities were not noted for the next three highest off-target combinations (ZF6a/ZF7×6-C at 10%, ZF7a/ZF15×6-C at 6%, and ZF7a/ZF9×6-C at 4% off-target activation). Overall, COMET ZFa are generally orthogonal from one another and are thus well-suited to composing genetic programs requiring multiple independent transcription units.

### Tuning transcription through protein engineering

#### Gene expression can be tuned through TF engineering

Having explored strategies for tuning gene expression by promoter engineering, we next investigated how COMET would be affected by two strategies for tuning via protein engineering: altering the affinity of the ZF for the DNA and altering the strength of the AD. For the first strategy, we mutated four arginine residues in the ZF that interact with the DNA backbone (**Fig. 4a**). Arginine-to-alanine substitutions in these positions ablate favorable charge interactions between the ZF and negatively-charged phosphates in the DNA backbone and were previously shown to decrease ZF1a-induced expression in yeast^15, 27, 28^. As expected, ZFa-mediated gene expression decreased with an increasing number of such substitutions (**Fig. 4b, Supplementary Fig. 6a,b**). Interestingly, while changing the promoter architecture affected only the maximum transcription (*m*) (**Fig. 2**), ZF mutations affected both the maximum transcription and the relative steepness of the ZFa dose response curve (*m* and *w*). Additionally, the changes in these values were correlated, revealing an axis along which ZFa R-to-A mutations tune TF strength. This result differs from our previous analysis of ZF domain choice, which affected *m* and *w* in an uncorrelated manner (**Supplementary Fig. 5c**). R-to-A mutations decreased ZFa-induced transcription in a manner that was similar across various numbers of binding sites in the promoter (**Supplementary Fig. 6c**), showing that this tuning can be applied across a variety of promoters. For the second tuning strategy, we tested two ADs in place of VP16: VP64^29^ and VPR^30^ (**Fig. 4c**). When fused in place of VP16, VPR produced the highest expression across several promoters, and VP64 was modestly stronger than VP16 in some cases (**Fig. 4d**). The relative effect of AD choice diminished as the number of ZF binding sites increased, suggesting that cooperative transcriptional activation by multiple weakly activating TFs (e.g., those based upon VP16), can approach the same magnitude of transcriptional activation mediated by fewer potently activating TFs (e.g., those based upon VPR). Replacing the AD on four other ZFa led to similar results (**Supplementary Fig. 6d**). Overall, these observations support the utility of multiple TF engineering strategies for tuning gene expression.

**Fig. 4.**
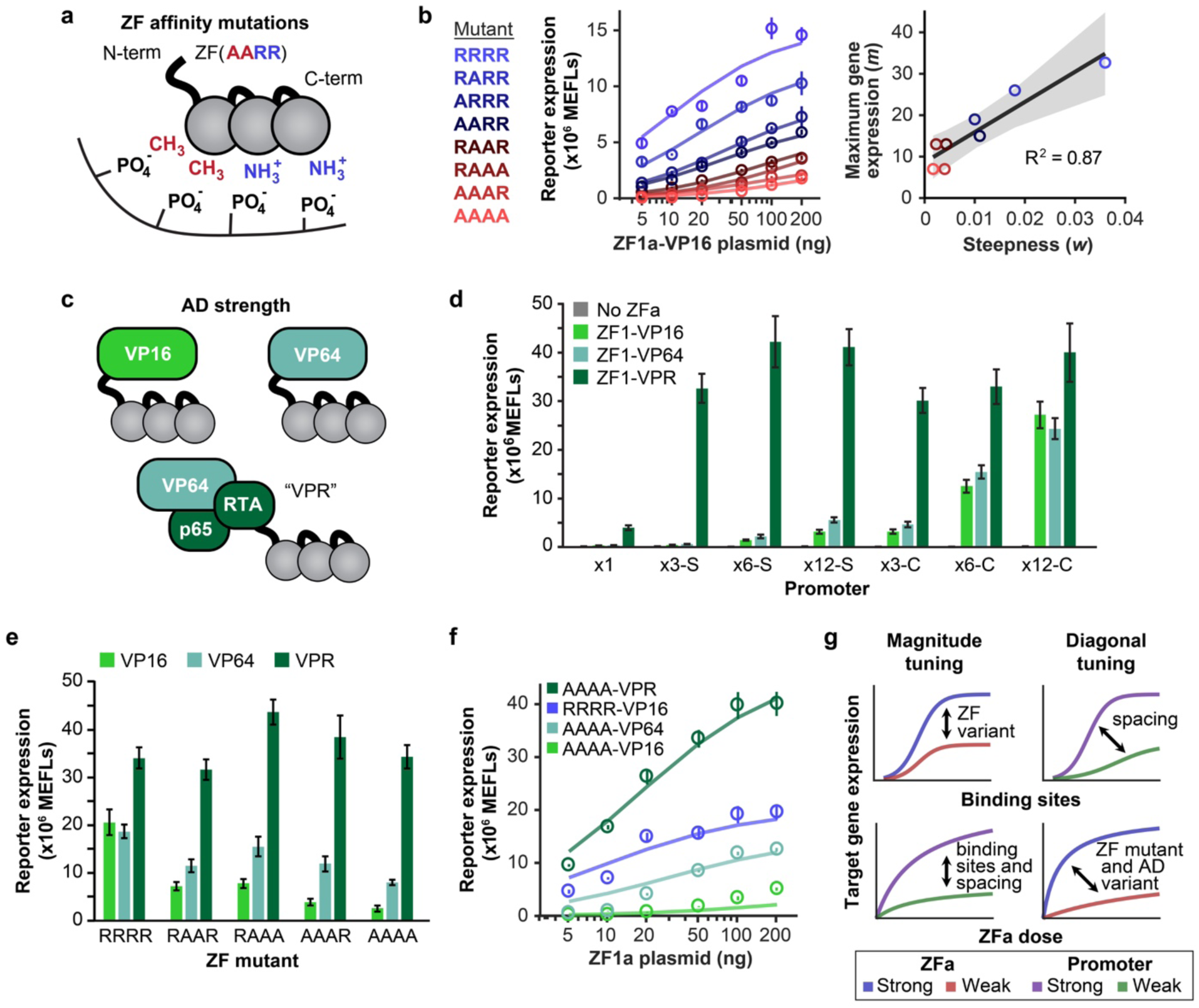
Tuning transcription through ZF mutants and AD variants. **a** The cartoon illustrates arginine-to-alanine (R-to-A) mutations in the ZF domain, which decrease the DNA-binding affinity. **b** *Left:* ZF mutations modulate the steepness and the maximum of the ZFa dose response profile. Circles represent experimental data and solid lines represent fitted transfer function models. *Right:* correlation between *m* and *w* parameters across mutants. The regression line is *m* = 7.3×10^2^*w* + 8.6, and the shaded region is the 95% confidence interval (one-tailed permutation test *p* < 0.001). **c** The cartoon depicts evaluated ADs. **d** Effects of AD on inducible reporter expression with different promoters. Gene expression varied with the choice of promoter (two-factor ANOVA *p* < 0.001) and choice of AD (*p* < 0.001), and an interaction was observed between these two variables (*p* < 0.001). **e** Combined effects of AD variants and ZF mutations were identified. Gene expression was affected by both the ZF mutations (two-factor ANOVA *p* < 0.001) and the AD (*p* < 0.001), with an interaction seen between these two variables (*p* < 0.001). Each mutant ZFa induced more reporter expression with VP64 than with VP16 (one-tailed Welch’s *t*-test, all *p* < 0.05) and with VPR than VP64 (one-tailed Welch’s *t*-test, all *p* < 0.01). All VPR-ZFa induced similar expression regardless of the use of a WT or mutant ZF (Tukey’s HSD test with α = 0.05). Error bars depict S.E.M. **f** The choice of AD affects the steepness and the maximum of the dose response. Circles represent experimental data and solid lines represent fitted transfer function models. Experiments were conducted in biologic triplicate, and data were analyzed as described in **Online Methods**. **g** The cartoon summarizes expected trends in output gene expression that result from tuning each modular feature of the ZFa and promoters. These design choices can produce either a vertical shift or diagonal shift in response profiles with respect to the number of binding sites and the dose of ZFa.

#### ZFa dose response profile can be tuned by TF engineering

To explore interactions between the two TF protein engineering strategies, we investigated whether stronger ADs could enhance gene expression conferred by TFs with low-affinity ZFs. As ZF binding affinity decreased, ZFa-mediated gene expression decreased substantially with VP16, yet only moderately with VP64 and not at all with VPR (**Fig. 4e**). We then compared the dose response for the weakest-binding ZFa mutant (AAAA) with each AD to the VP16 ZFa bearing a wild-type (WT) ZF domain (**Fig. 4f, Supplementary Fig. 6e**). As AD strength increased, both *m* and *w* increased, as was observed when varying DNA binding affinity. Although the two domains of a ZFa are physically modular, since they affect the same parameters in the transfer function, we find that the domains are functionally intertwined; i.e., these domain choices cannot be parametrically disentangled. Therefore, the two TF protein engineering strategies should be considered jointly when tuning a ZFa. In summary, our observations illustrate how gene expression can be tuned through selection of physical features—ZF domain choice, mutations that affect DNA binding affinity, AD choice, and the number, spacing, and arrangement of binding sites in the promoter—and together this ensemble of designs provides a variety of realizable response profiles (**Fig. 4g, Supplementary Fig. 6f**).

### Design of inhibitory TFs

#### ZFi effectively inhibit transcription

Inhibitors comprise a key component of a versatile TF toolkit. We hypothesized that removing the AD from the ZFa would result in an inhibitor that binds DNA without inducing transcription (ZF inhibitor, ZFi) (**Fig. 5a**). We built a promoter with six compact binding sites for ZF1 and in which each ZF1 site overlapped with a ZF2 site to allow for pairwise testing of ZFi and ZFa with fully or partially overlapping sites (**Supplementary Fig. 7a**). Co-expressing ZF1a with ZF1i or ZF2i (using equimolar amounts of plasmid) resulted in a ∼50% decrease in inducible expression compared to ZF1a only, and inhibition mediated by partially overlapping ZF2i resembled that mediated by fully overlapping ZF1i (**Fig. 5b, Supplementary Fig. 7b**). This pattern held across various ZFi doses, and nearly complete inhibition was attained at high ZFi doses (**Supplementary Fig. 8a**). We hypothesized that transcriptional inhibition could be increased by incorporating a bulky domain to sterically block ZFa from binding adjacent sites in the promoter or to block the recruitment of RNAPII or associated factors. To test this hypothesis, we fused DsRed-Express2 (abbreviated in this study as “DsRed”) to the ZF domain. Co-expression of ZFi-DsRed and ZFa (using equimolar amounts of plasmid) reduced reporter expression by 90–95%, and at higher ZFi-DsRed doses the inhibition was essentially complete, even when using stronger promoters based upon the CMV minimal promoter (**Fig. 5b, Supplementary Figs. 8b-c**).

**Fig. 5.**
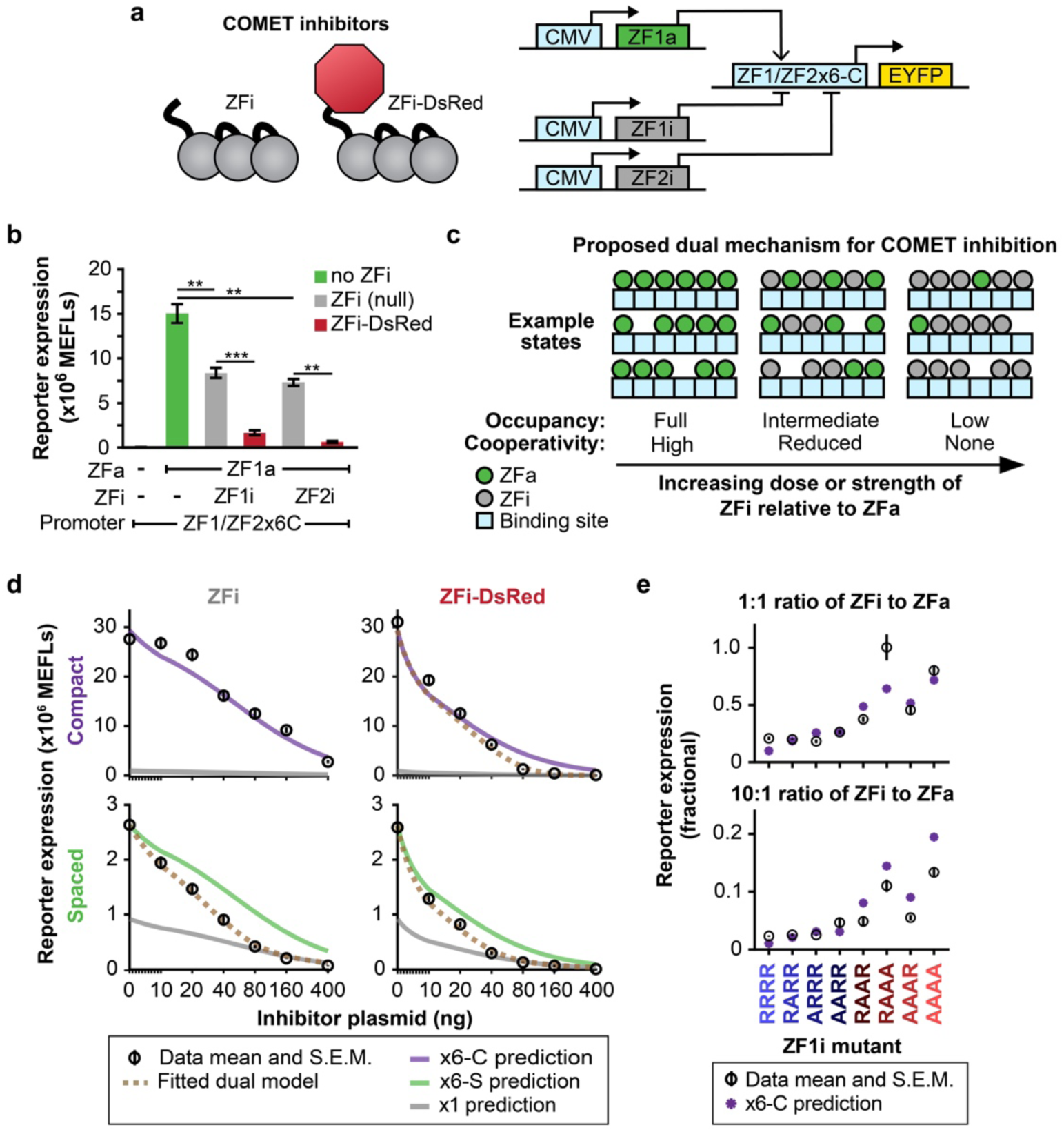
Transcriptional inhibition. **a** The schematic depicts two types of inhibitors that were evaluated. A ZF1/ZF2×6-C hybrid promoter is activated by ZF1a and inhibited by ZF1i or ZF2i. **b** ZFi and ZFi-DsRed differentially inhibit transcription (one-tailed Welch’s *t*-test: \*\**p* < 0.01, \*\*\**p* < 0.001). **c** The cartoon summarizes the proposed conceptual model of ZFi-mediated inhibition. Within each cell, a promoter can occupy states with different configurations of ZFa and ZFi. Several example states are shown for three conditions of increasing dose or strength of inhibitor (i.e., DNA-binding affinity) relative to activator. **d** ZFi and ZFi-DsRed differ from standard competitive inhibitors. Predictions for competitive inhibition alone, for various promoter configurations, are depicted with solid lines (**Online Methods**). COMET inhibitors track the dotted lines, which represent fits to the dual mechanism model, except in the case of ZFi paired with x6-C, which tracks the competitive inhibition-only prediction. Each condition uses ZF1a at a dose of 40 ng. X-axes are scaled linearly from 0–10 ng and logarithmically above 10 ng. **e** Measured and predicted reporter expression were compared for a panel of ZFi mutants. Each condition uses ZF1a(RAAR) at a dose of 40 ng and the ZF1×6-C compact promoter. Experiments were conducted in biologic triplicate, and data were analyzed as described in **Online Methods**. Error bars represent the S.E.M.

#### COMET inhibitors use a dual mechanism

To help understand the mechanism of ZFi-mediated transcriptional inhibition, we considered that within each cell, promoters occupy an ensemble of states that depend on the promoter architecture and the ZFa and ZFi that are present (**Fig. 5c**). As the relative dose of ZFi to ZFa increases, the distribution of the ensemble should shift toward states that are more inhibited; a trend towards more inhibition should also occur by increasing the relative DNA binding affinity of the ZFi versus that of the ZFa. Given our understanding of ZFa-mediated transcriptional activation, we speculated that the inhibitors should act via a *dual mechanism* with these properties: (i) competitive inhibition: since each site in the promoter can accommodate at most one TF, the binding of an inhibitor should prevent the binding of an activator; and (ii) decreased cooperativity: since inhibitors intersperse between activators, the spacing between activators should widen, and the *effective m* should resemble that of a promoter with lower cooperativity.

To explore this proposed mechanism of inhibition, we developed a model that describes the activity of ZFa and ZFi at a single-site promoter by representing physical interactions without a transfer function (**Supplementary Fig. 8d, Online Methods**). Simulated trends for ZFa dose responses with perturbations to DNA-binding affinity broadly agreed with experimental data (**Supplementary Fig. 8e, Fig. 4**). However, simulated trends for ZFa dose responses for varying AD strengths (at the simulated *single-site* promoter) differed qualitatively from the trends observed experimentally for a *multi-site* promoter (**Fig. 4f**). The difference in outcomes for the single-site and multi-site cases is consistent with our expectation that cooperative ZFa-mediated RNAPII recruitment would be observed only for the latter case (**Fig. 2**). Notably, the model also showed less responsiveness of reporter output to ZFi (at the simulated single-site promoter) than was experimentally observed for a multi-site promoter (**Supplementary Fig. 8f**), again suggesting that for multi-site promoters, ZFi may impair ZFa-mediated transcription by disrupting cooperative RNAPII recruitment.

To experimentally test the proposed dual mechanism of inhibition, we conducted dose responses for the ZFi and ZFi-DsRed inhibitors using the ZF1×6-S and ZF1×6-C promoters, with ZFa dose held constant (**Fig. 5d**). When ZFi was applied to the compact promoter, reporter expression matched the concise model for competitive inhibition alone. However, for the other three cases, observed reporter expression began to deviate with increasing doses of inhibitor, and by high doses it showed complete loss of cooperative RNAPII recruitment. The inhibitor dose at which the experiment began to deviate from the model was lower for ZFi-DsRed compared to ZFi and for spaced promoters compared to compact promoters. At intermediate doses of inhibitor, reporter expression ramped down toward single-site promoter behavior (**Fig. 5c** middle column, **Fig. 5d** dotted lines, **Online Methods**), and by high doses the ramp down was complete (**Fig. 5c** right column). The highest dose of ZFi-DsRed, used with the compact promoter, resulted in a profound 400-fold decrease in reporter expression. To further examine the case where the employed inhibitor did not disrupt cooperative RNAPII recruitment (i.e., ZFi used with the x6-C promoter), we paired a panel of ZFi varying in DNA-binding affinity with a reduced-affinity ZFa mutant (**Fig. 5e**). For all cases examined, ZFi-mediated inhibition was still predicted by competitive inhibition alone (**Online Methods**). We conclude that the compact promoter is more capable of cooperative RNAPII recruitment than is the spaced promoter, and that ZFi is a weaker inhibitor than is ZFi-DsRed, such that the dual inhibition mechanism applies to three of the four types of inhibitor-promoter pairings evaluated, and the pairing most responsive to inhibition is ZFi-DsRed with a compact promoter.

### Genomic integration of COMET TFs

#### COMET design rules extend to the genomic context

Since some applications require stable integration of genetic programs, we investigated how COMET parts function upon stable integration into the genome, and in particular, whether COMET design rules gleaned from transient transfections might extend to performance in the genomic context. As a representative test set, we generated a panel of stable cell lines that each constitutively express a ZFa and contain one of several COMET promoters—varying the number of ZF binding sites and spacing between binding sites—that drive expression of a fluorescent reporter protein. To enable comparisons using a consistent site of genomic integration, we used site-specific BxB1 recombinase-mediated integration into the AAVS1 safe harbor locus of HEK293FT landing pad cells.^31^ In this process, COMET components were cloned into transcription unit positioning vectors (TUPVs) followed by one-step assembly into all-in-one integration vectors (IVs). The IVs used include a constitutive fluorescent protein marker and antibiotic resistance markers, a COMET promoter-driven mKate2 reporter, and either a constitutively expressed VP16-ZF1a or a blank control sequence (**Supplementary Fig. 9a–c**). Following lipofection of cells with these vectors, drug selection, and FACS recovery based on the constitutive fluorescent marker (**Supplementary Fig. 9d**), we obtained cell lines that enable a comparison of COMET-driven gene expression in the stable genomic context (**Fig. 6c, Supplementary Fig. 10c**) to trends for the delivery of the same parts by transfection of separate plasmids (**Fig. 6a, Supplementary Fig. 10a**) or transfection of all-in-one vectors (**Fig. 6b, Supplementary Fig. 10b**).

**Fig. 6.**
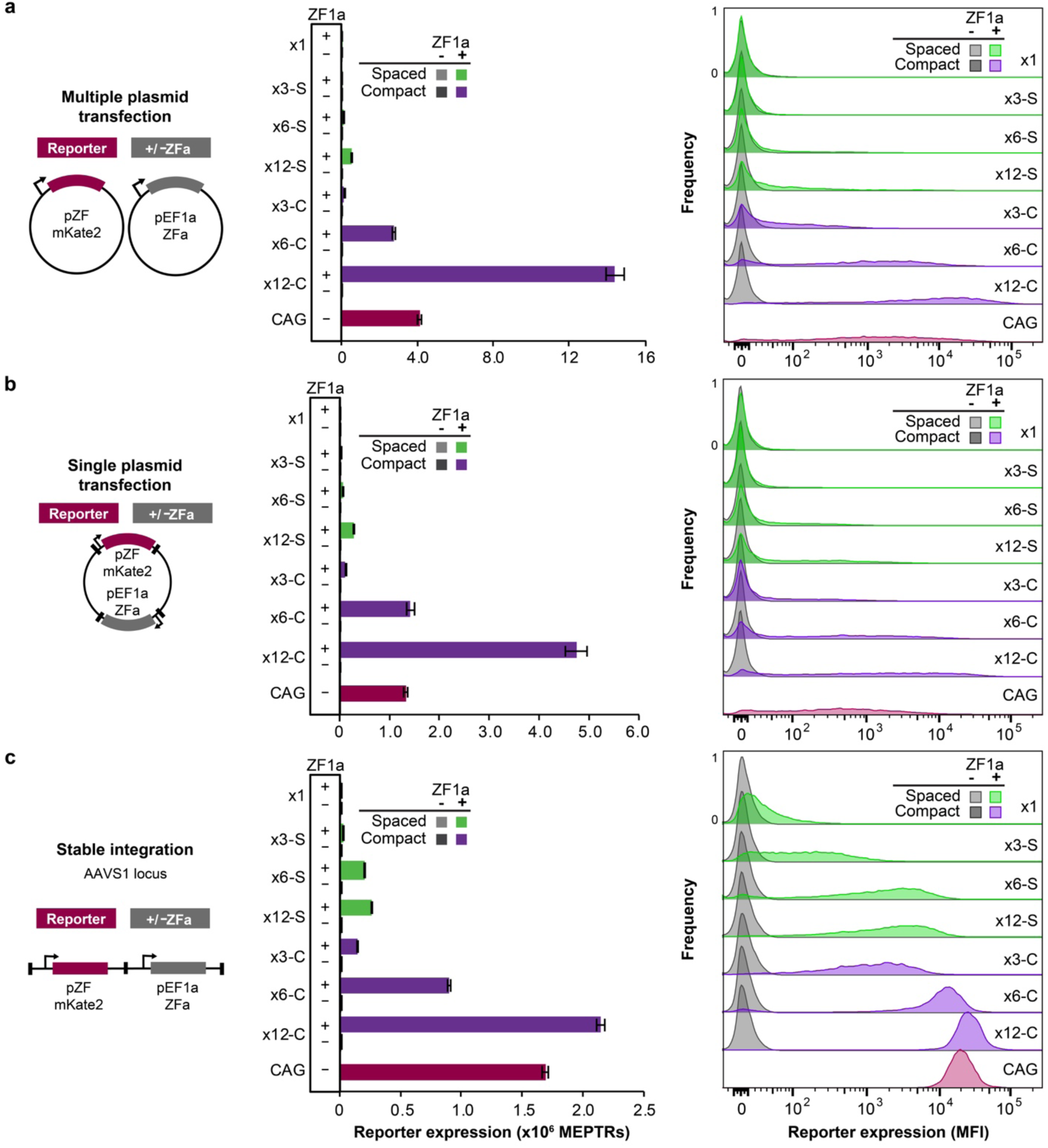
Characterization of promoter design rules in the genome. The cartoons summarize the systems used to evaluate promoter performance characteristics across three contexts: **a** multiple plasmid transient transfection, **b** single plasmid transient transfection, and **c** single-copy stable integration at a genomic safe harbor locus. The promoters included here comprise 1, 3, 6, or 12 ZF1 binding sites positioned using spaced or compact architectures upstream of the YB_TATA minimal promoter driving an mKate2 reporter gene. Constitutive EBFP2 was used as a transfection control in the transient transfection context and as a marker for genomic locus activity in the stable context. Bar graphs and histograms show reporter expression for EBFP2-expressing cells. In all contexts, ZFa-induced gene expressed increased with the number of binding sites on spaced and compact promoters (ANOVA *p* < 0.00001). To profile the range of inducible expression conferred by each promoter, stable cell lines and transiently transfected cells were characterized using two distinct sets of flow cytometry settings (voltages), each of which was independently calibrated to yield comparable absolute fluorescence units (bar graphs). Experiments were conducted in biologic triplicate, and data were analyzed as described in **Online Methods**. Error bars represent the S.E.M.

Overall, genomically integrated ZFa and COMET promoters drove gene expression following trends that are consistent with those observed in transient transfection. Compact promoters drove more expression than did spaced promoters for each number of binding sites tested. For each architecture, reporter expression increased with the number of binding sites. Interestingly, for compact promoters, increasing the number of binding sites also led to more homogeneous reporter expression profiles spanning only a single order of magnitude—matching the tight distribution expected of a constitutively expressed gene in a landing pad.^31^ For the strongest promoters (x6-C and x12-C), tight distributions of reporter expression contributed to high fold inductions (8,000 and 14,000, respectively, compared to corresponding reporter-only cell lines). The promoter containing a single ZF1 site, placed in a favorable position with respect to the TATA box (**Fig. 1d**), did confer modest but significant gene expression compared to the control promoter lacking any ZF1 site (**Supplementary Fig. 10d**), although the expression induced by this ZFa from a x12-C promoter was 800-fold higher (**Fig. 6c**). Together, these observations indicate that COMET TFs can drive expression from a cognate promoter that is either genomically integrated or transfected, and that the design rules used to tune expression in transient transfections may be transferrable, at least qualitatively, to the genomic context.

### Design and evaluation of small molecule-responsive TFs

#### Rapamycin-activated ZFa (RaZFa) control gene expression

Chemical inducibility is a useful strategy for conferring external and temporal control over gene expression. We designed a small molecule-responsive ZFa by fusing FBKP and FRB domains, which heterodimerize upon exposure to rapamycin^32^, onto ZF and AD, respectively (**Fig. 7a**). We expected that without rapamycin, the ZF would bind DNA and not induce transcription, and that with rapamycin, FKBP and FRB would dimerize to reconstitute a functional ZFa. Indeed, when RaZFa with each of the three ADs were tested with ZF1×6-S and ZF1×6-C promoters, all cases showed rapamycin-induced reporter expression (**Fig. 7b, Supplementary Fig. 11a**).

**Fig. 7.**
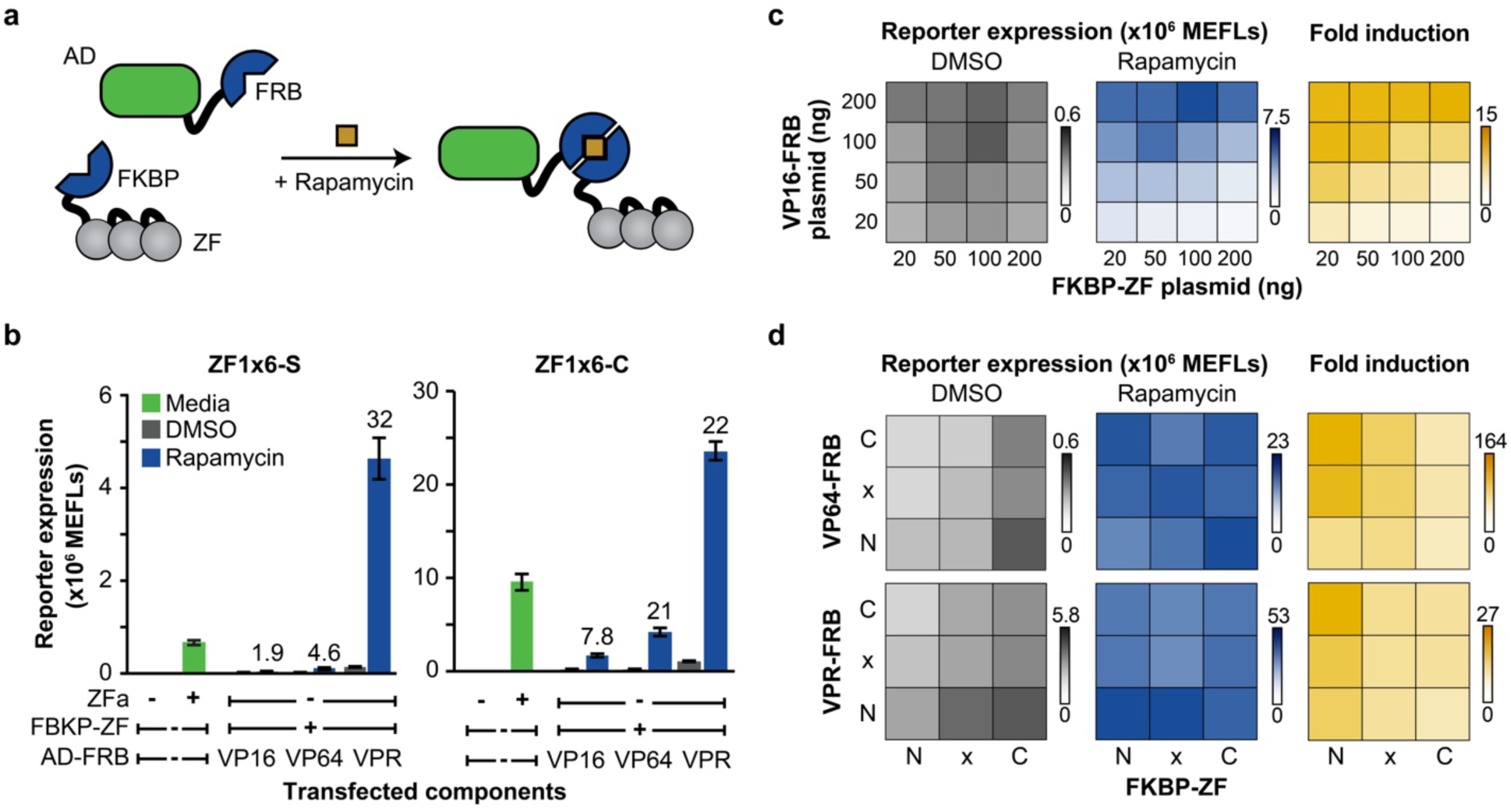
Engineering small molecule-responsive TFs. **a** The cartoon illustrates chemically-responsive control of gene expression using rapamycin-inducible ZFa (RaZFa). **b** The effects of promoter architecture and AD on RaZFa performance were evaluated. For all RaZFa on both promoters, reporter expression was significantly higher with rapamycin than DMSO (one-tailed Welch’s *t-*test, all *p* < 0.05). Fold induction is shown above the rapamycin case for relevant conditions. **c** Gene expression in the absence of rapamycin was affected by VP16-FRB dose (two-factor ANOVA *p* < 0.001) and FKBP-ZF dose (*p* < 0.001), with no interaction between these variables (*p* = 0.14). Reporter expression after rapamycin addition was affected by VP16-FRB dose (two-factor ANOVA *p* < 0.001) and FKBP-ZF dose (*p* < 0.001) with a significant interaction between these variables (*p* < 0.001). **d** Effects of subcellular localization tags: N = nuclear, x = no localization, C = cytoplasmic. For VP64-based RaZFa, gene expression in the absence of rapamycin was affected by AD-FRB localization (two-factor ANOVA *p* = 0.01) and FKBP-ZF localization (*p* < 0.001), with no interaction between these variables (*p* = 0.39). For VP64-based RaZFa, gene expression after rapamycin addition was not affected by AD-FRB localization (two-factor ANOVA *p* = 0.26) but was affected by FKBP-ZF localization (*p* = 0.02), with an interaction (*p* = 0.001). For VPR-based RaZFa, gene expression in the absence of rapamycin was affected by AD-FRB localization (two-factor ANOVA *p* < 0.001) and FKBP-ZF localization (*p* < 0.001), with an interaction (*p* = 0.03). For VPR-based RaZFa, gene expression in the presence of rapamycin was affected by AD-FRB localization (two-factor ANOVA *p* < 0.001) but not by FKBP-ZF localization (*p* = 0.29), with no interaction (*p* > 0.05). Experiments in **c** and **d** use a ZF1×6-C promoter. Experiments were conducted in biologic triplicate, and data were analyzed as described in **Online Methods**. Error bars represent the S.E.M.

#### RaZFa component ratios can impact performance

We observed that for each AD tested in **Fig. 7b**, fold-increase in reporter output was lower for the RaZFa (+/– rapamycin) than for the ZFa (+/– ZFa). For the RaZFa using VP16, this effect appeared to be attributable to low induced reporter expression. We hypothesized that if FKBP-ZF were present in excess, it might competitively inhibit the reconstituted RaZFa from binding the promoter. To investigate this possibility, we varied the doses and ratios of RaZFa components (**Fig. 7c, Supplementary Fig. 12a**). High FKBP-ZF levels diminished the expression as a ZFi would, and excess VP16-FRB increased the inducible expression, resulting in high fold induction when paired with lower doses of FKBP-ZF. However, VP64-based RaZFa and VPR-based RaZFa were less affected by component ratios (**Supplementary Fig. 12b,c**). Thus, it appears that the relative weakness of VP16-mediated transcriptional activation makes VP16-based RaZFa more sensitive to excess FKBP-ZF.

#### Altering RaZFa subcellular localization decreases ligand-independent reporter expression

Since high background in the absence of rapamycin limited the fold induction for VP64-based and VPR-based RaZFa, we evaluated strategies to decrease the background. First, to determine the source of the background, we transfected VPR-FRB with various ZF-fusions. VPR-FRB alone, in the absence of any exogenous protein with a DNA-binding domain, promoted a very low amount of reporter expression, and this background was greater in the presence of ZF-fusion proteins, even in the absence of rapamycin (**Supplementary Fig. 12d**), suggesting that the ZF may bind the promoter in such a way that transient promoter-AD interactions induce some transcription. To circumvent this putative undesired mechanism, we removed the nuclear localization signal (NLS) from each RaZFa component or replaced the NLS with a nuclear export signal (NES), and evaluated these variants pairwise (**Fig. 7d, Supplementary Fig. 12e**). For both VP64-based and VPR-based RaZFa, NES tagging of AD-FRB and NLS tagging of FKBP-ZF decreased the background. These modifications had little effect on rapamycin-induced reporter expression, such that the fold induction in reporter expression with rapamycin treatment improved compared to when both components were tagged for nuclear import.

The decrease in background reporter expression following the addition of NES to AD-FRB was expected, but the increase in background following the addition of NES to FKBP-ZF was not. To explain these observations, we hypothesize that while low levels of nuclear FKBP-ZF are sufficient to allow AD-FRB to drive transcription from the promoter, at higher nuclear levels the FKBP-ZF may act as an inhibitor, with the FKBP domain sterically occluding access to the promoter by AD-FRB. Furthermore, the decrease in background associated with the NES tag on AD-FRB was not due to decreased component expression; western blot analysis showed that adding the NES *increased* VP64-FRB and VPR-FRB expression (**Supplementary Fig. 12f**). We also observed that the expression of VP64-FRB was low relative to other components, but a functional assay in which the dose of VP64-FRB plasmid was increased to levels much higher than those used in **Supplementary Fig. 12b** resulted in increased background and diminishment of inducible reporter expression (**Supplementary Fig. 12g**). Thus, the relatively low expression of VP64-FRB is not limiting for RaZFa-driven reporter expression.

This optimization across component ratios and subcellular localization led to improved rapamycin-inducible gene expression (greater fold induction) for RaZFa with each of the three ADs (**Supplementary Fig. 13a**), and these TFs were responsive across several orders of magnitude of rapamycin concentration (**Supplementary Fig. 13b**). In summary, COMET TFs are amenable to external control by small molecule-induced reconstitution, with features that are tunable through the choice of promoter and AD.

### Implementing Boolean logic with COMET

#### Hybrid promoters enable AND logic

Finally, we explored whether COMET could be used to encode Boolean logic functions within individual promoters. Our characterization of reporter output as a function of ZF binding site number and spacing (**Fig. 1c,e**) suggested a strategy for designing hybrid promoters comprised of alternating sites that could be activated by combinations of ZFa to implement AND logic (**Fig. 8a**). We tested promoters containing one to four pairs of sites for ZF2 and ZF3 (**Fig. 8b, Supplementary Fig. 14a**). In each case, the highest reporter expression occurred only when both ZFa were present, and this expression was greater than the sum of those induced by each ZFa individually, demonstrating AND gate behavior. In surveying the dose response landscape of the three-pair hybrid promoter, AND gate behavior was observed even at low ZFa levels; 5 ng of each plasmid encoding ZF2a and ZF3a together produced more reporter expression than did 200 ng of either plasmid encoding either ZFa alone (**Fig. 8c, Supplementary Fig. 14b**). The steep OFF-ON transition along the entire perimeter of the landscape is due to the effective transition between x3-S and x6-C architectures—an advantageous behavior of COMET that differs from previously reported transcriptional AND gates such as those utilizing tTA and Gal4 (**Fig. 8d, Supplementary Fig. 14c, Online Methods**)^25^.

**Fig. 8.**
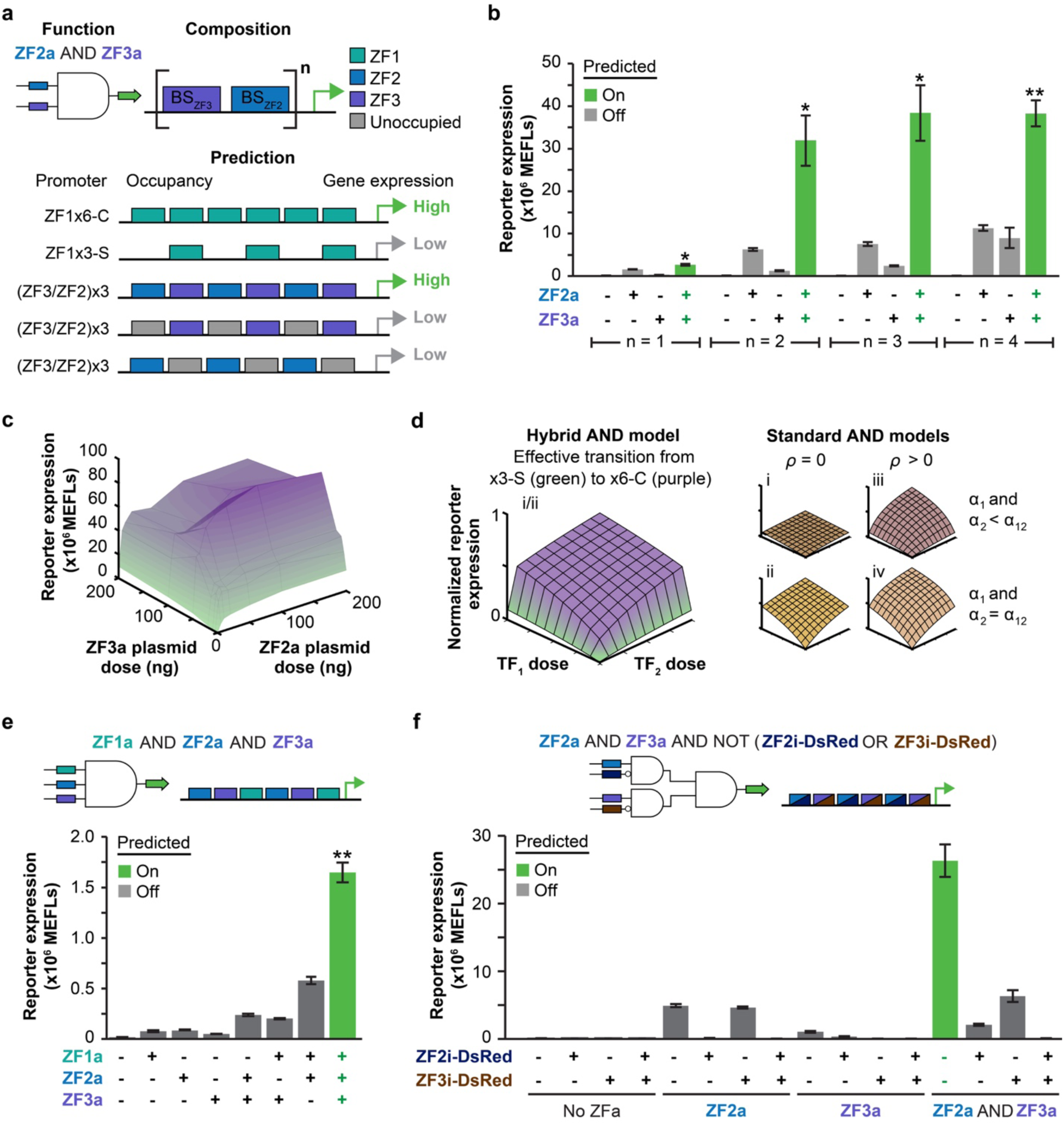
Composing Boolean logic. **a** The cartoon summarizes a strategy for single-layer, promoter-based logic gates with ZF-TFs. We hypothesized that AND gate promoters could be designed by using multiple repeats of a paired ZF3/ZF2 motif. Full occupancy of this promoter by both ZF2a and ZF3a mimics a fully occupied x6-C promoter, and partial occupancy (with either ZFa alone) mimics an x3-S promoter. Thus, there is a large increase in gene expression when the promoter is occupied by two types of ZFa compared to one type. **b** Candidate two-input AND gates were constructed using one to four repeats of paired binding sites in the promoter. AND gate behavior is considered significant if reporter expression with both ZFa is greater than the sum of reporter expression with each ZFa individually (one-tailed Welch’s *t*-test: \**p* < 0.05, \*\**p* < 0.01). **c** Two-input dose response for the AND gate with three repeats of paired binding sites. **d** A theoretical model of COMET AND behavior is compared with other models of transcriptional AND gates; the latter vary in whether activators have multiplicative cooperativity (*ρ*) and whether maximum activation (*α*) is equivalent for TFs individually and together (**Online Methods**). **e** A three-input AND gate was constructed using two repeats of a triplet binding site motif. AND gate behavior is considered significant if reporter expression with all three ZFa is greater than the sum of reporter expression with each ZFa individually, and also greater than the sum from each of the three combinations with two co-expressed ZFa and the other ZFa individually (one-tailed Welch’s *t-*test, \*\**p* < 0.01 for all four of these tests). **f** A four-input gate was constructed using the binding site arrangement shown. Experiments were conducted in biologic triplicate, and data were analyzed as described in **Online Methods**. Error bars represent the S.E.M.

We extended this hybrid promoter strategy to generate candidate three-input AND gates for ZF1a, ZF2a, and ZF3a. A promoter with one site for each ZFa did not produce AND gate behavior (**Supplementary Fig. 14d**). This outcome is consistent with the expected similarity in reporter expression for x2-S/C versus x3-C promoter architectures, corresponding to cases when two or three ZFa are present, respectively. However, a promoter with two sites per ZFa did produce AND gate behavior; reporter expression when all three ZFa were present was higher than the sum of the levels when any two ZFa were present plus the level conferred by the third (**Fig. 8e**). Thus, COMET’s modular features innately enable hybrid promoters that utilize cooperative transactivation to compose two-input and three-input AND gates.

#### Complex logic functions can be composed by combining ZFa and ZFi

Finally, we investigated whether inhibitors could be combined with activators to build complex logic functions using design rules elucidated in this study. As a test case, we designed a four-input logic function that takes both activators and inhibitors as inputs (**Fig. 8f, Online Methods**). Before testing the combined circuit, we characterized individual interactions between activators and inhibitors. We tested how ZF1i-DsRed, ZF2i-DsRed, and ZF3i-DsRed individually inhibited gene expression from an AND gate promoter (with sites for ZF3 and ZF2, with overlap for ZF1) when ZF2a and ZF3a were both present and found that ZF2i-DsRed and ZF3i-DsRed were the most effective at inhibiting expression (**Supplementary Fig. 14e,f**). Testing the 16 combinations of ZF2a, ZF2i-DsRed, ZF3a, and ZF3i-DsRed showed that all cases produced the expected outcomes (**Fig. 8f, Online Methods**)—expression was highest with both ZFa and no ZFi-DsRed, low with only one ZFa or with only one ZFi-DsRed, and off in the remaining cases. Thus, COMET components and design principles can be employed to compose complex functions including single-layer logic.

## DISCUSSION

We anticipate that the COMET toolkit—an ensemble of TFs and promoters for controlling gene expression in mammalian cells—will be a useful resource for building genetic programs. Currently, the engineering of mammalian cellular functions is often a slow process involving multiple incremental iterations of the design-built-test-learn cycle. However, in prokaryotes, the design and construction of genetic programs has been streamlined by the development of large libraries of well-characterized and orthogonal components in concert with computational tools such as Cello^33^. COMET similarly provides a large library of TFs and promoters with tunable features, and the characterization of these components provided a foundation for a mathematical model. We used the model to elucidate mechanisms by which the activators and inhibitors operate at promoters and fitted parameters to describe how these activities vary across the design choices examined in this study. This integrated approach transcends the identification of general qualitative trends (e.g., increasing the number of binding sites in a promoter generally increases inducible gene expression) to yield quantitative and often mechanistic understanding as to how design choices affect TF-promoter activity. This insight could not have been deduced from prior knowledge, including biophysical intuition or even characterization of similar ZFa and promoters in yeast^15^. Whether the design rules elucidated here ultimately enable large-scale model-driven design is an important question worthy of subsequent investigation.

COMET comprises a framework that readily accommodates new parts. The current COMET library includes 44 activating and 12 inhibitory TFs and 83 cognate promoters. Of the 44 ZFa, 19 were ported from a toolkit originally characterized in yeast,^15^ with only minor changes in the linker between protein domains. The remaining activators and inhibitors represent design iterations implemented by combining the ZF domains with functional domains. This highlights the modularity of the COMET toolkit, in that functional domains, ZF domains, and promoters can be characterized and assigned parameters (**Fig. 2, Online Methods**) and then utilized for customized gene regulatory functions. Thus, COMET not only comprises the components described in this work but also serves as an extensible toolkit.

Our combined experimental and computational investigation revealed properties and design rules that guide the use of COMET parts. By selecting TF-promoter combinations, one can select a magnitude of output gene expression from a range spanning three orders of magnitude. The design rules explain, at a high level, many functional consequences of choices such as ZF domain, mutations in the ZF domain that impact binding affinity, the AD, competition between activating and inhibitory TFs, and the number, spacing, and arrangement of binding sites in the promoter. These characterizations also revealed that COMET-mediated gene expression confers dose response landscapes that differ from those of other systems such as tTA and Gal4^25^, and as consequence, COMET may be better suited for applications such as building hybrid promoters. The COMET system is also amenable to incorporation of other functional modalities such as using rapamycin-induced dimerization domains to confer chemically inducible gene expression, and the use of other dimerizing domains is certainly possible.

A key insight from this study is that COMET promoter strength arises from cooperative recruitment of transcriptional machinery, which is an effect that varies with the spacing between binding sites. This mechanism differs from that of previously characterized ZF-TF systems in which cooperativity is directly engineered into TFs through protein-protein interaction domains such as PDZ or leucine zippers^9, 15, 34^. While these previous strategies usefully enable tuning performance characteristics such as dose response curves, they are potentially limited by the availability, orthogonality (with respect to both synthetic and endogenous components), and geometric requirements of the protein-protein interaction domains employed. In contrast, the scalability of COMET thus far appears limited only by the availability of orthogonal ZFs; these domains can be constructed using technologies such as OPEN^16^ as well as other methods, and this remains an active area of research.

COMET’s promoter design-based cooperativity confers several useful properties. First, it allows one to achieve both low ZFa-independent (background) expression and high fold induction, even though these two objectives typically present a trade-off^20^. For example, pairing an x6-C ZF binding site array with the “weak” YB_TATA minimal promoter yielded such desirable performance, whereas achieving a high ON state with TFs lacking cooperative transcriptional activation may require using a stronger minimal promoter (such as CMV minimal promoter) at the cost of elevated background (**Supplementary Fig. 2d**).

Promoter design-based cooperativity also enabled the implementation of logic gates, which have several advantageous features. Unlike other previously described logic gates for mammalian cells that require different architectures for activation and inhibition^9^, a single COMET promoter can serve as the template for both activating and inhibitory logic gates, which can be constructed without extensive tuning (**Fig. 8**). This simplifies the design process, enables the reuse of previously characterized parts, and increases the number of inputs that can be integrated at a single promoter, which may ultimately decrease the number of components required to construct genetic circuits. Inhibitory COMET TFs that function via modulation of effective cooperativity, rather than KRAB-mediated recruitment of chromatin regulators, confer the benefits of speed and reversibility, which may be advantageous given the relatively slow timescales of KRAB-mediated chromatin repression and subsequent reactivation^35^. Even without KRAB, the ZFi-DsRed are sufficiently potent to completely inhibit COMET TF-mediated transcriptional activation (**Fig. 5, Supplementary Fig. 8**).

Another advantage of promoter-based cooperativity is that it should enhance the specificity with which ZFa activate target promoters. A limitation to the minimal three-finger ZF-TF strategy investigated here is that any single 9 bp ZF binding sequence may occur many times in a genome. However, the probability that two binding sites would occur at the same locus is unlikely, and the chance that three or more sites would co-occur is vanishingly small. Moreover, the potent activation reported in **Fig. 1** also required the ZF binding array to be proximal to a transcriptional start site, which should further boost the distinction between on-target and off-target transcription. Indeed, in a genomic context (**Fig. 6**), although ZF1a drove modest expression from a x1 promoter (in which the ZF binding site was placed favorably close to the TATA box), the expression from a x12-C promoter was 800-fold greater. The protein engineering design rules elucidated here also suggest that specificity could be further increased, if desired, by the choice of AD and ZF domain. For example, selection of a weaker AD could necessitate that multiple ZFs bind in a compact configuration at a promoter in order to drive transcription (**Fig. 4d**). Reducing the affinity with which a ZF binds DNA could also be combined with selection of a weaker AD to shift the dose response curve, such that a promoter is activated only at high concentrations of ZFa (**Fig. 4f**). Thus, a potential advantage of pairing weaker ZFa with multi-site promoters is the possibility of dramatically boosting the effective specificity of the ZFa for driving transcription from a target promoter. Since chromatin state, and thus cell type, likely impacts the tradeoff between on-target and off-target gene regulation, we suggest that this question is worthy of exploration in the future use of COMET for specific applications.

Our study also revealed several properties of COMET that are not easily explained by simple design rules. It is not yet clear why some ZFa combinations exhibit limited crosstalk when no sequence similarity in ZF binding sites is apparent (**Fig. 3c**); our empirical evaluation simply identifies how such crosstalk can be avoided by careful selection. For example, interestingly, some non-specific activation of gene expression at COMET promoters was conferred by potent ADs (e.g., VPR) when ZF domains were separately expressed but not driven to physically associate (i.e., by addition of rapamycin), suggesting some potential for interactions between strong ADs and promoters whose states are regulated by distinct proteins. Operationally, these phenomena present minor complications that can be circumvented by careful system selection and attentiveness to potential artifacts during the development and design of new functions.

Going forward, an important investigation will be to evaluate how the trends observed here are conserved or diverge as the COMET toolkit grows and is applied to new applications. The fundamental modularity of the components renders it straightforward to incorporate additional TFs and cognate promoters. Currently, the magnitude of gene expression that a new ZF domain will confer in a ZFa and with a cognate promoter is not known *a priori*. However, our analyses suggest that such a new part will confer some function, and that the magnitude of transactivation may be tuned by careful selection, or screening in the absence of prior knowledge, of nucleotides flanking the ZF binding site(s) in a cognate promoter. Similarly, it is not yet possible to know for certain whether a new ZFa will be orthogonal to ZFa and promoters that are already characterized, although the analyses presented here provide a blueprint as to how such new incorporations may be evaluated and quantitatively compared. We expect that the specific quantitative parameters determined in this study may be limited to the implementations used here, including the methods for DNA delivery and the cell type in which the characterizations were performed. However, since the fundamental mechanisms of transcription are maintained across contexts, we expect that the observed trends will extend across cell types and delivery methods. For instance, the rank order of promoter strength across the number of binding sites was conserved between transient transfection and genomic integration (**Fig. 6**). A practical method of evaluating such applications in general would be to select a focused library of COMET parts, guided by the characterizations here, to empirically identify which combinations provide the desired function, and if needed, further tune system performance using the strategies described in this study.

A particularly exciting opportunity is to use COMET with other synthetic biology technologies. For example, COMET could be integrated into synthetic receptors that utilize orthogonal TFs as outputs, such as MESA or synNotch, to generate cellular programs for sensing, processing, and responding to environmental cues^25, 36–38^. Alternatively, COMET could be used to regulate the expression of synthetic components, such as GEMS receptors, that interface with endogenous signaling pathways^39^. In general, we expect that COMET will be useful for prototyping and implementing sophisticated cellular functions for both fundamental research and cellular engineering applications.

## Supporting information

Plasmid map archive

Software files archive

Supplementary Table Archive

## Acknowledgments

This work was supported in part by the National Cancer Institute (NCI) of the National Institutes of Health (NIH) under Award Number F30CA203325; the National Institute of Biomedical Imaging and Bioengineering of the NIH under Award Number 1R01EB026510; the National Institute of General Medical Sciences (NIGMS) of the NIH under Award Number T32GM008152 (to Hossein Ardehali); the Northwestern University Flow Cytometry Core Facility supported by Cancer Center Support Grant (NCI 5P30CA060553); and the Northwestern Institute for Cellular Engineering Technologies (iCET). The content is solely the responsibility of the authors and does not necessarily represent the official views of the NIH. The authors would like to thank Everett Allchin, Lauren Battaglia, Pear Dhiantravan, Benjamin Leibowitz, Sophia Li, Siyuan Feng, and Viswajit Kandula for their assistance with cloning in this paper, and Amy Hong, Cameron McDonald, Justin Finkle, and Sebastian Bernasek for useful discussion on the computational model. We thank Ahmad Khalil (Boston University) for sharing plasmids encoding the ZFa from his 2012 study^15^.

## Author Contributions

P.S.D. and J.N.L conceptualized COMET. P.S.D, J.W.D, J.J.M., and H.I.E created reagents, designed and performed experiments, and analyzed the data. J.J.M developed the computational models and code. P.S.D and J.J.M drafted the original manuscript, and P.S.D, J.W.D, H.I.E, and J.J.M created the figures. J.N.L. and N.B. supervised the work. All authors edited and approved the final manuscript.

## ONLINE METHODS

### General *DNA assembly*

Plasmid cloning was performed primarily using standard PCR and restriction enzyme cloning with Vent DNA Polymerase (New England Biolabs (NEB)), *Taq* DNA Polymerase (NEB), Phusion DNA Polymerase (NEB), restriction enzymes (NEB; Thermo Fisher), T4 DNA Ligase (NEB), Antarctic Phosphatase (NEB), and T4 PNK (NEB). Golden gate assembly and Gibson assembly were also utilized. Most plasmids were transformed into chemically competent TOP10 *E. coli* (Thermo Fisher) and grown at 37°C, except for integration vectors, which were transformed into chemically competent Stable *E. coli* (NEB) and grown at 30°C.

### Cloning strategy for COMET vectors

The COMET plasmids are in pcDNA backbones for high expression in HEK293FT cells. Restriction sites were chosen to allow for modular swapping of parts with restriction enzyme cloning. Furthermore, reporter constructs can be assembled by one-step Golden Gate reactions employing synthesized oligonucleotides. A complete list of all plasmids constructed for and utilized in this manuscript is available in **Supplementary Tables 1–8**, and plasmid maps are available per **Supplementary Note 3**.

### Source vectors for DNA assembly

ZF-containing and VP16-containing vectors were a generous gift from Ahmad Khalil^1^. VP64 and VPR were sourced from SP-dCas9-VPR, which was a gift from George Church (Addgene plasmid # 63798)^2^. DsRed-Express2 was obtained by site directed mutagenesis of pDsRed2-N1, which was a gift from David Schaffer (University of California, Berkeley). EBFP2 was sourced from pEBFP2-Nuc, which was a gift from Robert Campbell (Addgene plasmid #14893)^3^. EYFP, FKBP, and FRB were sourced from plasmids we previously described (Addgene plasmids #58855, #58877, and #58876, respectively)^4^. NanoLuciferase was synthesized as a GeneArt DNA String (Life Technologies/Thermo Fisher). The mMoClo (pLInk2, pLink4, and pLink8, Destination Vector, BxB1 Recombinase Expression Vector) plasmids were a gift from Ron Weiss^5^. The CHS4 insulator was sourced from PhiC31-Neo-ins-5xTetO-pEF-H2B-Citrin-ins, which was a gift from Michael Elowitz (Addgene plasmid #78099)^6^. The CAG promoter was sourced from pR26R CAG/GFP Asc, which was a gift from Ralf Kuehn (Addgene plasmid #74285)^7^. The SV40 minimal promoter was sourced from pYC0866 (4xHRE_minSV40-sfGFP-CMV_dsRed Exp), which was a gift from Yvonne Chen^8^. EF1α and TetON3G were sourced from pLVX-Tet3G (Clontech), and TRE3GV was sourced from pLVX-TRE3G (Clontech). Barcodes used for the TUPVs were designed by the Elledge lab^9^. BlastR was sourced from lenti dCAS-VP64_Blast, which was a gift from Feng Zhang (Addgene plasmid #61425)^10^.

### Plasmid backbones

All plasmid backbones are modified versions of the pcDNA3.1/Hygro(+) Mammalian Expression Vector (Thermo Fisher V87020).

- To make pPD003, the SV40 promoter and Hygromycin resistance gene that it drove were removed, while leaving the SV40 origin of replication and SV40 poly(A) signal intact. Additionally, a sense mutation in the *AmpR* gene was introduced to remove a BsaI restriction site.
- To make pPD005 (referred to as “pcDNA”), the BpiI site in the bGH poly(A) signal was mutated to enable Golden Gate reactions with BpiI, and the BsaI site in the 5’-UTR was mutated to enable Golden Gate reactions with BsaI. The BpiI site was in a region of the BGH poly(A) tail that when deleted does not alter the efficiency of the polyadenylation^11^.

### Template plasmids for ZF reporter plasmids

- pPD027 (the first-generation ZF reporter template) was constructed by inserting a synthesized region (containing two BsaI sites for Golden Gate-mediated ZF binding site array insertion and a YB_TATA minimal promoter^8^) between the BglII and NheI sites and inserting EYFP between the NheI and NotI sites of pPD003.
- pPD032 and pPD033, which are the templates for ZF reporters with the binding site array moved further upstream of the minimal promoter, were constructed by inserting spacer regions into the BamHI site between the ZF binding array insertion template and the YB_TATA minimal promoter. The spacer inserts were amplified by PCR from the region of pPD003 upstream of the CMV promoter prior to insertion. These three templates (pPD027, pPD032, and pPD033) were used to construct all spaced reporters shown in Fig. 1b–e and Fig. 2a.
- pPD152 (the second-generation ZF reporter template) was constructed to enable multi-round insertion of larger ZF binding arrays using alternating rounds of Golden Gate with BsaI and BpiI. To do so, the region of pDPD027 between the AatII and NotI sites (the ZF binding array insertion site through the end of the EYFP coding sequence) was inserted between the corresponding sites of pPD005. pPD152 was used to make all of the ZF1 compact binding site reporters shown in Fig. 1 and Fig. 2a (including pPD290 (ZF1×6-C YB_TATA EYFP), which was used as the reporter plasmid in the majority of the experiments), and the logic promoters in Fig. 8.
- pPD540 (the third-generation ZF reporter template) was constructed to swap the palindromic “sticky ends” (5’ or 3’ overhangs) of the ZF binding array insertion site to non-palindromic sticky ends. The use of palindromic sticky ends, which were originally designed to allow construction of ZF binding arrays with either Golden Gate or EcoRI and BamHI, risks the insertion of multiple copies of the same insert in Golden Gate reactions. This redesign enabled us to inset promoters of sizes that could not be cheaply synthesized as a single insert as multiple inserts in a single round of Golden Gate. This was accomplished by synthesizing a new upstream region (containing two BsaI sites for Golden Gate-mediated ZF binding site array insertion with non-palindromic sticky ends and a YB_TATA minimal promoter) and inserting this upstream region between the BglII and NheI sites of pPD152.

### Golden Gate assembly of ZF reporter plasmids

Golden Gate assembly^12^ was used to construct most of the reporter plasmids from a reporter template. Promoter insets were synthesized as 15–100 bp oligonucleotides (some promoters were synthesized as multiple inserts) by Integrated DNA Technologies or Life Technologies (Thermo Fisher). The coding and reverse strands were synthesized separately and designed to anneal, resulting in dsDNA with a 4 nt sticky end overhang on each side. The coding and reverse oligonucleotides were mixed (6.5 µL H_2_O, 1 µL T4 Ligase Buffer, 0.5 µL T4 PNK (10 U/µL; NEB), 1 µL of each 100 µM oligonucleotide) and phosphorylated at 37°C for 1 h. They were then denatured at 95°C for 5 min and cooled slowly to room temperature (here, approximately 22°C) to allow for annealing. The mix was then diluted 50-fold to make a 200 nM stock or 500-fold to make a 20 nM stock. While we made most of the constructs with the 200 nM stock, we later discovered that the 20 nM stock resulted in higher-efficiency reactions.

BsaI Golden Gate reaction mixtures comprise 1 µL T4 ligase buffer, 1 µL 10x BSA (1 mg/mL), 0.5 µL BsaI-HF (20 U/µL; NEB), 0.5 µL T4 Ligase (400 U/µL; NEB), 10 fmol of vector, 1 µL of each insert (diluted to 200 nM or 20 nM), and water to 10 µL total volume. The reaction was incubated at 37°C for 1 h, 55°C for 15 min, and 80°C for 20 min, and then cooled to room temperature. Up to 10 µL of reaction was immediately transformed into up to 50 µL of chemically competent Top10 *E. coli*. For reactions that did not yield many colonies on the first cloning attempt or did not produce colonies with the correct plasmids, the reaction conditions were changed to: 30 cycles of 37°C for 1 minute then 16°C for 1 minute, 55°C for 15 min, 80°C for 20 min, and cool to room temperature.

Some of the larger ZF binding site arrays were assembled through sequential rounds of alternating BsaI and BpiI Golden Gate reactions. BpiI Golden Gate reaction mixtures comprise 1 µL T4 ligase buffer, 1 µL 10x BSA (1 mg/mL), 0.4 µL BpiI-FD (Thermo Fisher), 0.4 µL µL T4 Ligase (400 U/µL; NEB), 10 fmol of vector, 1 µL of each insert (diluted to 200 nM or 20 nM), and water to 10 µL. The reaction was incubated at 37°C for 30 min, 50°C for 5 min, and 80°C for 10 min, and then cooled to room temperature prior to transformation.

### Non-Golden Gate assembly of some ZF reporters

Although Golden Gate assembly was the primary strategy for cloning the promoters, the first-generation templates were not readily amenable to synthesis and insertion of ZF binding site arrays. Therefore, some spaced promoters with large numbers of binding sites used in Fig. 1c were constructed by PCR amplification of 1–8 binding sites from other reporter plasmids and insertion of these binding sites between the EcoRI and BamHI sites upstream of reporter constructs with 1–8 binding sites in the promoter. Likewise, the ZF reporters with ZF binding site arrays moved further upstream of the minimal promoter shown in Fig. 1d were constructed by PCR-amplifying the ZF binding site arrays from other constructs and inserting between the EcoRI and BamHI sites of pPD032 and pPD033.

Additionally, COMET reporter constructs were designed to include a limited set of minimal promoters; restriction enzyme cloning was employed to accomplish this as well. pPD1028 (ZF1×6-C SV40_Min EYFP) was cloned from pPD270, cut with XbaI and ApaI. Into this construct we inserted two fragments of DNA: SV40_min^8^ was PCR-amplified and cut with BsaI and ApaI, and EYFP was PCR-amplified from pPD270 and cut with BsaI and ApaI. pPD1029 (ZF1×6-C CMV_min EYF) was cloned from pPD270, cut with XbaI and ApaI. CMV_min was synthesized by IDT and placed upstream of an EYFP gene in a pcDNA-based vector. A fragment comprising CMV_min and EYFP was then PCR-amplified, digested with BsaI and ApaI, and inserted into the digested pPD270.

### Assembly of ZFa and ZFi

The first five ZFa tested in Fig. 1b were constructed by PCR-amplifying the ZFa sequence from^1^ (including the N-terminal 3x-FLAG tag, SV40 NLS, VP16 AD, and ZF) and inserting between the NheI site and NotI site of pPD005. Cognate ZFi were constructed by whole-plasmid PCR-mediated deletion of the VP16 AD. During the AD deletion process, BamHI and KpnI sites were added between the SV40 NLS and the ZF, which were later used to insert a PCR-amplified DsRed Express2, thereby creating cognate ZFi-DsRed. Subsequent ZFa (*i.e.*, any new ZFa tested in Fig. 3a) were constructed by replacing DsRed-Express2 with a PCR-amplified VP16 (BamHI/KpnI) and replacing the ZF domain with a PCR-amplified ZF domain (KpnI/NotI) from^1^.

### Assembly of ZF mutants

ZFa mutants were synthesized as multiple sets of complementary oligonucleotides, which were annealed and then inserted via Golden Gate assembly into a vector designed to encode ZFa upon insertion of all inserts. Reactions were performed with BpiI as described in ***Golden Gate assembly of ZF reporter plasmids***. ZFi mutants were generated by whole-plasmid PCR-mediated deletion of the VP16 AD.

### Assembly of RaZFa

RaZFa components were constructed by multi-step restriction enzyme-based cloning. The SV40 NLS was part of the original ZFa constructs^1^, and the NES sequence was obtained from^13^.

### Gibson assembly

Gibson assembly^14, 15^ was used to specify ADs on ZF1a. Gibson reactions were performed by PCR addition of homology arms onto the target DNA. Components were mixed together: 17 fmol of backbone, 51 fmol of each insert, 7.5 μL of Gibson Master Mix, and water to 10 µL. 7.5 µL of Gibson Master mix contains 2 μL 5X isothermal reaction buffer (0.5 M Tris-HCl pH 7.5, 0.05 M MgCl_2_, 1 mM dNTP, 5 mM NAD, 0.05 M DTT), U T5 exonuclease, 0.25 U Phusion DNA Polymerase, and 40 U *Taq* DNA Ligase (NEB) in water. The reaction was incubated at 50°C for 1 h, and 5 µL was transformed into chemically competent Top10 *E. coli* (Thermo Fisher). In subsequent cases, ADs were moved onto other ZF by restriction digest.

### Construction of plasmids for mMoClo

We made several changes to the mMoClo plasmids originally described^5^ in order to incorporate them into the workflow for our laboratory, in which many constructs are prototyped using pcDNA-based expression vectors. Details can be found in **Supplementary Fig. 9** and **Supplementary Tables 5, 6**. We modified the Destination Vector provided by the Weiss lab by adding two repeats of the CHS4 insulator into two places in the vector. The insulators upstream of the attB site are, upon genomic integration, inserted downstream of the LP, insulating the LP from the genome (and vice versa). The insulators downstream of the RB Globin polyA terminator of the puromycin resistance gene insulate this transcription unit from TU1. This new vector is termed pPD630 (Integration Vector). We cloned pLink1, pLink3, pLink5, pLink6, pLink7, and pLink9 site directed mutagenesis via whole plasmid PCR of pLink2.

The TUPVs were cloned by making several alterations to pcDNA (pPD005), in 3 steps. In the first step, two repeats of the CHS4 insulator were placed downstream of the BGH polyA tail. Second, to enable Golden Gate cloning of the TUPV library, three pairs of BsaI sites were inserted into the vector with PCR. The first pair was upstream of the promoter, the second pair was inserted between the BGH polyA tail and the insulator, and the third pair was inserted downstream of the insulator. In the third reaction, three pairs of annealed oligonucleotides were inserted into these BsaI sites via a Golden Gate reaction. The first insert, to be placed upstream of the promoter, comprised a BpiI site, TUPV-specific sticky end, and TUPV-specific 5’ barcode (barcodes unique to each TUPV enable sequencing of the TUPV contents after TUPVs are combined into an integration vector). The second insert, to be placed between the BGH/polyA tail and the insulator, comprised a TUPV-specific 3’ barcode. The third insert, to be placed downstream of the insulator, comprised a BpiI site and a TUPV-specific sticky end. In this manner, 9 TUPVs each with their own unique 5’ and 3’ barcodes and 5’ and 3’ sticky ends were cloned (pPD471–479). This initial library uses a CMV promoter as the core promoter for each TU, which was placed upstream of a multiple cloning site (MCS). A second library of 9 TUPVs was then constructed by replacing the CMV promoter with the CAG promoter by restriction enzyme digest with SnaBI and NheI (pPD561–569). A third library of 9 TUPVs was constructed by replacing the promoter with EF1alpha (between MluI and NheI) and the MCS replaced with an EBFP2-P2A-BlastR gene (between NheI and NotI) (pJM450–458). Although this third library no longer contains the full pcDNA MCS, it retains the NheI and NotI genes that flank the COMET ZFa and ZFi and many of the RaZFa components.

### Transferring COMET parts into mMoClo

COMET reporters and ZFa were transferred into TUPVs using restriction enzyme cloning. To construct mKate2 reporters in TUPV1, mKate2 was cloned into the MCS of pPD561 using NheI and NotI restriction sites downstream of a CAG promoter to create pHIE041. Binding site arrays were PCR-amplified from pPD152, pPD287, pPD290, pPD296, pPD063, pPD069, and pPD095 and inserted to replace CAG in pHIE041 using BglII and NheI, resulting in pHIE042–049. To construct constitutively expressed VP16-ZF1a in TUPV2 (pJM466), the EBFP-P2A-BlastR in pJM451 was replaced with PCR-amplified VP16-ZF1a from pD100 using NheI and NotI.

### mMoClo Assembly of Integration Vectors

The mMoClo integration vectors were assembled through a BpiI-mediated Golden Gate reaction. Each 20 μL reaction comprised 2 µL 10x T4 ligase buffer, 2 µL 10x BSA (1 mg/mL stock), 0.8 µL BpiI-FD, 0.8 µL T4 DNA Ligase (400 U/µL stock), 20 fmol integration vector backbone (pPD630), and 40 fmol of each transcription unit and linker plasmid to be inserted. The reaction was incubated at 37°C for 15 min, then subjected to 55 iterations of thermocycling (37°C for 5 min, 16°C for 3 min, repeat), followed by 37°C for 15 min, 50°C for 5 min, 80°C for 10 min to terminate the reactions; then the mixture was cooled to room temperature (optionally held at 4°C if the reaction ran overnight) and placed on ice prior to immediate transformation into bacteria.

### Plasmid preparation

TOP10 *E. coli* were grown overnight in 100 mL of LB with the appropriate selective antibiotic. The following morning, cells were pelleted at 3000 x g for 10 min and then resuspended in 4 mL of a solution of 25 mM Tris pH 8.0, 10 mM EDTA, and 15% sucrose. Cells were lysed for 15 min by addition of 8 mL of a solution of 0.2 M NaOH and 1% SDS, followed by neutralization with 5 mL of 3 M sodium acetate (pH 5.2). Precipitate was pelleted by centrifugation at 9000 x g for 20 min. Supernatant was decanted and treated with RNAse A for 1 h at 37°C. 5 mL of phenol chloroform was added, and the solution was mixed and then centrifuged at 7500 x g for 20 min. The aqueous layer was removed and subjected to another round of phenol chloroform extraction with 7 mL of phenol chloroform. The aqueous layer was then subjected to an isopropanol precipitation (41% final volume isopropanol, 10 min at room temperature, 9000 x g for 20 min), and the pellet was briefly dried and resuspended in 420 µL of water. The DNA mixture was incubated on ice for at least 12 h in a solution of 6.5% PEG 20,000 and 0.4 M NaCl (1 mL final volume). DNA was precipitated with centrifugation at maximum speed for 20 min. The pellet was washed once with ethanol, dried for several h at 37°C, and resuspended for several h in TE buffer (10 mM Tris, 1 mM EDTA, pH 8.0). DNA purity and concentration were confirmed using a Nanodrop 2000 (Thermo Fisher).

### Cell culture

HEK293FT cells and HEK293FT-LP cells were cultured in DMEM (Gibco 31600-091) with 10% FBS, 6 mM L-glutamine (2 mM from Gibco 31600-091 and 4 mM from additional Gibco 25030-081), penicillin (100 U/μL), and streptomycin (100 μg/mL) (Gibco 15140122), in a 37°C incubator with 5% CO_2_. Cells were subcultured at a 1:5 to 1:10 ratio every 2–3 d using Trypsin-EDTA (25300-054). The HEK293FT-LP line was a gift from Ron Weiss.^5^

### Transfection experiments

Experiments were conducted by transient transfection of HEK293FT cells using the calcium phosphate method. For transfection experiments, cells were plated at a minimum density of 1.5 x 10^5^ cells/well in a 24-well plate in 0.5 mL of DMEM, supplemented as described above. After at least 6 h, by which time the cells had adhered to the plate, they were transfected via the calcium phosphate method. Plasmids for each experiment were mixed in H_2_O, and 2 M CaCl_2_ was added to a final concentration of 0.3 M CaCl_2_. The exact DNA amounts added to the mix per well and plasmid details for each experiment are listed in the following sections and can be cross-referenced with **Supplementary Tables 11–18** for further details. This mixture was added dropwise to an equal-volume solution of 2x HEPES-Buffered Saline (280 mM NaCl, 0.5 M HEPES, 1.5 mM Na_2_HPO_4_) and gently pipetted up and down four times. After 2.5–4 min, the solution was mixed vigorously by pipetting eight times. 100 µL of this mixture was added dropwise to the plated cells, and the plates were swirled gently. The next morning, the medium was aspirated and replaced with fresh medium. In some assays, fresh medium contained 0.05% DMSO or 0.05% DMSO with 0.1 µM rapamycin. At 36–48 h post-transfection and at least 24 h post-media change, cells were harvested for flow cytometry with FACS Buffer (PBS pH 7.4 with 2–5 mM EDTA and 0.1% BSA) or with Trypsin-EDTA, which was then quenched with medium, and the resulting cell solution was added to at least 2 volumes of FACS buffer. Cells were spun at 150 x g for 5 min, FACS buffer was decanted, and fresh FACS buffer was added. All experiments were performed in biologic triplicate.

### Western Blotting

For western blotting, HEK293FT cells were plated at 7.5 x 10^5^ cells/well in 2 mL of DMEM and transfected as above, using 400 μL of transfection reagent per well (the reaction scales with the volume of medium). At 36–48 h after transfection, the cells were lysed with 500 μL of RIPA (150 mM NaCl, 50 mM Tris-HCl pH 8.0, 1% Triton X-100, 0.5% sodium deoxycholate, 0.1% sodium dodecyl sulfate) with protease inhibitor cocktail (Pierce/Thermo Fisher cat# A32953) and incubated on ice for 30 min. The lysate was cleared by centrifugation at 14,000 x g for 20 min at 4°C and the supernatant was harvested. A BCA assay was performed to determine protein concentration, and after a 10-minute incubation with Lamelli buffer (final concentration 60 mM Tris-HCl pH 6.8, 10% glycerol, 2% sodium dodecyl sulfate, 100 mM dithiothreitol, and 0.01% bromophenol blue) at 70°C, 0.5 μg of total protein was loaded onto a 4-15% Mini-PROTEAN TGX Precast Protein Gel (Bio-Rad) and run at 50 V for 10 min followed by 100 V for at least 1 h. Wet transfer was performed onto an Immuno-Blot PVDF membrane (Bio-Rad) for 45 min at 100 V. Ponceau-S staining was used to confirm successful transfer. Membranes were blocked for 30 min with 3% milk in Tris-buffered saline pH 8.0 (TBS pH 8.0: 50 mM Tris, 138 mM NaCl, 2.7 mM KCl, HCl to pH 8.0), washed once with TBS pH 8.0 for 5 min, then incubated for 1 h at room temperature or overnight at 4°C in primary solution antibody (Mouse-anti-FLAG M2 (Sigma F1804, RRID: AB_262044), diluted 1:1000 in 3% milk in TBS pH 8.0). Primary antibody solution was decanted, and the membrane was washed once with TBS pH 8.0 then twice with TBS pH 8.0 with 0.05% Tween, for 5 min each. Secondary antibody (HRP-anti-Mouse (CST 7076, RRID: AB_330924), diluted 1:3000 in 5% milk in TBST pH 7.6 (TBST pH 7.6: 50 mM Tris, 150 mM NaCl, HCl to pH 7.6, 0.1% Tween)) was applied for 1 h at room temperature, and the membrane was washed three times for 5 min each time with TBST pH 7.6. The membrane was incubated with Clarity Western ECL Substrate (Bio-Rad) for 5 min, and then exposed to film, which was developed and scanned. Images were cropped with Photoshop CC (Adobe). No other image processing was employed. Original images are available on request.

The western blot shown in **Supplementary Fig. 12f** was conducted twice with comparable results. The first experiment included only the RaZFa component (no additional loading control) to confirm the presence of only one band in each lane (data not shown). In the second experiment, 40 ng of pPD798 (encoding a 3X-FLAG tagged NanoLuciferase) was co-transfected with the RaZFa components to provide a control for loading and transfection.

### Analytical flow cytometry

Flow cytometry was run on a BD LSRII or BD LSR Fortessa Special Order Research Product (Robert H. Lurie Cancer Center Flow Cytometry Core). The following lasers and filter sets were used for data acquisition:

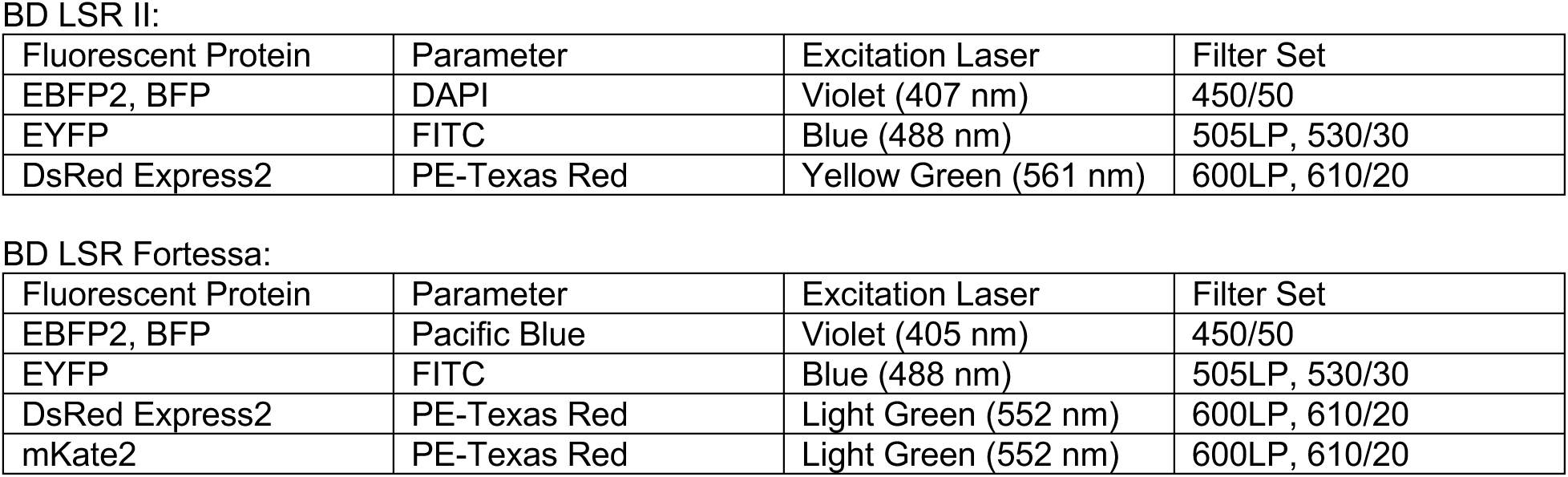

**BD LSR II:**
| Fluorescent Protein | Parameter | Excitation Laser | Filter Set |
| --- | --- | --- | --- |
| EBFP2, BFP | DAPI | Violet (407 nm) | 450/50 |
| EYFP | FITC | Blue (488 nm) | 505LP, 530/30 |
| DsRed Express2 | PE-Texas Red | Yellow Green (561 nm) | 600LP, 610/20 |

**BD LSR Fortessa:**
| Fluorescent Protein | Parameter | Excitation Laser | Filter Set |
| --- | --- | --- | --- |
| EBFP2, BFP | Pacific Blue | Violet (405 nm) | 450/50 |
| EYFP | FITC | Blue (488 nm) | 505LP, 530/30 |
| DsRed Express2 | PE-Texas Red | Light Green (552 nm) | 600LP, 610/20 |
| mKate2 | PE-Texas Red | Light Green (552 nm) | 600LP, 610/20 |

Approximately 2,000–3,000 single, transfected cells were analyzed per sample.

### Flow Cytometry Data Analysis

Samples were analyzed using FlowJo v10 software (FlowJo, LLC). As illustrated in **Supplementary Fig. 15**, the HEK293FT cell population was identified by FSC-A vs. SSC-A gating, and singlets were identified by FSC-A vs. FSC-H gating. To distinguish transfected and non-transfected cells, a control sample of cells was generated by transfecting cells with a mass of pcDNA (empty vector) equivalent to the mass of DNA used in other samples in the experiment. For the single-cell subpopulation of the pcDNA-only sample, a gate was made to identify cells that were positive for the constitutively driven fluorescent protein used as a transfection control in other samples, such that the gate included no more than 1% of the non-fluorescent cells. The mean fluorescence intensity (MFI) of the single-cell transfected population was calculated and exported for further analysis.

To calculate reporter expression, MFI in the FITC channel was averaged across three biologic replicates. From this number, the autofluorescence of the cells was subtracted. To calculate the autofluorescence of the cells, in each experiment, a control group of cells transfected with DNA encoding the fluorescent protein transfection control and pcDNA were used. The background-subtracted MFI was converted to Mean Equivalents of Fluorescein (MEFLs) by multiplying by a coefficient determined in each experiment, as described below. Standard error was propagated through all calculations.

### Conversion of arbitrary units to standardized fluorescence units

As shown in **Supplementary Fig. 16**, to determine the conversion factor for MFI to MEFLs, Rainbow Calibration Particles (Spherotech, RCP-30-5) or UltraRainbow Calibration Particles (Spherotech URCP-100-2H) were run with each flow cytometry experiment. This reagent contains six (RCP) or nine (URCP) subpopulations of beads, each of a specific size and with a known number of various fluorophores. The total bead population was identified by SSC vs. FSC gating, and the subpopulations were identified through two fluorescent channels. The MEFL values corresponding to each subpopulation were supplied by the manufacturer. A calibration curve was generated for the experimentally determined MFI vs. manufacturer supplied MEFLs, and a linear regression was performed with the constraint that 0 MFI equals 0 MEFLs. The slope from the regression was used as the conversion factor, and error was propagated.

### Integration of cargo into landing pad cell lines

From exponentially growing HEK293LP cells, 0.5 x 10^5^ cells were plated per well (0.5 mL medium) in 24-well format, and cells were cultured for 24 h to allow cells to attach and spread. When cells reached 50– 75% confluence, Bxb1 recombinase was co-transfected with the integration vector by lipofection with Lipofectamine LTX with PLUS Reagent (ThermoFisher 15338100). 300 ng of BxB1 expression vector was mixed with 300 ng of integration vector and 0.5 μL of PLUS reagent in a 25 μL total volume reaction, with the remainder of the volume being OptiMEM (ThermoFisher/Gibco 31985062). In a separate tube, 1.9 μL of LTX reagent was mixed with 23.1 μL of OptiMEM. The DNA/PLUS Reagent mix was added to the LTX mix. pipetted up and down four times, and then incubated at room temperature for 5 min. 50 μL of this transfection mix was added drop-wise to each well of cells, which was mixed by gentle swirling. Cells were cultured until the well was ready to split (typically 3 d), without any media changes.

### Selection and expansion of landing pad cell lines

Cells were harvested from the 24-well plate when confluent by trypsinizing and transferring to a single well of a 6-well plate in 2 mL of medium, and then cells were cultured until they reached 50-70% confluence. Then, medium was aspirated and replaced with 2 mL of fresh media containing appropriate selection antibiotic 1 μg/mL puromycin (Invivogen ant-pr) or 6 μg/mL blasticidin (Alfa Aesar/ThermoFisher J61883). Medium was replaced daily with fresh medium containing antibiotics until cell death was no longer evident. Selection was first performed in puromycin for 7 d, then cells were expanded for 7 d without antibiotics. Cells were then cultured in both puromycin and blasticidin to maintain selective pressure until flow sorting.

### Sorting of landing pad cell lines

Cells were harvested by trypsinizing, resuspended at approximately 10^7^ cells per mL in pre-sort medium (DMEM with 10% FBS, 25 mM HEPES (Sigma H3375), and 100ug/mL gentamycin (Amresco 0304)), and held on ice until sorting was performed. Cells were sorted using a BD FACS Aria 4-laser Special Order Research Product (Robert H. Lurie Cancer Center Flow Cytometry Core) with the following optical configuration:

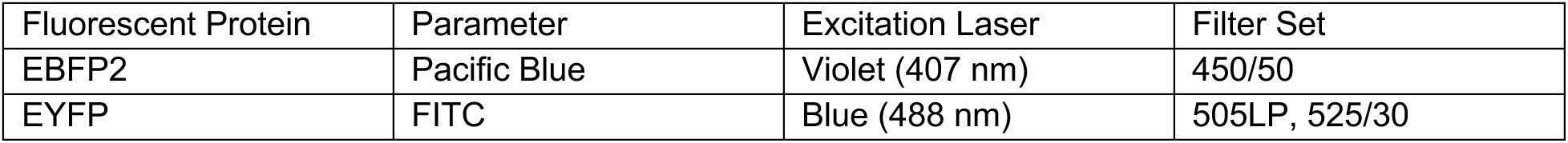

The sorting strategy was as follows: single cells were first gated to exclude all EYFP positive cells (as EYFP positive cells still have an intact landing pad locus, suggesting a mis-integration event occurred) and to include only EBFP2+ cells. Then a gate was drawn on EBFP2 expression, utilizing the line that demonstrated the least amount of silencing (ZF1×12-C_mKate2 + ZF1a) to capture the 90^th^ to 98^th^ percentile of EBFP2 expressing cells (the top 2% were excluded to exclude cells suspected to possess two or more integrated copies of the cargo vector). The gate drawn using this line was used for all other lines as well. No gating was performed on mKate2 reporter expression. 15,000 cells were collected for each line in post-sort medium (DMEM with 20% FBS, 25 mM HEPES, and 100 μg/mL gentamycin), and cells were held on ice until they could be centrifuged at 150 x g for 5 min and resuspended in DMEM. Cells were plated in a 24-well plate and expanded until used in experiments. Gentamycin was included in the culture medium for one week after sorting.

### Experiments involving landing pad cell lines

Stable cell lines were plated in 0.5 mL of DMEM in triplicate in 24-well format at a density expected to generate 50% confluent wells. The day after plating (24 h), cells were harvested with Trypsin-EDTA, as described in **Transfection Experiments**. For transfection experiments designed to accompany landing pad line experiments (e.g., Fig. 6 a,b), cells were plated and transfected 2 d prior to the assay and harvested as described in **Transfection Experiments**. Flow cytometry was run on a BD LSR Fortessa, as described in **Analytical flow cytometry**.

For characterization, approximately 10,000 single, EBFP2-expressing cells were analyzed per sample, where EBFP2 is a marker for locus activity in the stable cells and a transfection control for transfected cells. Stable cells were analyzed using higher laser voltages than those used for transfected cells to effectively capture the range of reporter expression conferred by the panel of COMET promoters; thus the results from this experiment are displayed in separate panels of **Fig. 6** despite the fact that data collection occurred on the same day.

### COMET Model Development and Analysis

This section provides an integrated discussion of model development, calibration, and analysis, to supplement the discussion in the main text. We first describe the development of the core model, and then discuss elaborations and models used for comparison. In developing the core model to investigate and predict COMET behavior, we account for two high level phenomena: cell heterogeneity using a statistical model, and gene regulation using a dynamical model.

#### Statistical model

Heterogeneity is represented by simulating genetic programs in a way that resembles their outcomes in cells, which vary in the expression of the components. The *in silico* population (**Z**) is an *N* x *P* matrix, where *N* is the number of cells (*n* = 1:*N*, for N = 200) and *P* is the number of plasmids (*p* = 1:*P*). Components that are encoded on separate plasmids are assigned separate columns. For example, ZF1a is assigned one column and the reporter is assigned another column.

**Z** is generated using the constrained sampling method, as described previously^16^. Briefly, this algorithm proceeds as follows:

1. Specify parameters for the target marginal distribution of gene expression.
  - Based on flow cytometric measurements of constitutively expressed fluorescent proteins from transfected plasmids, the distribution for each protein was log-bimodal Gaussian. This distribution can be described by the parameters *μ*_1_ = 1.95, *σ*_1_ = 0.3, *μ*_2_ = 3.4, and *σ*_2_ = 0.6 arbitrary units (a.u.) on a log_10_-scaled axis.
2. Specify a target correlation coefficient to model the expression of genes from co-transfected plasmids.
  - A Pearson correlation of *r* = 0.8 was used based on experimentally determined correlations.
3. Based on the target correlation, specify a lower bound and upper bound of acceptable values.
  - Values should be chosen that are close to the target, such as 0.765 and 0.835.
4. Generate a joint distribution using the parameters for the marginal distribution and the target correlation coefficient. This output is a candidate matrix for population variation.
  - Distributions can be generated using the multivariate normal random number generator (`mvnrnd`) in MATLAB.
5. Compute the correlation coefficient matrix (*P* x *P*).
6. While any non-diagonal entries in the correlation coefficient matrix are outside of the range of acceptable values specified in step 3, repeat steps 4 and 5.
7. For the accepted matrix, normalize the values in each column to a mean of one, to obtain the population matrix **Z**.
  - We recommend plotting the plasmid distributions and correlations to confirm their resemblance to the target outcomes.

#### Dynamical model of ZFa-mediated transcription

Gene regulation is represented using a system of ODEs. The example below depicts a ZFa inducing a reporter. Transcription of ZFa RNA is constitutive and scales linearly with plasmid dose. Transcription of reporter RNA depends on ZFa protein concentration as described by a transfer function *f*, which is described in subsequent sections. RNA degradation, protein translation, and protein degradation are represented as first-order processes. The *k* terms are fixed parameters that are either defined as equal to 1 unit or are based on a previous estimate: *k_transcription_* = 1 arbitrary transcription unit, *k_degRNA_* = 2.7 h^-1^ based on a previous study^17^, *k_translation_* = 1 arbitrary translation unit, *k_degZFa_* = 0.35 h^-1^ based on another study^18^, and *k_degReporter_* = 0.029 h^-1^ based on another study^19^.

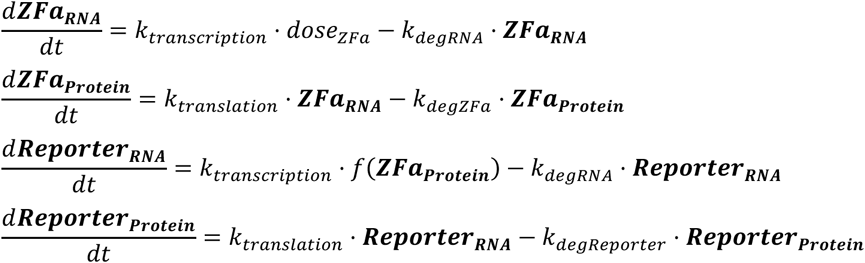

Although the rate constants for transcription and translation for both the ZFa and reporter are 1 a.u., in practice these processes likely differ between genes. As a result, 1 a.u. of ZFa RNA can correspond to a different number of molecules in a living cell than 1 a.u. of reporter RNA, and likewise for 1 a.u. of each protein. However, importantly, 1 a.u. for a given species (*e.g.*, reporter protein) *can* be treated as equivalent across simulation conditions (*e.g.*, varied ZFa plasmid doses), and these are comparisons of greatest interest in our analysis.

As described in the main text, for a ZFa-inducible promoter, *f* is defined as:

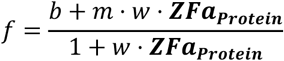

where *b* is a non-negative value for TF-independent (leaky or background) transcription; *m* is a unitless value for maximum activation (for ZF1a, *m* ≥ 1) that depends on the number and spacing of binding sites and the TF; and *w* is a positive value for the steepness of the ZFa dose response. The ZFa variable refers to the simulated protein level, which is a function of plasmid dose, but is in distinct units from and is not equivalent to plasmid dose.

Experiments showed that leaky expression from COMET promoters (across varying numbers of ZF1 sites) was ∼10% of the inducible expression observed with a high ZF1a dose and x1 (one binding site) promoter.

The parameter *m* describes the maximum transcription that a specific ZFa can drive at a promoter with a specific architecture (number and spacing of binding sites). An *m* value of 1 is defined for ZF1a with a x1 promoter. We found that values for *m* vary with the numbers of binding sites (*BS*) in a manner that can be represented by sigmoid functions, as shown below for ZF1a. The max argument ensures that *m* does not go below 1 and that it increases monotonically with the number of binding sites. Thus, for ZF1a:

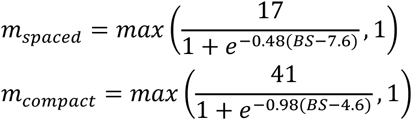

For TFs such as ZF2a that follow similar binding site-response behavior, we find that the sigmoids appear vertically stretched or squashed. This change can be represented through the numerator in the fraction for the *m* function, *e.g.*, ZF2a with a compact promoter has a numerator of 60.

#### Calibration

The full model was implemented in MATLAB with modifications to the terms for RNA production to account for cell heterogeneity:

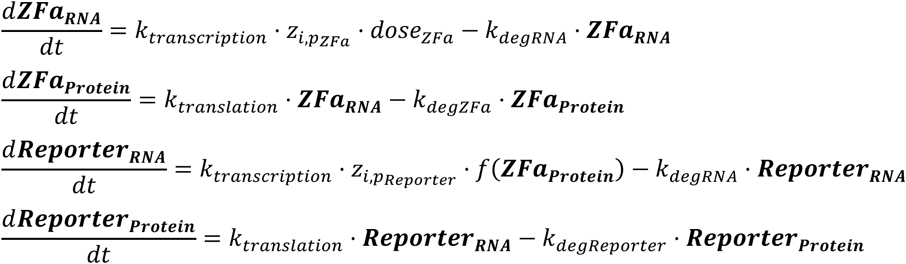

where *z* denotes the intracellular and intercellular variation, using values for the *i*^th^ cell and *p*^th^ plasmid. The model was run by iterating through each cell in the population (over a 42 h simulated duration corresponding to the experimental duration), and the population mean average was calculated.

In experiments from which data were used to estimate parameters, a ZF1a dose response with x6-C promoter was always included as a fiducial marker for normalizing experiment-specific MEFLs to model-specific a.u. that would be consistent across simulations. Parameters were estimated from dose response data as follows. For each ZFa: data were normalized based on the within-experiment ZF1a data series; *m* was specified based on maximum observed (or expected) reporter expression; *b* was determined from the data point for leaky expression; and *w* was fit by minimizing the sum of squares error between experimental data and simulated population means. The experiments from which parameters were estimated used a consistent dose of reporter plasmid (200 ng per well of cell culture in a 24-well plate).

#### Standard model of transcription

Fig. 2 compares the COMET model with a standard model of transcription that uses a greater number of parameters^20^. Fractional activation *f* by a TF (*y*) with promoter affinity *w* and Hill cooperativity *n* for TF-DNA binding, at a promoter that has one binding site, exhibits leaky transcription *α*_0_, and can be maximally activated by the TF to an amount *α*, is represented as:

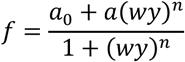

This formulation can be extended to other scenarios. For two TFs (*y*_1_ and *y*_2_) with respective maximal activation *α*_1_ and *α*_2_, a combined activation *α*_12_, and TF cooperativity *ρ* for RNAP recruitment, at a promoter with one site per TF, the formulation is:

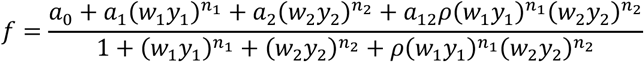

If in this scenario both TFs are the same (*i.e.*, one TF species can bind up to two sites), and additionally if maximal activation is 100% (*i.e.*, *α* = 1), this simplifies to:

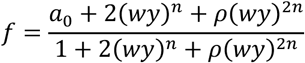

In a scenario without Hill cooperativity for TF-DNA binding (*i.e.*, *n* = 1) and without TF cooperativity (*i.e.*, *ρ* = 1), this further simplifies to:

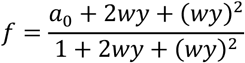

We extend the above case to any number of binding sites. Adding sites could affect *ρ* for each term in the numerator and denominator, but for simplicity we constrain the possible values by assuming all *ρ* = 1. This assumption is applied in the lower plots of the first and second landscapes in **Fig. 2c**. Examples are shown below for three, four, five, and six binding sites. Coefficients are derived using Pascal’s triangle:

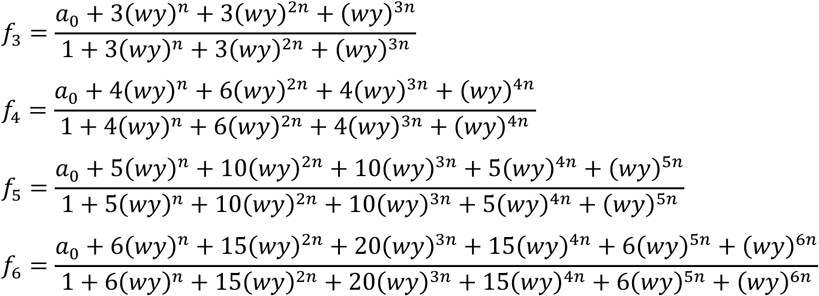

For the third and fourth landscapes in **Fig. 2c**, *m* values for spaced and compact promoters were substituted for *α* in each term of the numerator and denominator. As an example, the equation for three sites is:

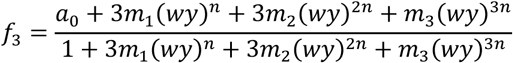

Since *m* values can exceed 1, *f* no longer formally represents *fractional* activation that is traditionally defined with the range of zero to one. This interpretational note also applies to *f* in the COMET model.

To investigate modes of transcriptional regulation, independent of the effects cell heterogeneity, the plots in **Fig. 2c,d** depict homogeneous (*i.e.*, one-cell) expression. The fits shown as lines in **Fig. 2a** depict mean averages calculated from a population of simulated cells. To compare the most salient features of each landscape in **Fig. 2c**, simulations were conducted using the parameter values below, and outcomes were scaled for a maximum attainable value of 1 within each model.

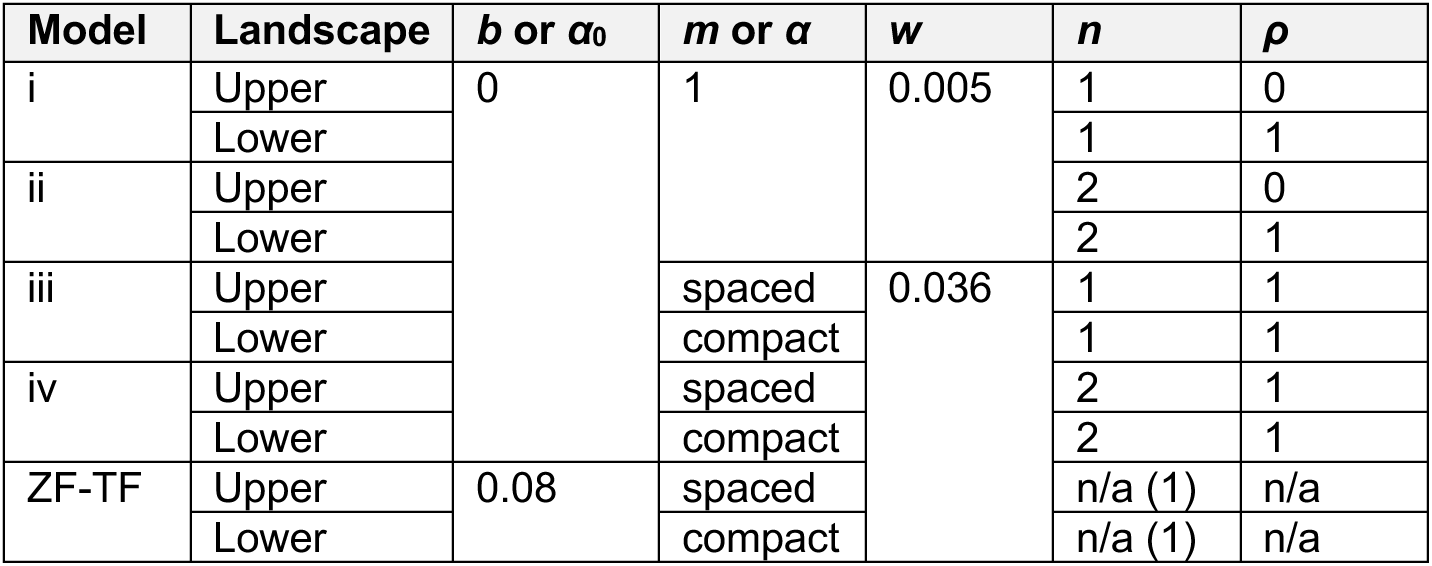

#### Mechanistic model of transcriptional regulation by ZFa

The mechanistic model depicted in **Supplementary Fig. 8d**, which incorporates a more detailed description of RNAPII recruitment, was developed as follows. To assess whether the concise formulation for *f* in COMET was consistent with a more detailed representation of gene regulation, we developed a model that represents interactions between molecular components—including those with tunable properties (ZFa, ZFi, and free AD, and for which doses can be specified) and those without tunable properties (RNAP and single-site reporter DNA). Here, the RNAP variable broadly represents the ensemble of factors that are recruited to initiate transcription, and this variable can bind to TF or AD. For simplicity, in this model, reporter DNA has one site that can be either unoccupied or occupied by a TF or a ZFi, and there is no leaky transcription.

The following initial conditions and parameters were used for the ODEs, which were run to steady state. Here, the goal was not to estimate parameter values. Therefore, the parameter values are based not on specific intracellular concentrations or rate constants, but rather those that we observed to produce steady-state trends that were interpretable and resembled the experimentally measured ZF1a dose response.

Initial values of variables; these depend on the condition specified for each plot in **Supplementary Fig. 8e,f**.

- Reporter DNA: 10 units
- RNAP: 200 units
- TF: dose response ranging from 0 to 200 units
- ZFi: 0 units; 200 if present at a constant amount; or a dose response of [5, 10, 20, 50, 100, 200]
- AD: 0 units; 200 if present at a constant amount; or a dose response of [5, 10, 20, 50, 100, 200]

Parameters for the reactions in **Supplementary Fig. 8de,f**.

- Association of TF (or ZFi) and DNA: *k*_a_ = 1 a.u.
- Dissociation of TF (or ZFi) and DNA: *k*_d_ = 100 a.u.; or varied as [500, 200, 100, 50, 20]
- Association of TF (or AD) and RNAP: *k*_f_ = 1 a.u.
- Dissociation of TF (or AD) and RNAP: *k*_r_ = 20 a.u.; or varied as [100, 50, 20, 10, 5]

The DNA.TF.RNAP variable was treated as a proxy for reporter expression. Metrics for TF interactions are *k*_a_/*k*_d_ for strength of TF-DNA interactions, and *k*_f_/*k*_r_ for the strength of RNAP recruitment. We observed that:

- The simulations qualitatively resembled experimental dose response trends.
- Increasing *k*_f_, *k*_a_, or TF dose led to more DNA.TF.RNAP (within a typical TF dose range). At values far above this range, or with excess non-productive components such as free AD, the dose response became non-monotonic due to non-productive sequestration of components.
- The effect of ZFi was to decrease DNA.TF.RNAP by occupying the DNA non-productively.
- Depending on components’ initial values, ZFa dose responses differed in two key features: the maximum attainable value and the slope, which are captured by *m* and *w* respectively in the concise model.

#### Mechanistic model of transcriptional regulation by ZFi (inhibitors)

The model used to generate predictions presented in **Fig. 5d,e** was developed as follows. Within the COMET framework, a competitive inhibitor is represented as:

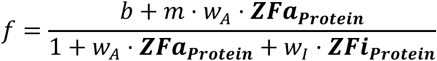

where *m* and *w_A_* correspond to the ZFa, and *w_I_* corresponds to the inhibitor. However, the observed effect of the inhibitors (**Fig. 5**) was greater than that predicted by competitive inhibition alone. We found that outcomes with ZFi-DsRed or a spaced promoter could be explained by also accounting for a decrease in *effective cooperativity* of the promoter. Removal of cooperativity from a multi-site promoter is a complex process involving an ensemble of promoter states within and between cells. For simplicity, we represent this as a non-mechanistic heuristic function that depends upon the amounts and properties of both the ZFa and the ZFi. The value *m* is replaced by a ramp-down function from baseline cooperativity without inhibitor to no cooperativity at a high amount of inhibitor:

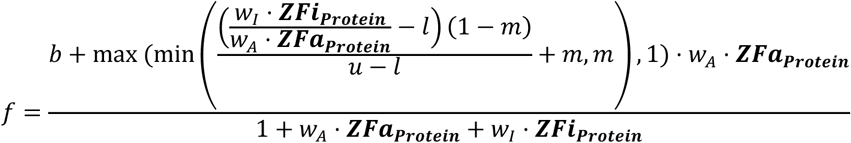

where *l* and *u* are empirically determined values for the weight-normalized ratio of inhibitor to activator at which the ramp-down from *m* to 1 begins and ends, respectively.

We found that compared to ZFi, the bulky ZFi-DsRed is a more effective inhibitor. Multiplying its weight by a factor of four improved the fit to data, and ramp-down parameters are adjusted accordingly to maintain the shape profile:

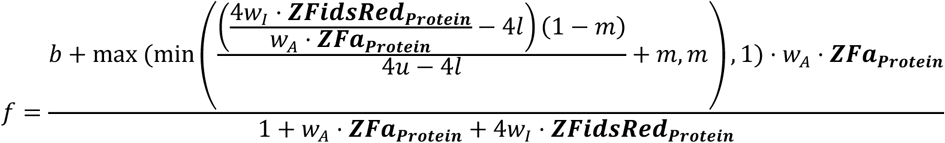

Ramp-downs in **Fig. 5d** use these values:

| Inhibitor | Promoter | $l$ | $u$ |
| --- | --- | --- | --- |
| ZFi | x6-compact | n/a | n/a |
|  | x6-spaced | 0 | 1.5 |
| ZFi-DsRed | x6-compact | 0.5 | 2 |
|  | x6-spaced | 0 | 1 |

For the inhibitor dose responses in **Fig. 5d**, cooperativity was more readily abolished with ZFi-DsRed than ZFi, and with a spaced promoter than a compact one. However, cooperativity was maintained with ZFi and a compact promoter, and this effect held across ZF1i mutants and doses reported in **Fig. 5e**.

#### Model of transcriptional AND gates

As depicted in **Fig. 8d**, we used the standard model from **Fig. 2** to investigate properties of an AND gate. For simplicity, leaky transcription (*a*_0_) is set to zero, and Hill coefficients (*n*_1_ and *n*_2_) are set to one. **Fig. 8d and Supplementary Fig. 14c** show four variations that differ in whether each TF’s maximal activation (*a*_1_ and *a*_2_) is less than or equal to the maximum activation with both present (*a*_12_ = 1), and synergy (*ρ*) is present or absent.

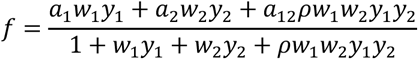

TFs were assigned identical properties, such that landscapes were symmetric about the dose response diagonal. Simulations used the homogeneous model and the following parameter values:

| Model | $a_1$ and $a_2$ | $a_{12}$ | $w_1$ and $w_2$ | $\rho$ |
| --- | --- | --- | --- | --- |
| i | 0.2 | 1 | 0.004 | 0 |
| ii | 1 | 1 |  | 0 |
| iii | 0.2 | 1 |  | 5 |
| iv | 1 | 1 |  | 5 |

In **Fig. 8d**, TF dose responses span 0 to 200 ng of plasmid, and target gene expression is linearly scaled to a maximum possible value of 1. Comparison between experiments and simulations shows that the hybrid COMET promoter exhibits hybrid cooperative activity—resembling x3-S with either ZFa individually and x6-C if both ZFa are present in sufficient amounts.

To explain this hybrid effect, we consider a scenario in which a ZFa induces transcription at a x6-C promoter for the reporter:

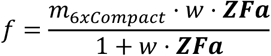

Hypothetically, if the pool of ZFa protein in a cell could be partitioned into two sub-pools of equal concentration, each with access to a distinct set of three alternating sites on the reporter promoter, then if only one sub-pool were active the promoter activity would decrease to:

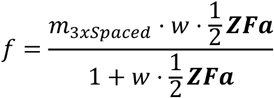

If the sub-pools differed in properties that affected *m* and *w*, then they could be treated as distinct TFs:

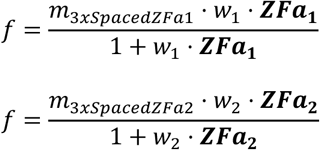

An inhibitor for either ZFa would act specifically on the corresponding binding sites, such that maximal inhibition would require inhibitor species that tile both sets of sites.

In the limit of high doses of both ZFa, the contribution of each individually to the total activation becomes:

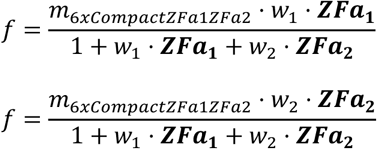

Together, these contributions sum to:

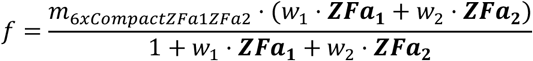

If both ZFa are identical, this expression becomes identical to the original expression.

Predictions made using the two-input AND gate model also guide interpretation of the circuits in **Fig. 8e,f**. In **Fig. 8e**, the three-input AND gate uses a similar principle as the two-input AND gate: promoter activity is effectively x2-S with each ZFa individually, and it transitions to x6-C if all three ZFa are present. In **Fig. 8f**, ZFa and ZFi-DsRed modulate the effective number (x0, x1, x3, x6) and spacing (S = spaced, C = compact) of binding sites, and whether there is competitive inhibition (Y = yes, N = no, Y/N = yes for some sites and no for others). The experimental outcomes align with the expectations below:

|  |  |  |  |  |  |  |  |  |  |  |  |  |  |  |  |  |
| --- | --- | --- | --- | --- | --- | --- | --- | --- | --- | --- | --- | --- | --- | --- | --- | --- |
| **ZF2a** | – |  |  |  | + |  |  |  | – |  |  |  | + |  |  |  |
| **ZF3a** | – |  |  |  | – |  |  |  | + |  |  |  | + |  |  |  |
| **ZF2i-DsRed** | – | + | – | + | – | + | – | + | – | + | – | + | – | + | – | + |
| **ZF3i-DsRed** | – |  | + |  | – |  | + |  | – |  | + |  | – |  | + |  |
| **Effective # of sites** | x0 | x0 | x0 | x0 | x3 | x1 | x3 | x1 | x3 | x3 | x1 | x1 | x6 | x3 | x3 | x1 |
| **Effective spacing** | n/a | n/a | n/a | n/a | S | n/a | S | n/a | S | S | n/a | n/a | C | S | S | n/a |
| **Competitive inhibition** | n/a | n/a | n/a | n/a | N | Y | N | Y | N | N | Y | Y | N | Y/N | N/Y | Y/Y |

Importantly, among the 16 combinations of the four TF inputs, only the target combination exhibits x6-C behavior. The resulting cooperativity leads to much higher reporter expression than other combinations with x3-S or x1 behavior.

## STATISTICAL ANALYSIS

Statistical details for each experiment are in the figure legends. Unless otherwise stated, there are three independent biologic replicates for each condition. The data shown reflect the mean across these biologic replicates of the mean fluorescence intensity (MFI) of approximately 2,000–3,000 single, transfected cells. Error bars represent the standard error of the mean (SEM). For main figures with heat maps, data are also shown in the corresponding supplemental figure as a bar graph with the mean and S.E.M.

ANOVA tests were performed using the Data Analysis Toolpak in Microsoft Excel. Tukey’s HSD tests were performed with *α* = 0.05. Pairwise comparisons were made using a one-tailed Welch’s *t*-test, which is a version of Student’s *t*-test in which the variance between samples is treated as not necessarily equal. The comparisons involved reporter only vs. reporter + ZFa in **Fig. 1, Fig. 3**; inhibited vs. uninhibited, or more inhibited vs. less inhibited, in **Fig. 5**; no binding sites vs. one binding site in **Supplementary Fig. 10**; DMSO vs. rapamycin in **Fig. 7**; and summed individual cases vs. co-expression in **Fig. 8**. For each comparison, the null hypothesis was that two samples were equal, and the alternative was that the latter was greater. The threshold for significance was set at 0.05. To decrease the false discovery rate, the Benjamini-Hochberg (BH) procedure was applied to each set of tests per figure panel; in all tests, after the BH procedure, the null hypothesis was rejected for *p*-values < 0.05. The outcome of each statistical test is indicated in the figure captions.

## CODE AVAILABILITY

All code is provided in MATLAB format in the supplementary material.

