## Supplementary material for "COMET: A toolkit for composing customizable genetic programs in mammalian cells": Software files archive: generate_TXF_distribution.m

```
%Donahue PS, Draut JW, Muldoon JJ, Edelstein HI, Bagheri N, & Leonard JN.
%COMET: A toolkit for composing customizable genetic programs in mammalian cells.

%File for generating a matrix representing a heterogeneous population of
%cells transfected with one or more plasmids


function [Z, rmat] = generate_TXF_distribution(nc, np)


%Input arguments:
%    nc:   number of cells in population   (at least 1)
%    np:   number of different plasmids    (at least 1, and this file permits up to 8)
%
%Output arguments:
%    Z:    population heterogeneity matrix (dimensions nc x np)
%    rmat: correlation coefficient matrix  (dimensions np x np)

%Notes:
%    Run time increases with choice of nc and np
%    This study used nc = 200
%    An accompanying file is provided for nc = 200 and np = 6


%****************************%
%**** Specify parameters ****%
%****************************%


%parameters for distribution associated with transfection methodology;
%there are two modes (labeled 1 and 2).
%proportion of the population for each Gaussian
c1  = 0.4;
c2  = 1 - c1;
%means
mu1 = 1.95;
mu2 = 3.4;
%standard deviations
s1  = 0.3;
s2  = 0.6;

%Target correlation coefficient for log-transformed
%This value can be empirically chosen to generate solutions for which the
%non-diagonal entries of the correlation coefficient matrix will fall
%within the range of acceptable values specified below.
r = 0.85;

%range of acceptable correlation coefficients for untransformed
%We chose values close to r = 0.8, which is the target correlation for
%the transfection methodology
rmin = 0.765;
rmax = 0.835;


%*****************************%
%**** Generate population ****%
%*****************************%


%initialize counter for number of attempts to generate a suitable matrix
counter = 0;


%the elseif statements below are for different choices of np.
%within each statement, candidate matrices are generally based on the
%specified rameters, and matrices are evaluated by whether the non-diagonal
%entries of the correlation coefficient matrix are within acceptable bounds

%one plasmid
if np == 1
    
    %populate
    Z1 = mu1 + s1 * mvnrnd(zeros(1, np), 1, c1 * nc);
    Z2 = mu2 + s2 * mvnrnd(zeros(1, np), 1, c2 * nc);
    
    %update counter
    counter = counter + 1;
    
    %un-log transform and stack Z1 and Z2
    Z = [10.^Z1; 10.^Z2];
    
    %correlation coefficient matrix (not relevant for the one-plasmid case)
    rmat = corrcoef(Z);
    
%two plasmids
elseif np == 2
    
    %Initialize
    rmat = zeros(np, np);
    
    %while conditions are not yet met
    while rmat(2, 1) <= rmin || rmat(2, 1) >= rmax
        
        %populate
        Z1 = mu1 + s1 * mvnrnd(zeros(1, np), [1, r; r, 1], c1 * nc);
        Z2 = mu2 + s2 * mvnrnd(zeros(1, np), [1, r; r, 1], c2 * nc);
        
        %update counter
        counter = counter + 1;
        
        %un-log transform and stack Z1 and Z2
        Z = [10.^Z1; 10.^Z2];
        
        %correlation coefficient matrix
        rmat = corrcoef(Z);
    end
    
%three plasmids
elseif np == 3
    
    %Initialize
    rmat = zeros(np, np);
    
    %while conditions are not yet met
    while      (rmat(2, 1) <= rmin || rmat(2, 1) >= rmax) ...
            || (rmat(3, 1) <= rmin || rmat(3, 1) >= rmax) ...
            || (rmat(3, 2) <= rmin || rmat(3, 2) >= rmax)
        
        %populate
        Z1 = mu1 + s1 * mvnrnd(zeros(1, np), [
            1, r, r;
            r, 1, r;
            r, r, 1], c1 * nc);
        Z2 = mu2 + s2 * mvnrnd(zeros(1, np), [
            1, r, r;
            r, 1, r;
            r, r, 1], c2 * nc);
        
        %update counter
        counter = counter + 1;
        
        %un-log transform and stack Z1 and Z2
        Z = [10.^Z1; 10.^Z2];
        
        %correlation coefficient matrix
        rmat = corrcoef(Z);
    end
    
%four plasmids
elseif np == 4
    
    %Initialize
    rmat = zeros(np, np);
    
    %while conditions are not yet met
    while      (rmat(2, 1) <= rmin || rmat(2, 1) >= rmax) ...
            || (rmat(3, 1) <= rmin || rmat(3, 1) >= rmax) ...
            || (rmat(4, 1) <= rmin || rmat(4, 1) >= rmax) ...
            || (rmat(3, 2) <= rmin || rmat(3, 2) >= rmax) ...
            || (rmat(4, 2) <= rmin || rmat(4, 2) >= rmax) ...
            || (rmat(4, 3) <= rmin || rmat(4, 3) >= rmax)
        
        %update counter
        counter = counter + 1;
        
        Z1 = mu1 + s1 * mvnrnd(zeros(1, np), [
            1, r, r, r;
            r, 1, r, r;
            r, r, 1, r;
            r, r, r, 1], c1 * nc);
        Z2 = mu2 + s2 * mvnrnd(zeros(1, np), [
            1, r, r, r;
            r, 1, r, r;
            r, r, 1, r;
            r, r, r, 1], c2 * nc);
        
        %un-log transform and stack Z1 and Z2
        Z = [10.^Z1; 10.^Z2];
        
        %correlation coefficient matrix
        rmat = corrcoef(Z);
    end
    
%five plasmids
elseif np == 5
    
    %Initialize
    rmat = zeros(np, np);
    
    %while conditions are not yet met
    while      (rmat(2, 1) <= rmin || rmat(2, 1) >= rmax) ...
            || (rmat(3, 1) <= rmin || rmat(3, 1) >= rmax) ...
            || (rmat(4, 1) <= rmin || rmat(4, 1) >= rmax) ...
            || (rmat(5, 1) <= rmin || rmat(5, 1) >= rmax) ...
            || (rmat(3, 2) <= rmin || rmat(3, 2) >= rmax) ...
            || (rmat(4, 2) <= rmin || rmat(4, 2) >= rmax) ...
            || (rmat(5, 2) <= rmin || rmat(5, 2) >= rmax) ...
            || (rmat(4, 3) <= rmin || rmat(4, 3) >= rmax) ...
            || (rmat(5, 3) <= rmin || rmat(5, 3) >= rmax) ...
            || (rmat(5, 4) <= rmin || rmat(5, 4) >= rmax)
        
        %update counter
        counter = counter + 1;
        
        %populate
        Z1 = mu1 + s1 * mvnrnd(zeros(1, np), [
            1, r, r, r, r;
            r, 1, r, r, r;
            r, r, 1, r, r;
            r, r, r, 1, r;
            r, r, r, r, 1], c1 * nc);
        Z2 = mu2 + s2 * mvnrnd(zeros(1, np), [
            1, r, r, r, r;
            r, 1, r, r, r;
            r, r, 1, r, r;
            r, r, r, 1, r;
            r, r, r, r, 1], c2 * nc);
        
        %un-log transform and stack Z1 and Z2
        Z = [10.^Z1; 10.^Z2];
        
        %correlation coefficient matrix
        rmat = corrcoef(Z);
    end

%six plasmids
elseif np == 6
    
    %Initialize
    rmat = zeros(np, np);
    
    %while conditions are not yet met
    while      (rmat(2, 1) <= rmin || rmat(2, 1) >= rmax) ...
            || (rmat(3, 1) <= rmin || rmat(3, 1) >= rmax) ...
            || (rmat(4, 1) <= rmin || rmat(4, 1) >= rmax) ...
            || (rmat(5, 1) <= rmin || rmat(5, 1) >= rmax) ...
            || (rmat(6, 1) <= rmin || rmat(6, 1) >= rmax) ...
            || (rmat(3, 2) <= rmin || rmat(3, 2) >= rmax) ...
            || (rmat(4, 2) <= rmin || rmat(4, 2) >= rmax) ...
            || (rmat(5, 2) <= rmin || rmat(5, 2) >= rmax) ...
            || (rmat(6, 2) <= rmin || rmat(6, 2) >= rmax) ...
            || (rmat(4, 3) <= rmin || rmat(4, 3) >= rmax) ...
            || (rmat(5, 3) <= rmin || rmat(5, 3) >= rmax) ...
            || (rmat(6, 3) <= rmin || rmat(6, 3) >= rmax) ...
            || (rmat(5, 4) <= rmin || rmat(5, 4) >= rmax) ...
            || (rmat(6, 4) <= rmin || rmat(6, 4) >= rmax) ...
            || (rmat(6, 5) <= rmin || rmat(6, 5) >= rmax)
        
        %update counter
        counter = counter + 1;
        
        %populate
        Z1 = mu1 + s1 * mvnrnd(zeros(1, np), [
            1, r, r, r, r, r;
            r, 1, r, r, r, r;
            r, r, 1, r, r, r;
            r, r, r, 1, r, r;
            r, r, r, r, 1, r;
            r, r, r, r, r, 1], c1 * nc);
        Z2 = mu2 + s2 * mvnrnd(zeros(1, np), [
            1, r, r, r, r, r;
            r, 1, r, r, r, r;
            r, r, 1, r, r, r;
            r, r, r, 1, r, r;
            r, r, r, r, 1, r;
            r, r, r, r, r, 1], c2 * nc);
        
        %un-log transform and stack Z1 and Z2
        Z = [10.^Z1; 10.^Z2];
        
        %correlation coefficient matrix
        rmat = corrcoef(Z);
    end

%seven plasmids
elseif np == 7
    
    %Initialize
    rmat = zeros(np, np);
    
    %while conditions are not yet met
    while      (rmat(2, 1) <= rmin || rmat(2, 1) >= rmax) ...
            || (rmat(3, 1) <= rmin || rmat(3, 1) >= rmax) ...
            || (rmat(4, 1) <= rmin || rmat(4, 1) >= rmax) ...
            || (rmat(5, 1) <= rmin || rmat(5, 1) >= rmax) ...
            || (rmat(6, 1) <= rmin || rmat(6, 1) >= rmax) ...
            || (rmat(7, 1) <= rmin || rmat(7, 1) >= rmax) ...
            || (rmat(3, 2) <= rmin || rmat(3, 2) >= rmax) ...
            || (rmat(4, 2) <= rmin || rmat(4, 2) >= rmax) ...
            || (rmat(5, 2) <= rmin || rmat(5, 2) >= rmax) ...
            || (rmat(6, 2) <= rmin || rmat(6, 2) >= rmax) ...
            || (rmat(7, 2) <= rmin || rmat(7, 2) >= rmax) ...
            || (rmat(4, 3) <= rmin || rmat(4, 3) >= rmax) ...
            || (rmat(5, 3) <= rmin || rmat(5, 3) >= rmax) ...
            || (rmat(6, 3) <= rmin || rmat(6, 3) >= rmax) ...
            || (rmat(7, 3) <= rmin || rmat(7, 3) >= rmax) ...
            || (rmat(5, 4) <= rmin || rmat(5, 4) >= rmax) ...
            || (rmat(6, 4) <= rmin || rmat(6, 4) >= rmax) ...
            || (rmat(7, 4) <= rmin || rmat(7, 4) >= rmax) ...
            || (rmat(6, 5) <= rmin || rmat(6, 5) >= rmax) ...
            || (rmat(7, 5) <= rmin || rmat(7, 5) >= rmax) ...
            || (rmat(7, 6) <= rmin || rmat(7, 6) >= rmax)
        
        %update counter
        counter = counter + 1;
        
        %populate
        Z1 = mu1 + s1 * mvnrnd(zeros(1, np), [
            1, r, r, r, r, r, r;
            r, 1, r, r, r, r, r;
            r, r, 1, r, r, r, r;
            r, r, r, 1, r, r, r;
            r, r, r, r, 1, r, r;
            r, r, r, r, r, 1, r;
            r, r, r, r, r, r, 1], c1 * nc);
        Z2 = mu2 + s2 * mvnrnd(zeros(1, np), [
            1, r, r, r, r, r, r;
            r, 1, r, r, r, r, r;
            r, r, 1, r, r, r, r;
            r, r, r, 1, r, r, r;
            r, r, r, r, 1, r, r;
            r, r, r, r, r, 1, r;
            r, r, r, r, r, r, 1], c2 * nc);
        
        %un-log transform and stack Z1 and Z2
        Z = [10.^Z1; 10.^Z2];
        
        %correlation coefficient matrix
        rmat = corrcoef(Z);
    end

%eight plasmids
elseif np == 8
    
    %Initialize
    rmat = zeros(np, np);
    
    %while conditions are not yet met
    while      (rmat(2, 1) <= rmin || rmat(2, 1) >= rmax) ...
            || (rmat(3, 1) <= rmin || rmat(3, 1) >= rmax) ...
            || (rmat(4, 1) <= rmin || rmat(4, 1) >= rmax) ...
            || (rmat(5, 1) <= rmin || rmat(5, 1) >= rmax) ...
            || (rmat(6, 1) <= rmin || rmat(6, 1) >= rmax) ...
            || (rmat(7, 1) <= rmin || rmat(7, 1) >= rmax) ...
            || (rmat(8, 1) <= rmin || rmat(8, 1) >= rmax) ...
            || (rmat(3, 2) <= rmin || rmat(3, 2) >= rmax) ...
            || (rmat(4, 2) <= rmin || rmat(4, 2) >= rmax) ...
            || (rmat(5, 2) <= rmin || rmat(5, 2) >= rmax) ...
            || (rmat(6, 2) <= rmin || rmat(6, 2) >= rmax) ...
            || (rmat(7, 2) <= rmin || rmat(7, 2) >= rmax) ...
            || (rmat(8, 2) <= rmin || rmat(8, 2) >= rmax) ...
            || (rmat(4, 3) <= rmin || rmat(4, 3) >= rmax) ...
            || (rmat(5, 3) <= rmin || rmat(5, 3) >= rmax) ...
            || (rmat(6, 3) <= rmin || rmat(6, 3) >= rmax) ...
            || (rmat(7, 3) <= rmin || rmat(7, 3) >= rmax) ...
            || (rmat(8, 3) <= rmin || rmat(8, 3) >= rmax) ...
            || (rmat(5, 4) <= rmin || rmat(5, 4) >= rmax) ...
            || (rmat(6, 4) <= rmin || rmat(6, 4) >= rmax) ...
            || (rmat(7, 4) <= rmin || rmat(7, 4) >= rmax) ...
            || (rmat(8, 4) <= rmin || rmat(8, 4) >= rmax) ...
            || (rmat(6, 5) <= rmin || rmat(6, 5) >= rmax) ...
            || (rmat(7, 5) <= rmin || rmat(7, 5) >= rmax) ...
            || (rmat(8, 5) <= rmin || rmat(8, 5) >= rmax) ...
            || (rmat(7, 6) <= rmin || rmat(7, 6) >= rmax) ...
            || (rmat(8, 6) <= rmin || rmat(8, 6) >= rmax) ...
            || (rmat(8, 7) <= rmin || rmat(8, 7) >= rmax)
        
        %update counter
        counter = counter + 1;
        
        %populate
        Z1 = mu1 + s1 * mvnrnd(zeros(1, np), [
            1, r, r, r, r, r, r, r;
            r, 1, r, r, r, r, r, r;
            r, r, 1, r, r, r, r, r;
            r, r, r, 1, r, r, r, r;
            r, r, r, r, 1, r, r, r;
            r, r, r, r, r, 1, r, r;
            r, r, r, r, r, r, 1, r;
            r, r, r, r, r, r, r, 1], c1 * nc);
        Z2 = mu2 + s2 * mvnrnd(zeros(1, np), [
            1, r, r, r, r, r, r, r;
            r, 1, r, r, r, r, r, r;
            r, r, 1, r, r, r, r, r;
            r, r, r, 1, r, r, r, r;
            r, r, r, r, 1, r, r, r;
            r, r, r, r, r, 1, r, r;
            r, r, r, r, r, r, 1, r;
            r, r, r, r, r, r, r, 1], c2 * nc);
        
        %un-log transform and stack Z1 and Z2
        Z = [10.^Z1; 10.^Z2];
        
        %correlation coefficient matrix
        rmat = corrcoef(Z);
    end
end


%for each plasmid
for p = 1:np
    %normalize such that the mean of each column equals one
    Z(:, p) = Z(:, p) / mean(Z(:, p));
end


%display counter value
disp(['counter: ', num2str(counter)]);


end
```
