## Supplementary material for "COMET: A toolkit for composing customizable genetic programs in mammalian cells": Software files archive: model_ZFa.m

```
%Donahue PS, Draut JW, Muldoon JJ, Edelstein HI, Bagheri N, & Leonard JN.
%COMET: A toolkit for composing customizable genetic programs in mammalian cells.

%Simulation of ZFa-inducible reporter expression.


function sim = model_ZFa(doseZFa, m, w, b, Z)


%Input arguments:
%    doseZFa: ZFa plasmid (ng)
%    m:       maximum activation parameter
%    w:       steepness parameter
%    b:       background (TF-independent transcription) parameter
%    Z:       one row from the population matrix (not the whole 2-D matrix)

%Output argument:
%    sim: simulation (dimensions: number of time points x number of state variables)

%Notes:
%    Output is in model-specific a.u.

%Example for ZF1a and the c'th cell in a heterogeneous population:
%    sim = model_ZFa(50, 32.7, 0.036, 0.08, Z(c, :));

%Example for a homogeneous simulation:
%    sim = model_ZFa(50, 32.7, 0.036, 0.08, [1, 1]);


%*****************%
%**** Specify ****%
%*****************%


%timecourse (h)
T = 0:42; %the first number is zero, and the second number corresponds to the time of measurement post-transfection

%general parameters
ktxCMV   = 1;     %transcription                         defined as 1 a.u.
ktxZF    = 9;     %transcription of regulated promoters  fitted
kdegR    = 2.7;   %degradation of RNA                    assumed half-life of 15 min.
ktl      = 1;     %translation                           defined as 1 a.u.
kdegZFP  = 0.35;  %degradation of ZF-TF                  assumed half-life of 2 h.
kdegRepP = 0.029; %degradation of reporter protein       half-life ~1 day.

%initial values (a.u.)
IV = zeros(4, 1);

%intercellular variation for the ZFa
za  = Z(1);

%intercellular variation for the reporter
zrep = Z(2);


%***************%
%**** Model ****%
%***************%


%simulate ODEs
[~, sim] = ode15s(@(t, y, options)[ 
    
    %Y1 ZFa RNA
      ktxCMV   * za * doseZFa                                          ... %constitutive transcription
    - kdegR    * y(1)                                                      %RNA degradation
    
    %Y2 ZFa protein
      ktl      * y(1)                                                  ... %translation
    - kdegZFP  * y(2)                                                      %protein degradation
    
    %Y3 Reporter RNA
      ktxZF    * zrep * (b + m * w * y(2)) ./ (1 + w * y(2))           ... %inducible transcription
    - kdegR    * y(3)                                                      %RNA degradation
    
    %Y4 Reporter protein
      ktl      * y(3)                                                  ... %translation
    - kdegRepP * y(4)                                                      %protein degradation
    
    ], T, IV);


end
```
