## Supplementary material for "COMET: A toolkit for composing customizable genetic programs in mammalian cells": Software files archive: COMET_README.rtf


Description of files: 

	•	generate_TXF_distribution.m: File for generating a matrix representing a heterogeneous population of cells transfected with one or more plasmids 

	•	Z_TXF.mat: An output from generate_TXF_distribution.m for 200 cells transfected with up to six plasmids. 

	•	model_ZFa.m: Simulation of ZFa-inducible reporter expression. 

	•	model_ZFa_ZFi_competitive.m: Simulation of ZFa-inducible and ZFi-inhibitable reporter expression. The ZFi effect is represented as purely competitive without affecting cooperative RNAPII recruitment. 

	•	model_ZFa_ZFi_dual.m: Simulation of ZFa-inducible and ZFi-inhibitable reporter expression. The ZFi effect is represented as both competitive and affecting cooperative RNAPII recruitment. 

	•	model_ZFa_ZFiDsRed_dual.m: Simulation of ZFa-inducible and ZFi-DsRed-inhibitable reporter expression. The ZFi-DsRed effect is represented as both competitive and affecting cooperative RNAPII recruitment. 

1. System requirements

The code can be run using Matlab (https://www.mathworks.com/products/matlab.html) and run on any operating system that supports it. Code was developed and tested on macOS Sierra.

2. Installation guide

No specific installation is required other than for Matlab.

3. Demo

Files can be run in the Matlab command line with the input arguments described and examples provided in the comments section at the start of each file, and they will produce the output arguments as described.

The expected run time for each function, other than generate_TXF_distribution.m, is << 1 second on a standard desktop computer.

4. Instructions for use

Parameter values for new TFs and promoters can be estimated as described in the Supplementary Information.
