## Supplementary Information for "COMET: A toolkit for composing customizable genetic programs in mammalian cells"

Patrick S. Donahue[1,2,3], Joseph W. Draut[1,8], Joseph J. Muldoon[1,2,8], Hailey I. Edelstein[1,8], Neda Bagheri[1,2,4,5,6,7], & Joshua N. Leonard[1,2,4,5,6,*]

[1]Department of Chemical and Biological Engineering, Northwestern University, Evanston, Illinois 60208, United States

[2]Interdisciplinary Biological Sciences Program, Northwestern University, Evanston, Illinois 60208, United States

[3]Medical Scientist Training Program, Northwestern University Feinberg School of Medicine, Chicago, Illinois 60611, United States

[4]Center for Synthetic Biology, Northwestern University, Evanston, Illinois 60208, United States

[5]Chemistry of Life Processes Institute, Northwestern University, Evanston, Illinois 60208, United States

[6]Member, Robert H. Lurie Comprehensive Cancer Center, Northwestern University, Evanston, Illinois 60208, United States

[7]Biology and Chemical Engineering, University of Washington, Seattle, Washington 98195, United States

[8]These authors contributed equally to this work

**Contents:**

Supplementary Figures 1–16

Supplementary Tables 1–18[*]

Supplementary Notes 1–3

[*] This document contains legends for all supplementary tables and the contents for Supplementary Tables 9 and 10; the remaining supplementary tables are provided as Microsoft Excel Worksheets.

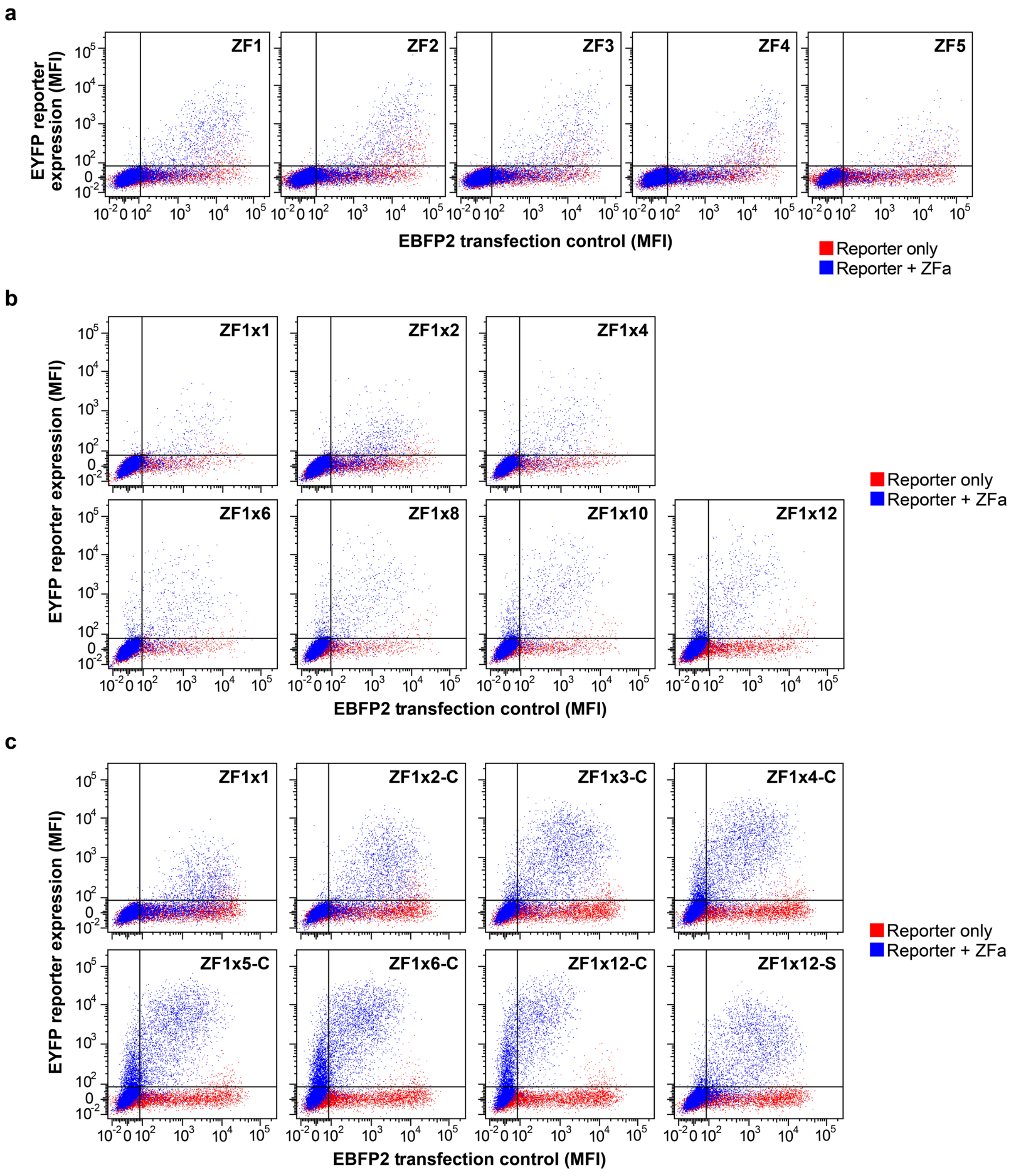


**Supplementary Fig. 1 Effects of the choice of ZF and the number and spacing of ZF binding sites.** (a) Representative flow cytometry plots from **Fig. 1b**. The joint distribution shows that inducible EYFP expression is higher in more highly transfected cells (i.e., those with more EBFP2 expression). Constitutively expressed EBFP2 is the transfection control. (b) Representative flow cytometry plots from **Fig. 1c**. (c) Representative flow cytometry plots from **Fig. 1e**. Squelching is suggested in conditions with strong COMET promoters (i.e., higher EYFP expression), which exhibit lower EBPF2 fluorescence than do conditions with weaker COMET promoters.

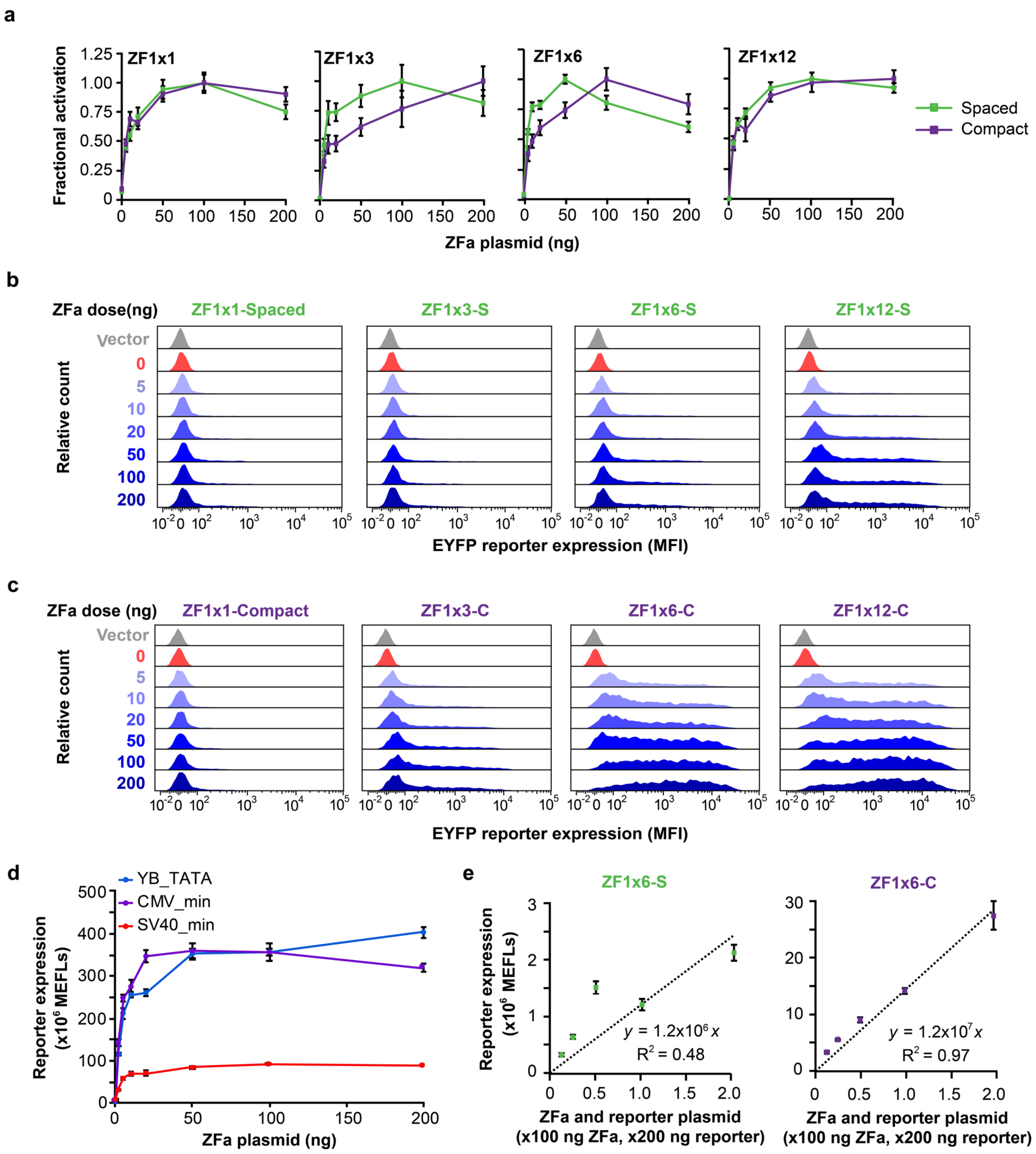
a
Fractional activation
ZF1x1
ZF1x3
ZF1x6
ZF1x12
Spaced
Compact
ZFa plasmid (ng)
b
ZFa dose(ng)
ZF1x1-Spaced
ZF1x3-S
ZF1x6-S
ZF1x12-S
Vector
0
5
10
20
50
100
200
Relative count
EYFP reporter expression (MFI)
c
ZFa dose (ng)
ZF1x1-Compact
ZF1x3-C
ZF1x6-C
ZF1x12-C
Vector
0
5
10
20
50
100
200
Relative count
EYFP reporter expression (MFI)
d
YB_TATA
CMV_min
SV40_min
Reporter expression (x10^6 MEFLs)
ZFa plasmid (ng)
e
ZF1x6-S
ZF1x6-C
Reporter expression (x10^6 MEFLs)
y = 1.2x10^6 x
R^2 = 0.48
y = 1.2x10^7 x
R^2 = 0.97
ZFa and reporter plasmid
(x100 ng ZFa, x200 ng reporter)

**Supplementary Fig. 2 Differences between spaced and compact promoters.** (a) Fractional activation for dose response profiles in **Fig. 2a**. Fractional activation was determined by dividing each data point by the maximum reporter expression induced by the ZFa on a given reporter. In several conditions, notably ZF1x6-S, excess ZFa (above 50 ng plasmid) resulted in a decrease in reporter expression. In this case, we hypothesize that unbound ZFa competes with bound ZFa for endogenous cofactors required for transcription. (b) Representative flow cytometry histograms for experiments with spaced reporters in **Fig. 2a**. Data were gated on single, transfected cells (**Supplementary Fig. 15**). Reporter expression increases with ZFa dose and number of binding sites. (c) Representative flow cytometry histograms from experiments with compact reporters in **Fig. 2a**. Data were gated on single, transfected cells. Reporter expression increases with ZFa dose and number of binding sites. Compared to the case of spaced promoters (panel b**)**, cases with compact promoters exhibit a greater fraction of cells that are distinguishably ON, i.e., expressing more EYFP than cells without ZFa. (d) Investigating different minimal promoters with COMET. ZF1a dose responses were conducted using ZF1x6-C promoters with either the YB_TATA, CMV, or SV40 minimal promoters. The SV40 minimal promoter produced low levels of gene expression, while the YB_TATA minimal promoter conferred a maximal gene expression level similar to that of the CMV minimal promoter. Although the CMV minimal promoter was more responsive at lower levels of ZFa expression, this promoter also had higher leaky gene expression (in the absence of ZFa) than did the YB_TATA minimal promoter (quantified in **Supplementary Table 10** with fitted parameters). Thus, the YB_TATA minimal promoter resulted in higher fold inductions than did the CMV minimal promoter (approximately 220-fold for YB_TATA, as compared to 60-fold for CMV_min) without sacrificing maximal gene expression. (e) Investigating maximal inducible EYFP expression. Cells were transfected with ZF1a plasmid and with reporter plasmid containing a ZF1x6-S (*left*) or ZF1x6-C (*right*) promoter. The ZFa plasmid and reporter plasmid were maintained at a ratio of 1:2 (ZFa:reporter) as the doses were scaled. On the x-axis, a value of 1 denotes a condition with 100 ng of ZF1a plasmid and 200 ng of reporter plasmid. In previous experiments, reporter expression typically plateaued at the level indicated by plasmid doses corresponding to 1 on the x-axis and could not be increased by the addition of more ZFa plasmid (**Supplementary Fig. 2a**). However, doubling the amount of *both* ZFa plasmid and reporter plasmid led to twice the reporter expression, which indicates the amount of plasmid was the limiting factor in gene expression as opposed to a downstream step such as translation. For the compact promoter, reporter expression scaled linearly with dose of these components (at a fixed ratio) (one-tailed permutation test $p = 0.001$), but no strong linear correlation was observed for the spaced promoter ($p = 0.10$). Error bars depict S.E.M.

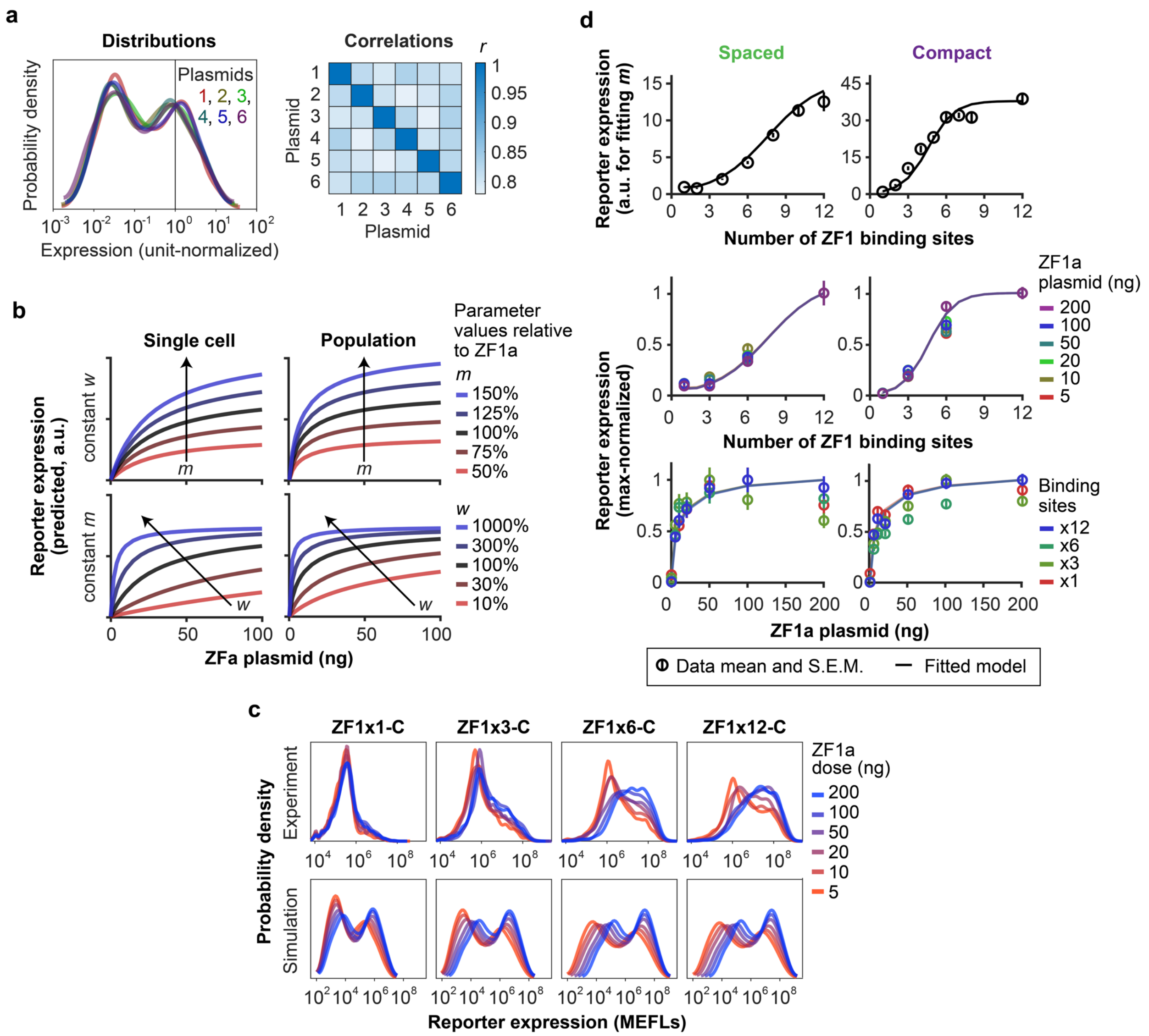

a
Distributions
Correlations
Plasmids
1, 2, 3,
4, 5, 6
Probability density
Expression (unit-normalized)
Plasmid
r
b
Single cell
Population
Parameter values relative to ZF1a
m
150%
125%
100%
75%
50%
w
1000%
300%
100%
30%
10%
constant w
constant m
Reporter expression (predicted, a.u.)
ZFa plasmid (ng)
c
ZF1x1-C
ZF1x3-C
ZF1x6-C
ZF1x12-C
ZF1a dose (ng)
200
100
50
20
10
5
Experiment
Simulation
Probability density
Reporter expression (MEFLs)
d
Spaced
Compact
Reporter expression (a.u. for fitting m)
Number of ZF1 binding sites
ZF1a plasmid (ng)
200
100
50
20
10
5
Reporter expression (max-normalized)
Binding sites
x12
x6
x3
x1
ZF1a plasmid (ng)
Data mean and S.E.M.
Fitted model

**Supplementary Fig. 3 A model for ZFa-regulated gene expression.** (a) A statistical model for cell heterogeneity is presented. Accounting for variation in gene expression enables more accurate explanations and predictions for genetic programs. An *in silico* population was generated for 200 cells and multiple plasmids using a previously described method[1]. Properties of simulated cell populations are summarized here. *Left:* marginal distributions from the model show intercellular variation in the relative expression of a constitutive gene matching expected distributions for single, transfected cells. Each distribution is normalized to a mean value of 1 arbitrary unit (a.u.) at the vertical line. *Right:* pairwise correlations capture intracellular variation, shown for the relative expression of six constitutively expressed genes on separate plasmids. (b) Advantages of using a population model. Simulated dose responses are shown for hypothetical ZFa with varying *m* or *w* parameters (rows). Outcomes are shown for a single cell and for the mean of a heterogenous population (columns), where a single cell refers to a cell expressing the mean amount of each component. The comparison shows how, for the same model parameter values, the presence of population-level heterogeneity can lead to greater observed reporter expression. As a result, fitting models to experimental data (which include population-level heterogeneity) using a standard homogeneous approach (which corresponds to the single cell case) can lead to poor estimates of model parameters. Approaches that account for cell heterogeneity mitigate this problem. Y-axes are linearly scaled. (c) Experimental (flow cytometry) and simulated distributions of reporter expression for different ZF1a doses and numbers of binding sites. The model captures the observed bimodal log-Gaussian distributions, and the fact that at increasing ZFa doses, the probability density shifts from the lower mode to the upper mode. Simulated reporter expression is presented in internally consistent model-a.u. which are linearly scaled to align with experimental MEFLs. While MEFLs are absolute units, the magnitude of reporter induction varies between experiments, and this is why simulated distributions are scaled. (d) Effects of the number and spacing of binding sites and the ZFa dose. Cross-sections from experimental data (circles) and simulated landscapes in **Fig. 2a** were normalized to the maximum value within each cross-section. We observed that these normalized curves followed characteristic profiles. Lines show end-point (long-time) mean reporter expression from dynamical population simulations. *Upper:* reporter expression varies sigmoidally with the number of binding sites (described in **Online Methods)**. Spaced and compact promoters have distinct profiles. *Middle:* the scaled sigmoidal shape of the response to the number of binding sites holds across ZFa plasmid doses. *Lower:* the scaled concave shape of the response to ZFa dose holds across the number and spacing of binding sites.

a

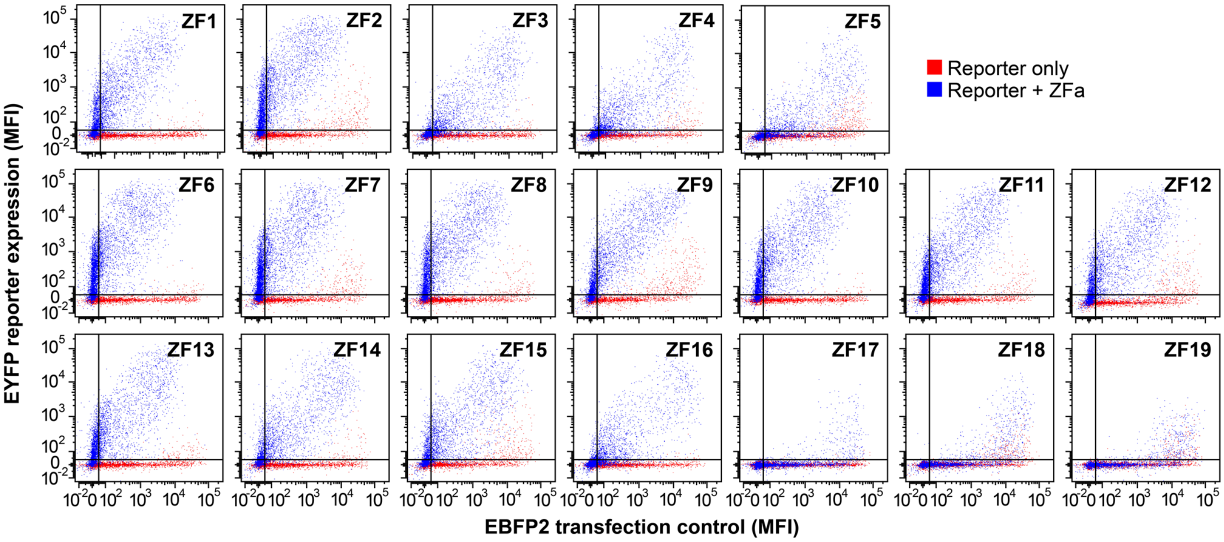


**Supplementary Fig. 4 Characterization of the panel of ZFa.** (a) Representative flow cytometry plots for **Fig. 3a**. We note several observations. (1) ZFa-induced EFYP increases with the transfection control protein (EBFP2). (2) Reporter-ZFa pairs that induce similar gene expression in the presence of a ZFa can vary in ZFa-independent gene expression (e.g., ZF6a and ZF7a). Thus, a different *b* (background) term in the COMET model is used for each promoter. (3) For many conditions, some cells that would be considered "not transfected" based on EBFP2 expression do express EYFP (e.g., the ZF2a case), indicating that ZFa are potent even when cells receive low amounts of plasmid. (4) Squelching is suggested by the observed decrease in EBFP2 expression between the "reporter only" and "reporter + ZFa" conditions for ZFa that induce high reporter expression.

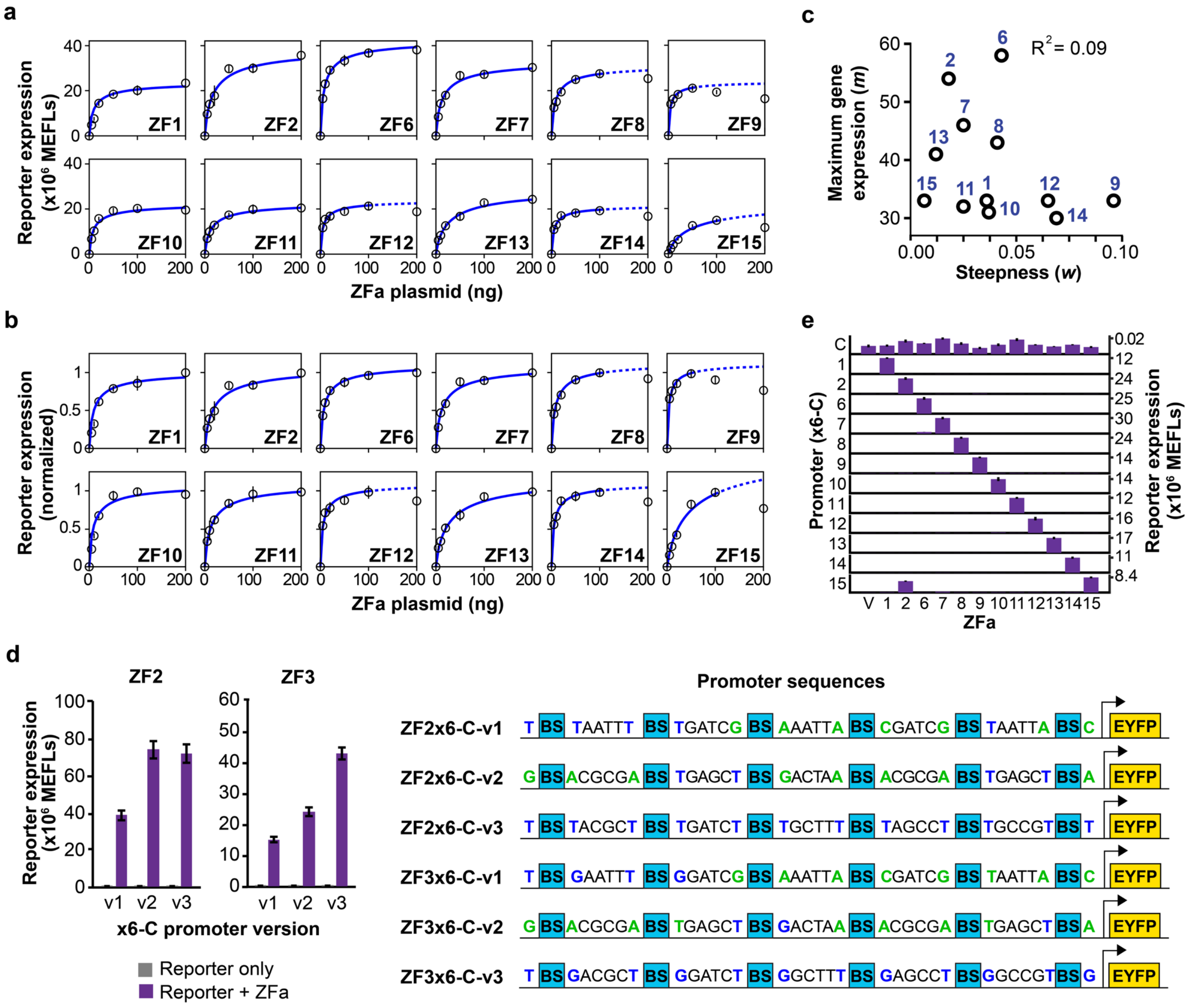


**Supplementary Fig. 5 Characterization of the panel of ZFa**. (a–c) Dose responses and model fits for the twelve strongest ZFa on x6-C promoters. Data are shown in MEFLs in (a) and normalized to the maximum of each reporter in (b). (c) Correlations in fitted parameters *m* and *w* for the twelve strongest ZFa on x6-C promoters. There is no significant correlation between *m* and *w* across the panel of ZFa (two-tailed permutation test $p = 0.34$). (d) ZFa induce transcription to varying extents depending on the flanking nucleotides. *(Right)* Comparison of compact promoters containing six binding sites with different nucleotides in the intervening base pairs between binding sites, including different flanking nucleotides. Blue denotes flanking nucleotides that were previously reported to confer strong ZF binding[2]. Green denotes flanking nucleotides with previously unreported effects. "BS" indicates the ZF binding site. *(Left)* For ZF2a and ZF3a, reporter expression varied with changes to spacer sequences. Reporter v1 constructs were not used elsewhere in this work, reporter v2 constructs were used in **Fig. 3a** and **Fig. 3c**, and reporter v3 constructs were used in **Fig. 3b**. Gene expression was significantly affected by the version of the ZF2 reporter used (ANOVA $p < 0.001$) and the version of the ZF3 reporter used (ANOVA $p < 0.001$). (E) Bar graph representation of the data in **Fig. 3c**; each series is plotted on the range from null reporter expression (0 MEFLs, horizonal line) to the value indicated. Error bars depict S.E.M.

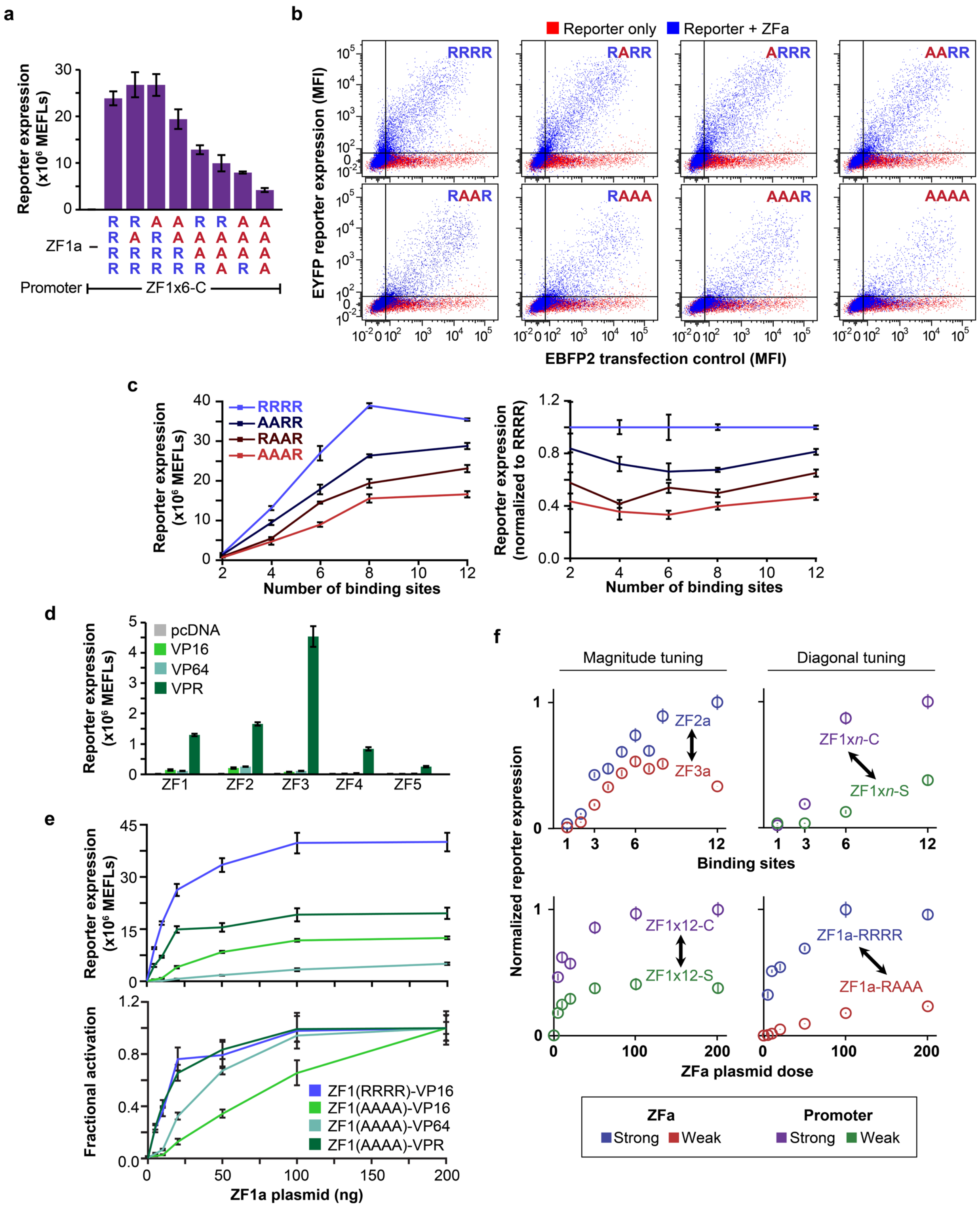
a
Reporter expression (x10⁶ MEFLs)
ZF1a –
Promoter
ZF1x6-C
b
Reporter only
Reporter + ZFa
RRRR
RARR
ARRR
AARR
RAAR
RAAA
AAAR
AAAA
EYFP reporter expression (MFI)
EBFP2 transfection control (MFI)
c
Reporter expression (x10⁶ MEFLs)
Reporter expression (normalized to RRRR)
Number of binding sites
d
pcDNA
VP16
VP64
VPR
ZF1
ZF2
ZF3
ZF4
ZF5
e
Fractional activation
ZF1(RRRR)-VP16
ZF1(AAAA)-VP16
ZF1(AAAA)-VP64
ZF1(AAAA)-VPR
ZF1a plasmid (ng)
f
Magnitude tuning
Diagonal tuning
ZF2a
ZF3a
ZF1xn-C
ZF1xn-S
Binding sites
Normalized reporter expression
ZF1x12-C
ZF1x12-S
ZF1a-RRRR
ZF1a-RAAA
ZFa plasmid dose
ZFa
Strong
Weak
Promoter
Strong
Weak

**Supplementary Fig. 6 Properties of ZFa with ZF mutants and AD variants.** (a) Effect of DNA affinity mutations on ZFa-induced reporter expression, using 100 ng of ZFa plasmid. ZF mutations affected reporter expression (ANOVA $p < 0.001$). (b) Representative flow cytometry plots from (a); the "reporter only" sample is the same across panels. (c) Effects of ZF mutations on reporter expression using promoters with varying numbers of binding sites. The WT ZF1a and three mutant ZF1a variants were each transfected along with compact promoters containing different numbers of binding sites. (*Left*) Reporter expression is presented in absolute units. Each ZFa reaches essentially maximum reporter expression with promoters containing 8–12 binding sites. Gene expression was affected by both the number of ZF binding sites (two-factor ANOVA $p < 0.001$) and the ZF mutations ($p < 0.001$). (*Right*) Reporter expression was normalized to the expression at each dose from the WT case. (d) VP16, VP64, and VPR activation domains (ADs) were fused to the panel of five ZF domains characterized in **Fig. 1b** and transfected into cells with a reporter plasmid containing a cognate x1 promoter driving an EYFP reporter. For each ZFa, VPR led to more reporter expression than VP16, and VP64 was similar to VP16. The ordering of reporter expression with VP16-ZF did not directly correspond to the ordering of reporter expression with VPR-ZF. Reporter expression was affected by both the choice of ZF (two-factor ANOVA $p < 0.001$) and the AD ($p < 0.001$), with an interaction between these two variables ($p < 0.001$). (e) Reporter expression data from **Fig. 4f** are shown with a linear x-axis, in absolute units (*left*) or normalized units (*right*). For each ZFa series, data were normalized to the maximum expression observed for the ZFa. The TFs vary in dose response profiles: WT ZF-VP16 and mutant ZF-VPR both reached maximal reporter expression at relatively low doses, mutant ZF-VP64 had a rare sigmoidal response, and mutant ZF-VP16 had a large linear range. The results show how the AD variant and ZF mutant can be chosen to customize response profiles. (f) Representative experimental datasets, corresponding to the summary schematic in **Fig. 4g**, were obtained from **Figs. 2a, 3b, and 4b**. Error bars depict S.E.M.

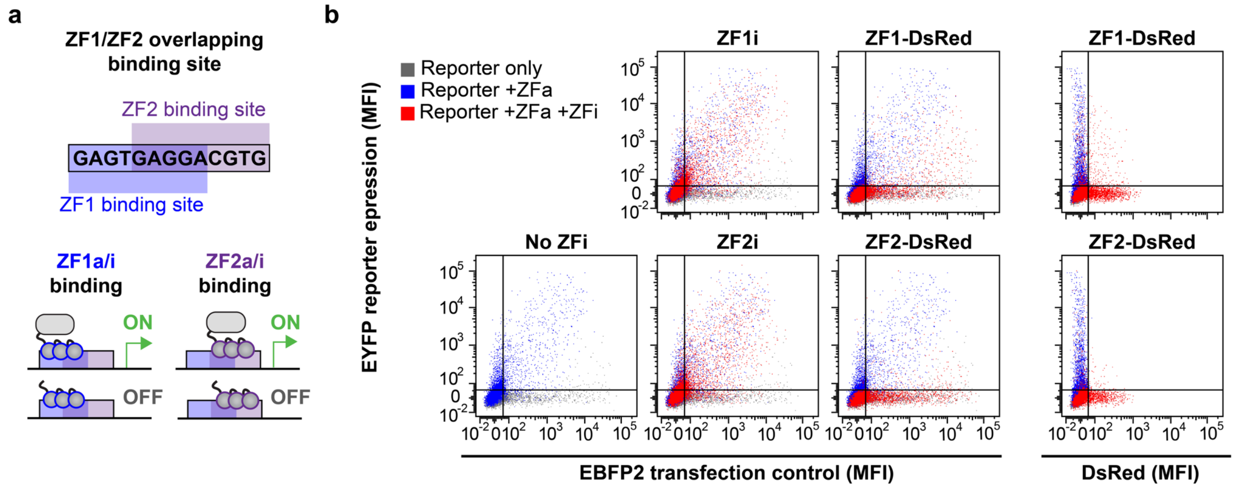


**Supplementary Fig. 7 Investigation of ZFi-mediated and ZFi-DsRed-mediated inhibition.** (a) The cartoon depicts a hybrid promoter regulated by a ZFa and a ZFi. Shown are potential states that any one overlapping binding site could take. The promoters that were evaluated experimentally contain multiple hybrid binding sites. The last five nucleotides of the ZF1 binding site are the same as the first five of the ZF2 binding site, and this property was used to construct a hybrid reporter by arranging six sites for ZF1 each separated by six base pairs, where the first four base pairs of this linker are the last four base pairs of the ZF2 site. (b) Representative flow cytometry plots from **Fig. 5b**.

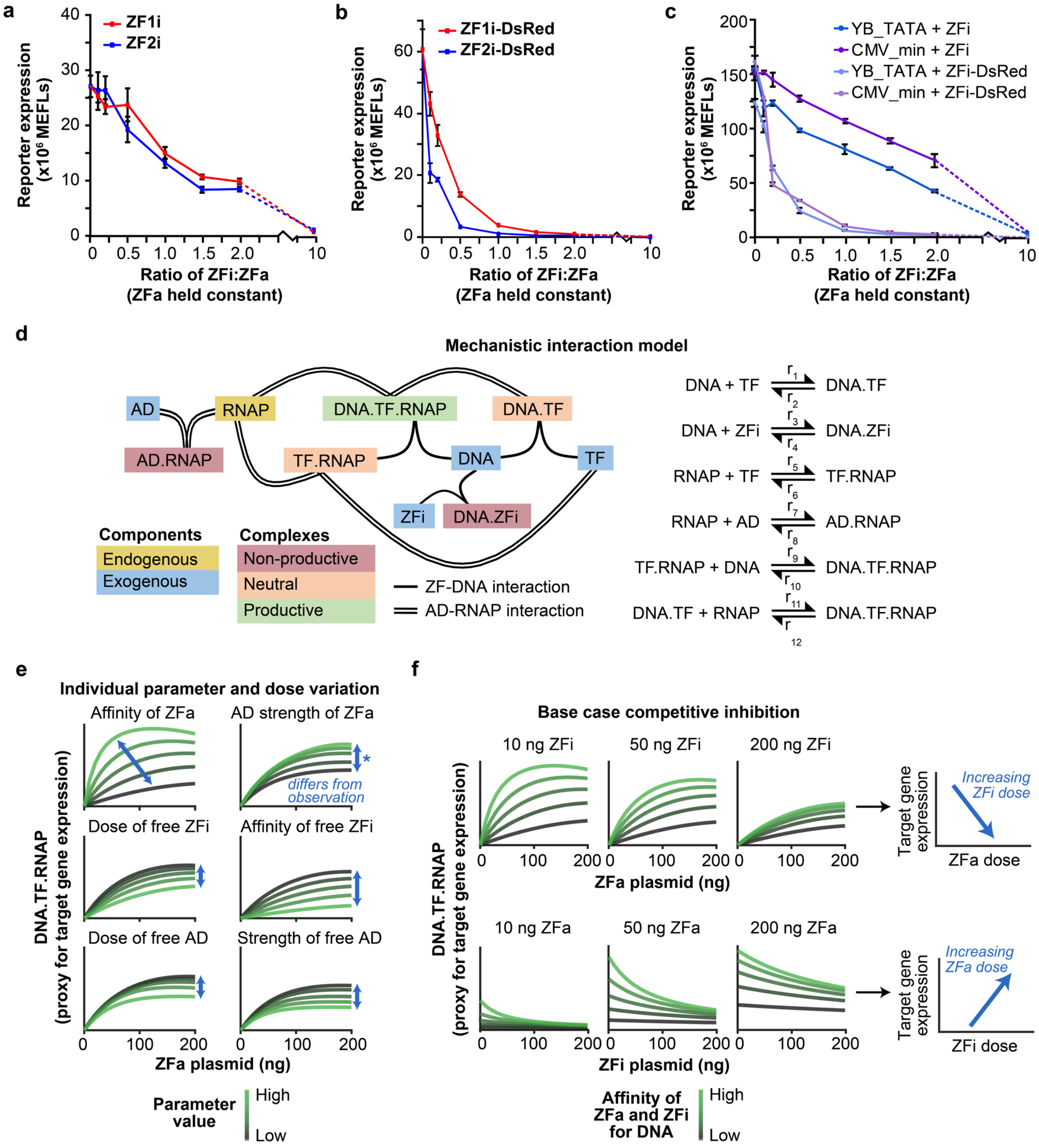
a
ZF1i
ZF2i
Reporter expression (x10⁶ MEFLs)
Ratio of ZFi:ZFa (ZFa held constant)
b
ZF1i-DsRed
ZF2i-DsRed
Reporter expression (x10⁶ MEFLs)
Ratio of ZFi:ZFa (ZFa held constant)
c
YB_TATA + ZFi
CMV_min + ZFi
YB_TATA + ZFi-DsRed
CMV_min + ZFi-DsRed
Reporter expression (x10⁶ MEFLs)
Ratio of ZFi:ZFa (ZFa held constant)
d
Mechanistic interaction model
AD
RNAP
DNA.TF.RNAP
DNA.TF
AD.RNAP
TF.RNAP
DNA
TF
ZFi
DNA.ZFi
Components
Endogenous
Exogenous
Complexes
Non-productive
Neutral
Productive
ZF-DNA interaction
AD-RNAP interaction
DNA + TF ⇌ DNA.TF (r1, r2)
DNA + ZFi ⇌ DNA.ZFi (r3, r4)
RNAP + TF ⇌ TF.RNAP (r5, r6)
RNAP + AD ⇌ AD.RNAP (r7, r8)
TF.RNAP + DNA ⇌ DNA.TF.RNAP (r9, r10)
DNA.TF + RNAP ⇌ DNA.TF.RNAP (r11, r12)
e
Individual parameter and dose variation
Affinity of ZFa
AD strength of ZFa
differs from observation
Dose of free ZFi
Affinity of free ZFi
Dose of free AD
Strength of free AD
DNA.TF.RNAP (proxy for target gene expression)
ZFa plasmid (ng)
Parameter value
High
Low
f
Base case competitive inhibition
10 ng ZFi
50 ng ZFi
200 ng ZFi
ZFa plasmid (ng)
Target gene expression
Increasing ZFi dose
ZFa dose
10 ng ZFa
50 ng ZFa
200 ng ZFa
ZFi plasmid (ng)
Increasing ZFa dose
ZFi dose
DNA.TF.RNAP (proxy for target gene expression)
Affinity of ZFa and ZFi for DNA
High
Low

**Supplementary Fig. 8 Further investigation of ZFi-mediated and ZFi-DsRed-mediated inhibition and the COMET mechanism.** (a, b) ZFi (a) and ZFi-DsRed (b) dose responses. In these experiments, the doses of ZFa plasmid and reporter plasmid were held constant while the dose of inhibitor plasmid was increased. Increasing the dose of ZFi led to decreased reporter expression (two-factor ANOVA $p < 0.001$), while the choice of ZF domain did not ($p = 0.35$). Increasing the dose of ZFi-DsRed also led to decreased reporter expression (two-factor ANOVA $p < 0.001$) and using the ZF2 domain resulted in more potent inhibition than using ZF3 ($p < 0.001$). (c) Dose responses of ZFi and ZFi-DsRed on the ZF1x6-C promoter with either the YB_TATA or CMV minimal promoters, as in (a) and (b). Both ZFi and ZFi-DsRed inhibited the YB_TATA and stronger CMV minimal promoters (two-factor ANOVAs $p < 0.001$). In both cases, the CMV minimal promoter was less inhibited than the YB_TATA promoter for a given dose of ZFi or ZFi-DsRed ($p < 0.001$), possibly due to the higher *w* value conferred on the ZFa response by the CMV minimal promoter. (d) A model for gene expression that uses more mechanistic detail than does the concise COMET model. The schematic on the left depicts reversible associations and dissociations (lines) between components (color-coded) as a network, and the corresponding biochemical reactions are listed on the right. The DNA variable represents a reporter gene with a promoter containing one binding site (x1). ZFi and free AD species can form non-productive complexes with DNA and RNAP, respectively. Here, the RNAP variable represents the ensemble of endogenous factors involved in transcription, and not specifically RNAPII (which does not directly physically interact with the AD). (e, f) Steady-state simulated values of DNA.TF.RNAP are a proxy for reporter expression here. Simulations were run for a single cell, i.e., homogeneously rather than for a heterogeneous population. (e) The abundance and properties of ZFa, ZFi, and free AD (not implemented experimentally) are expected to have the effects shown on reporter expression for a x1 promoter. The arrow in each panel indicates whether tuning affects the dose response trend vertically or diagonally. Although the simulated diagonal tuning of ZFa affinity agrees with experiments in **Fig. 4**, the effects of AD strength do not match those observed (denoted by an asterisk)—in that the x6-compact (*multi-site*) promoter experiments and associated transfer function model fits show diagonal tuning while the *single-site* promoter simulations show vertical tuning. This difference is consistent with the phenomenon of cooperativity through TF-mediated RNAPII recruitment at multi-site promoters (**Fig. 2, Online Methods**). That is, for a cooperative (multi-site) promoter, when a weak activator is present at a sufficiently high dose, it can disrupt the dual mechanism by which the inhibitor operates, thereby greatly increasing transcription. Conversely, when a weak inhibitor is present at such a promoter at a sufficiently high dose, the inhibitor can disrupt activator binding via the dual mechanism, thereby greatly decreasing transcription. In both cases, the TF plasmid doses over which these transitions occur differ based upon TF strength, and therefore TF plasmid dose responses at cooperative promoters are described by diagonal tuning rather than vertical tuning. We note that additionally, at a very high dose or affinity of ZFa, the mechanistic interaction model used here shows a non-monotonic dose response due to the formation of TF.RNAP and DNA.TF at the expense of DNA.TF.RNAP. Non-monotonicity is considered a non-ideal behavior (observed in a subset of ZFa plasmid dose response experiments in **Supplementary Fig. 5a,b**); this is not represented in the concise COMET model but can be captured in the mechanistic interaction model used here. The lower two rows show the simulated effects as DNA and RNAP are sequestered into non-productive complexes. (f) Simulated effects of competition between an activator and inhibitor. With a x1 promoter, the inhibitor mechanism is competitive without loss of cooperativity. Rows show simulated dose responses with respect to one TF (ZFa or ZFi) while the other TF is held constant. The arrow in each row indicates the diagonal tuning that results from altering both the ZFa and ZFi affinity for DNA. Error bars depict S.E.M.

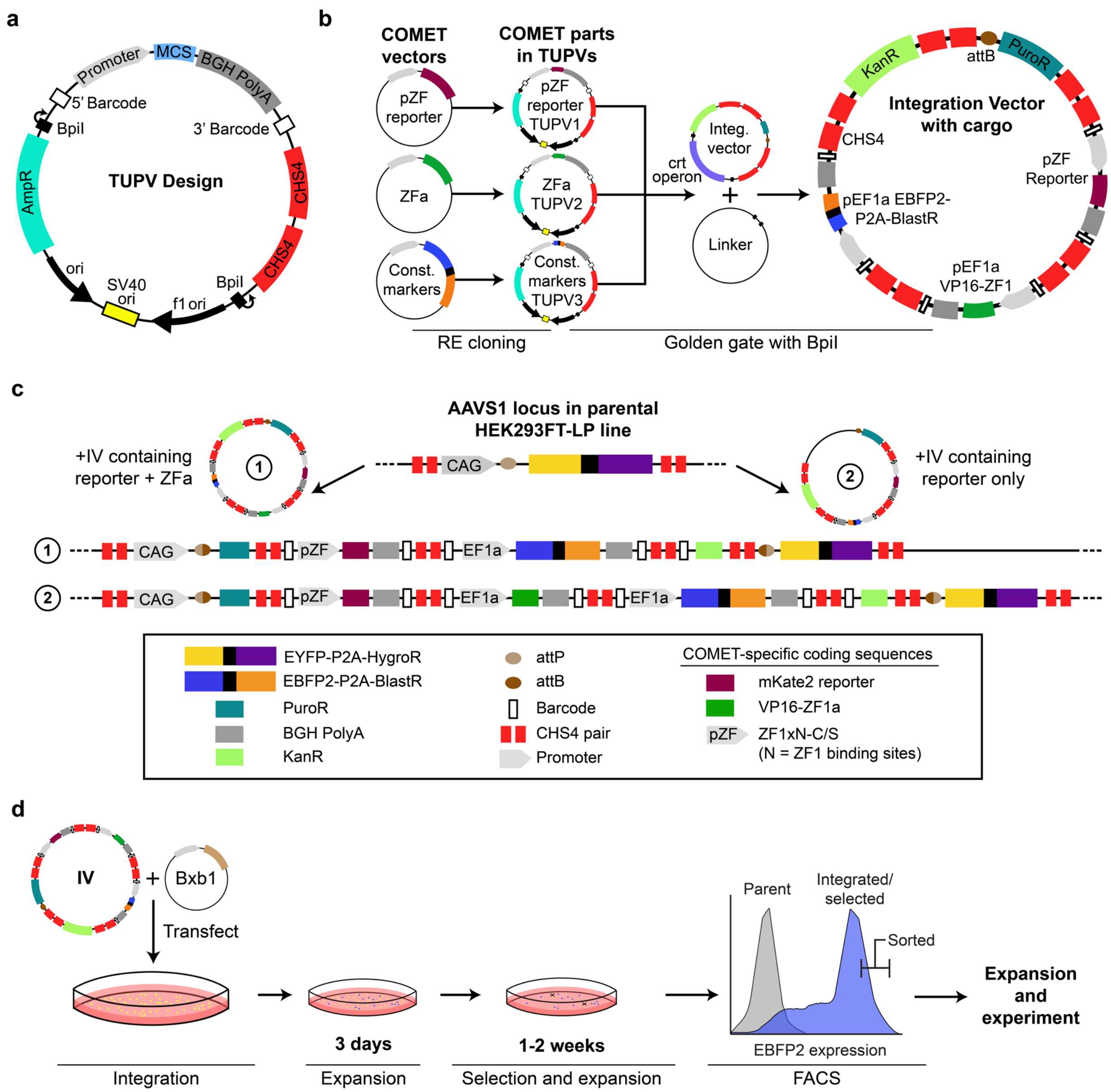
a
TUPV Design
Promoter
MCS
BGH PolyA
5' Barcode
3' Barcode
BpiI
AmpR
CHS4
CHS4
ori
SV40 ori
f1 ori
BpiI
b
COMET vectors
COMET parts in TUPVs
pZF reporter
pZF reporter TUPV1
ZFa
ZFa TUPV2
Const. markers
Const. markers TUPV3
crt operon
Integ. vector
Linker
Integration Vector with cargo
KanR
attB
PuroR
CHS4
pZF Reporter
pEF1a EBFP2-P2A-BlastR
pEF1a VP16-ZF1
RE cloning
Golden gate with BpiI
c
AAVS1 locus in parental HEK293FT-LP line
+IV containing reporter + ZFa
+IV containing reporter only
CAG
pZF
EF1a
EYFP-P2A-HygroR
EBFP2-P2A-BlastR
PuroR
BGH PolyA
KanR
attP
attB
Barcode
CHS4 pair
Promoter
COMET-specific coding sequences
mKate2 reporter
VP16-ZF1a
ZF1xN-C/S
(N = ZF1 binding sites)
d
IV
Bxb1
Transfect
3 days
1-2 weeks
Parent
Integrated/ selected
Sorted
EBFP2 expression
Expansion and experiment
Integration
Expansion
Selection and expansion
FACS

**Supplementary Fig. 9 Workflow for landing pad (LP) integration and stable cell line generation.** (a) The pcDNA-based transcription unit positioning vector (TUPV) permits transfer of COMET components into the mMoClo system[3] through restriction enzyme (RE) cloning. Notable features include the pcDNA multiple cloning site (MCS) for coding sequences and convenient RE sites placed at the 5' side of the promoter. Barcodes unique to each TUPV, found 5' of the promoter and 3' of the terminator, enable sequencing of the TUPV contents after TUPVs are combined into an integration vector[4]. Two repeats of the CHS4 insulator were included to mitigate transcriptional readthrough[5]. Details of these TUPVs and their features are in **Online Methods** and **Supplementary Tables 5, 6.** (b) After COMET and other components are transferred into TUPVs by RE cloning, they can be combined into a single plasmid with one to nine transcription units through a BpiI-mediated golden gate reaction. The integration vector serves as the backbone for the reaction and contains an attB site (for downstream integration into the LP) 5' of a promoter-less puromycin resistance gene, dual CHS4 insulators, a *crtRed* operon for Red-White colony screening that is replaced by the inserts from the TUPVs and Linker Vector, and dual CHS4 insulators to protect the 3' end of the LP locus after integration. The end result is the integration vector with cargo comprising multiple transcription units. (c) Integration of the LP cargo into the LP by BxB1-mediated recombination of the attB site on the IV and the genomic attP site integrates cargo into the AAVS1 safe harbor locus in the HEK293FT-LP cell line. Cells with successful integrations are no longer yellow (EYFP+) or hygromycin resistant; instead, they are blue (EBFP+) and resistant to puromycin and blasticidin. (d) After IV delivery by transient transfection and integration via BxB1, cells are expanded, selected with puromycin and blasticidin, and then sorted via FACS based upon EBFP2 expression (a typical gate used for this step is illustrated on the histogram). Additional details of this methodology and rationale are in **Online Methods**.

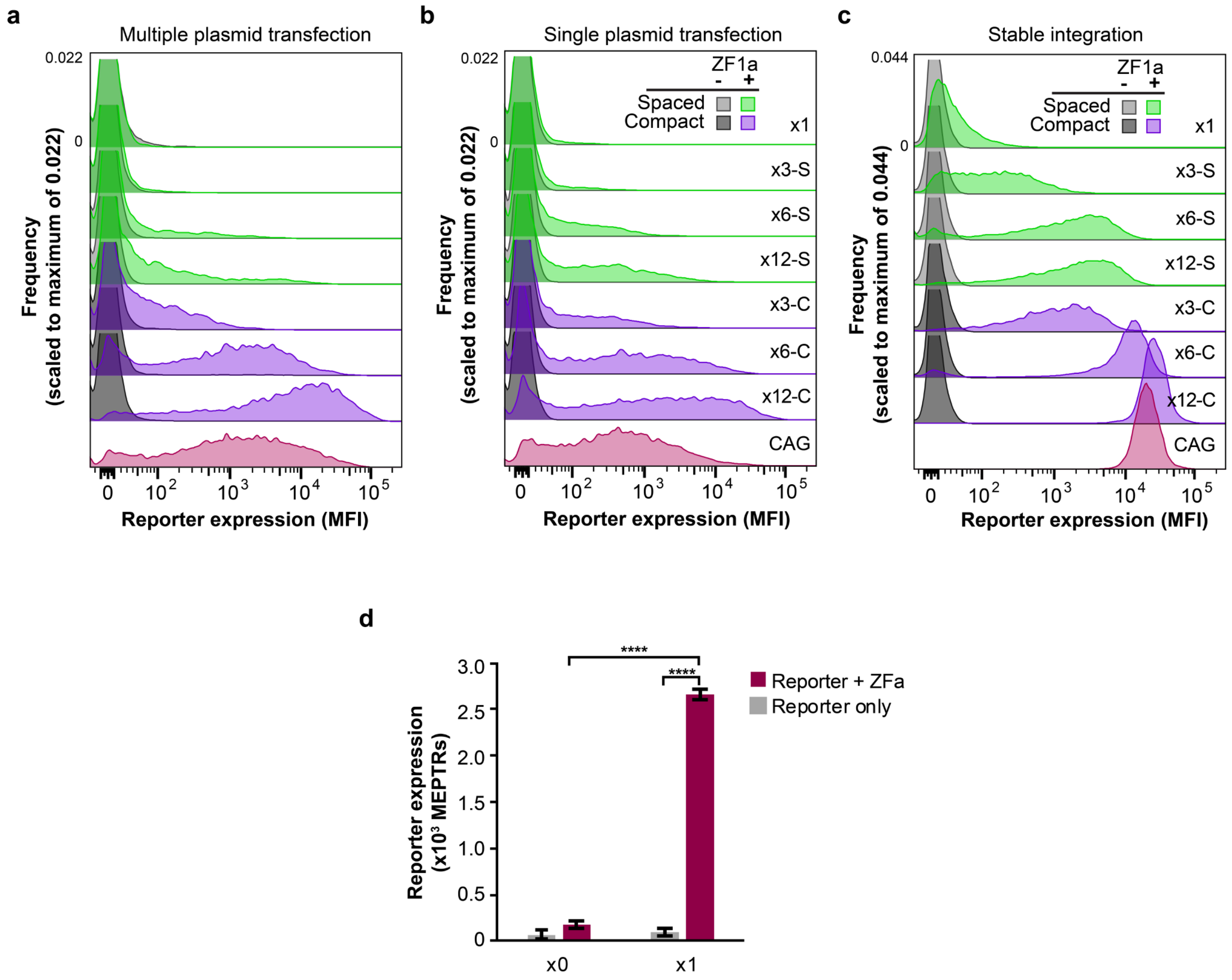


**Supplementary Fig. 10 Characterization of genomically integrated COMET TFs.** Representative histograms of reporter expression shown in **Fig. 6** are presented here with truncation of the y-axis (frequency) to better visualize differences between populations in each expression context: (a) multiple plasmid transfection, (b) single plasmid transfection, and (c) stable integration. Most trends observed in the stable genomic integration context appear to follow the trends observed in the context of transient transfection; these trends include an increase in reporter expression with addition of binding sites, and an increase in reporter expression for compact versus spaced promoter architectures. (d) A genomically-integrated promoter containing a single integrated ZF1 binding site induces low but significant reporter expression compared to background (one-tailed Welch's *t*-test ****$p < 0.0001$). All cell lines except for the ZF1x0 +ZFa line were sorted by FACS (outlined in **Online Methods**) before expansion and assaying. Data in all panels reflect reporter expression for cells that express EBFP2—a constitutive fluorescent marker driven from a gene that is genomically integrated with the cargo. Error bars depict S.E.M.

**a**

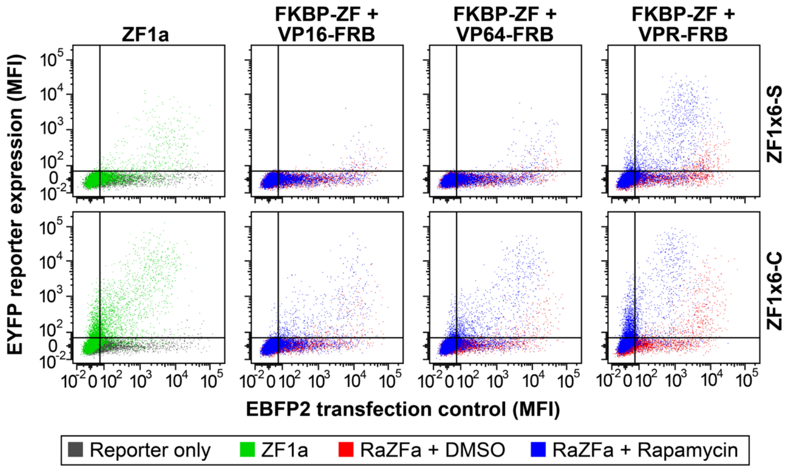


**Supplementary Fig. 11 Characterization and tuning of RaZFa activity.** (a) Representative flow cytometry plots from **Fig. 7b**. Rapamycin induces gene expression via RaZFa from COMET promoters similar to how a ZFa induces gene expression from these promoters. The effects of AD strength, as first seen in **Fig. 4**, apply similarly to RaZFa.

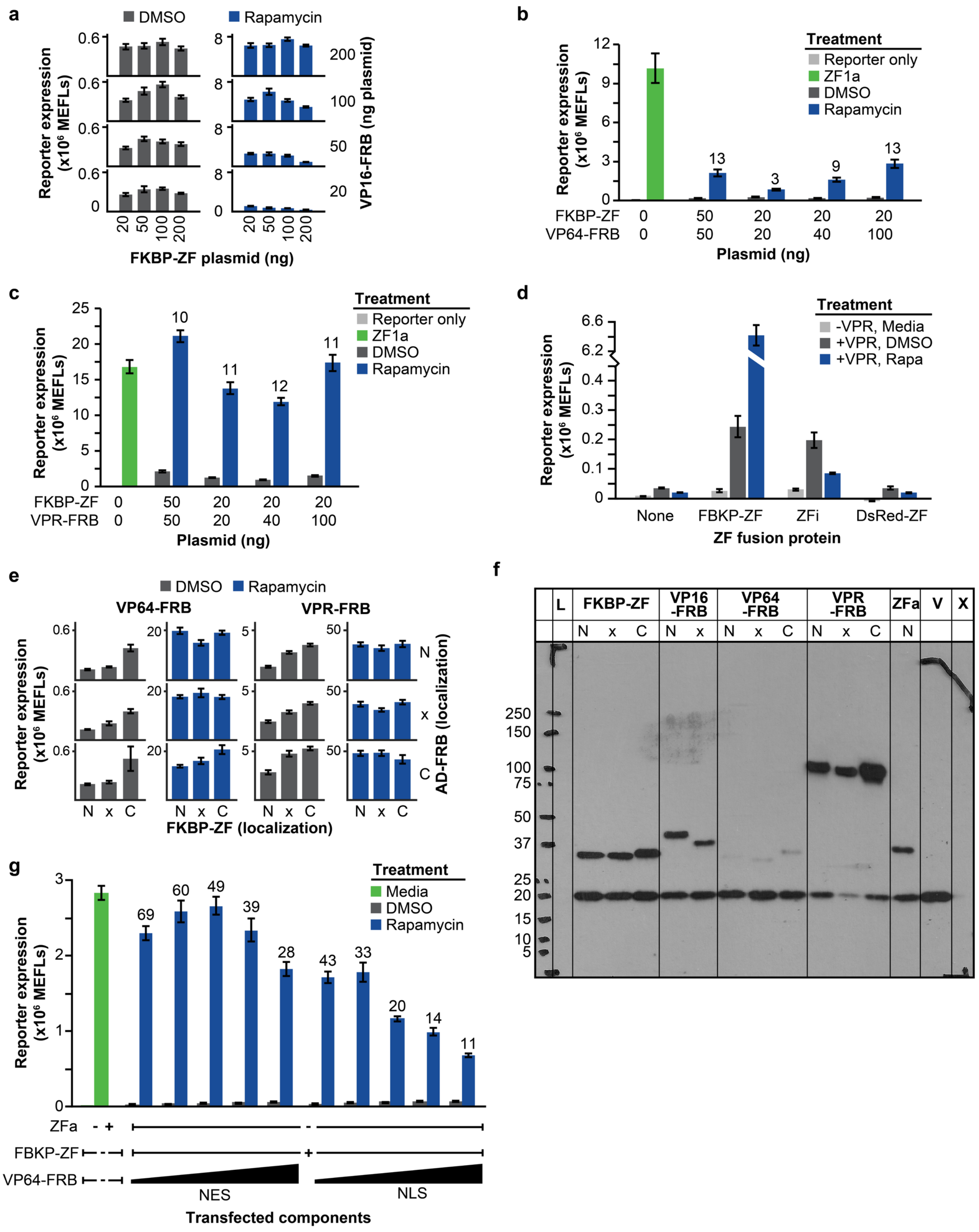
a
DMSO
Rapamycin
Reporter expression (x10^6 MEFLs)
0.6
0
8
200
100
50
20
VP16-FRB (ng plasmid)
20 50 100 200
FKBP-ZF plasmid (ng)
b
Treatment
Reporter only
ZF1a
DMSO
Rapamycin
12
9
6
3
0
13
3
9
13
FKBP-ZF 0 50 20 20 20
VP64-FRB 0 50 20 40 100
Plasmid (ng)
c
Treatment
Reporter only
ZF1a
DMSO
Rapamycin
25
20
15
10
5
0
10
11
12
11
FKBP-ZF 0 50 20 20 20
VPR-FRB 0 50 20 40 100
Plasmid (ng)
d
Treatment
-VPR, Media
+VPR, DMSO
+VPR, Rapa
6.6
6.2
0.4
0.3
0.2
0.1
0
None
FBKP-ZF
ZFi
DsRed-ZF
ZF fusion protein
e
DMSO
Rapamycin
VP64-FRB
VPR-FRB
0.6
20
5
50
N
x
C
AD-FRB (localization)
N x C
FKBP-ZF (localization)
f
L
FKBP-ZF
VP16 -FRB
VP64 -FRB
VPR -FRB
ZFa
V
X
N x C N x N x C N x C N
250
150
100
75
50
37
25
20
15
10
5
g
Treatment
Media
DMSO
Rapamycin
3
2
1
0
69
60
49
39
28
43
33
20
14
11
ZFa - +
FBKP-ZF
+
VP64-FRB
NES
NLS
Transfected components

**Supplementary Fig. 12 Characterization and tuning of RaZFa activity.** (a) Data from **Fig. 7c** are shown as a bar graph. (b–c) Effect of component doses on the performance of (b) VP64-based RaZFa and (c) VPR-based RaZFa. Unlike the case of VP16-based RaZFa (**Fig. 7c**), decreasing the FKBP-ZF dose and increasing the AD-FRB dose did not lead to an increase in fold induction for VPR-based RaZFa. For all RaZFa, reporter expression was significantly higher with rapamycin than with the DMSO vehicle (one-tailed Welch's *t*-test, all $p < 0.01$). (d) Investigation of rapamycin-independent RaZFa activity. With a ZF1x6-C promoter, VPR-FRB was transfected alone or with FKBP-ZF, ZFi, or ZFi-DsRed, and cells were treated with rapamycin or DMSO vehicle. (e) Data from **Fig. 7d** are shown as a bar graph. (f) Western blot of RaZFa components with tags for subcellular localization (N: nuclear, NLS; C: cytoplasmic, NES). All components (upper bands) contain an N-terminal 3x-FLAG tag. Cells were co-transfected with a 3x-FLAG-tagged NanoLuciferase (lower band) as a loading and transfection control. VP64-FRB is expressed less than the other AD-FRB fusions, and VPR-FRB is expressed more. The NES tag increases protein expression level. (g) Increasing the dose of VP64-FRB above the doses tested in (b). To investigate whether the low expression of VP64-FRB limited the rapamycin-induced activation of reporter expression, VP64-FRB with a NES or NLS was transfected at increasing doses with FKBP-ZF (20 ng of FKBP-ZF; 100, 200, 400, 600, or 800 ng of VP64-FRB). While rapamycin-inducible reporter expression increased with VP64-FRB dose, so did rapamycin-independent reporter expression, resulting in lower fold induction at high VP64-FRB doses. At high doses, squelching (as evidenced by decreased rapamycin-inducible reporter expression) was also evident and was more pronounced for the NLS-tagged than the NES-tagged VP64-FRB. For all RaZFa, reporter expression was significantly higher in the presence of rapamycin compared to the DMSO vehicle cases (one-tailed Welch's *t*-test, all $p < 0.01$). Error bars depict S.E.M.

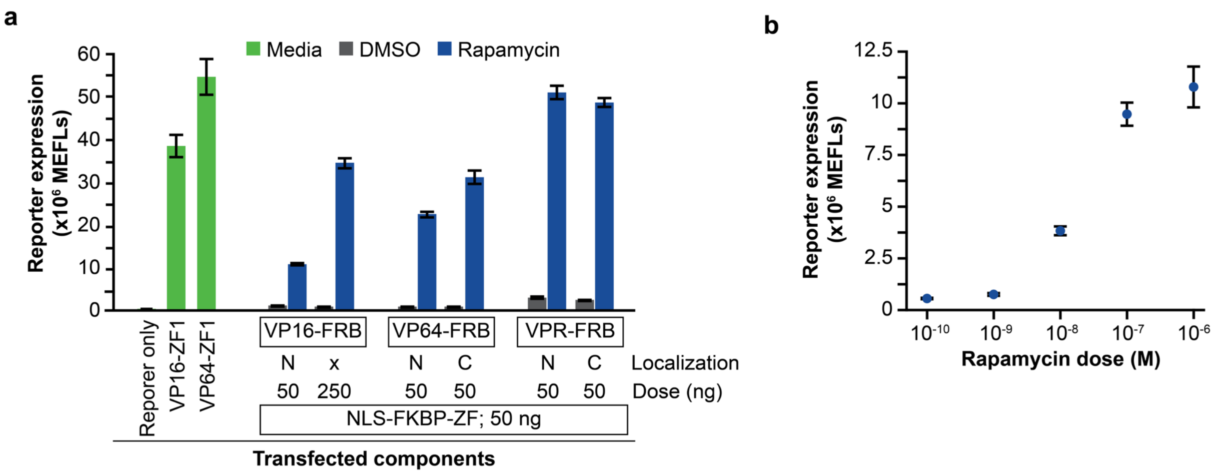


**Supplementary Fig. 13 Results of RaZFa tuning.** (a) Comparison of initial RaZFa performance with performance after optimization. Cells were transfected with various doses of RaZFa components, with various subcellular localization tags. For VP16-based RaZFa, increasing the dose of the VP16-FRB component five-fold and removing the localization tag increased reporter fold induction from 11 to 38 (~360% increase). For VP64-based RaZFa, localizing the VP64-FRB component to the cytoplasm rather than the nucleus increased fold induction from 25 to 40 (60% increase). For VPR-based RaZFa, localization of the VPR-FRB domain to the cytoplasm had a more modest effect on fold induction (17 to 20; 18% increase). Despite the substantial induced state, VPR-based RaZFa also exhibit high background. (b) Rapamycin dose response of VP64-based RaZFa. The range for inducibility spans approximately 1 nM to 1 μM rapamycin.

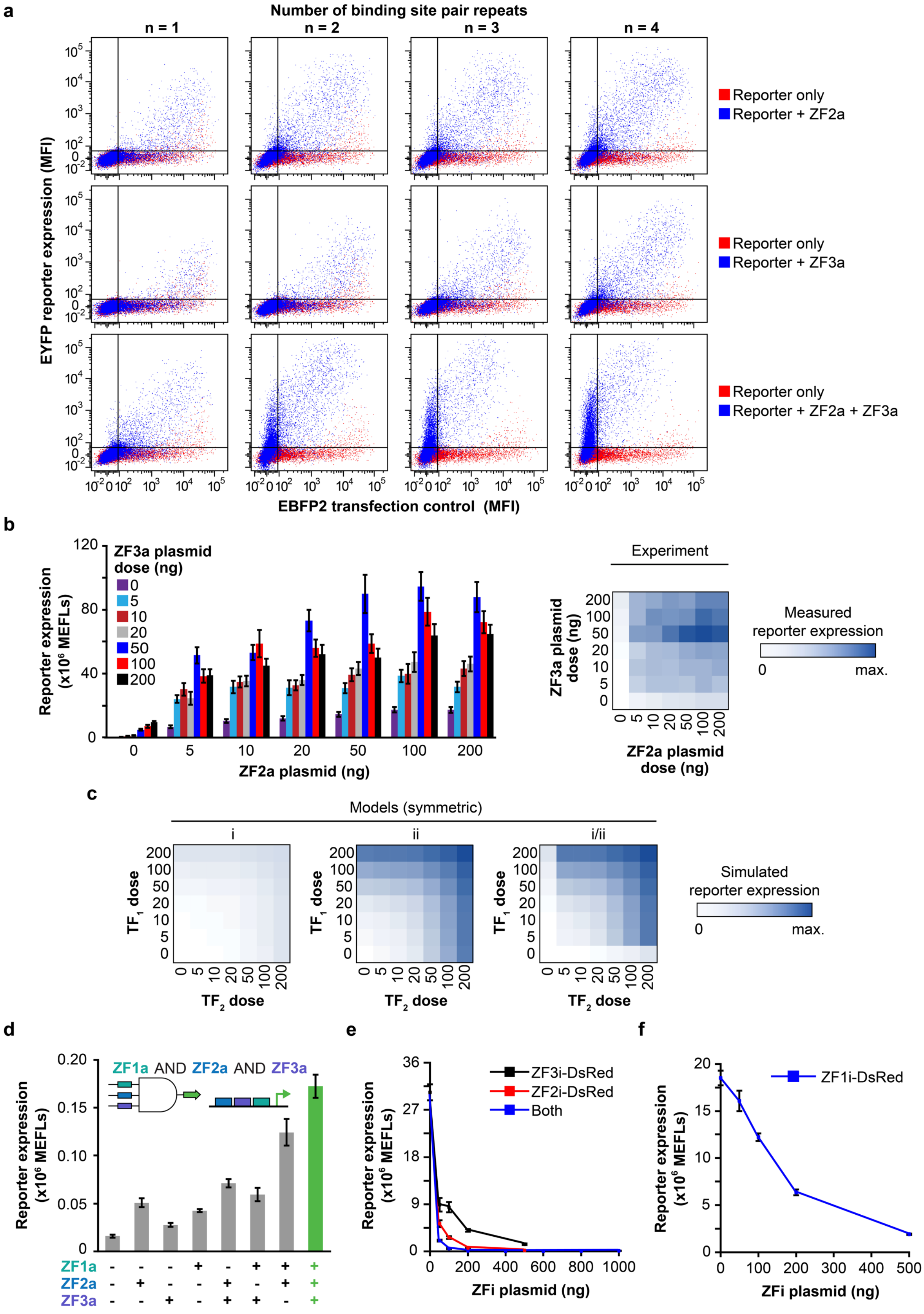
a
Number of binding site pair repeats
n = 1
n = 2
n = 3
n = 4
EYFP reporter expression (MFI)
EBFP2 transfection control (MFI)
Reporter only
Reporter + ZF2a
Reporter only
Reporter + ZF3a
Reporter only
Reporter + ZF2a + ZF3a
b
ZF3a plasmid dose (ng)
Reporter expression (x10^6 MEFLs)
ZF2a plasmid (ng)
Experiment
ZF3a plasmid dose (ng)
ZF2a plasmid dose (ng)
Measured reporter expression
0
max.
c
Models (symmetric)
i
ii
i/ii
TF1 dose
TF2 dose
Simulated reporter expression
0
max.
d
ZF1a AND ZF2a AND ZF3a
Reporter expression (x10^6 MEFLs)
ZF1a
ZF2a
ZF3a
e
ZF3i-DsRed
ZF2i-DsRed
Both
Reporter expression (x10^6 MEFLs)
ZFi plasmid (ng)
f
ZF1i-DsRed
Reporter expression (x10^6 MEFLs)
ZFi plasmid (ng)

**Supplementary Fig. 14 Implementing Boolean logic with COMET.** (a) Representative flow cytometry plots from **Fig. 8b**. The "reporter only" sample is the same across panels for a given promoter. (b) Data from **Fig. 8c** are presented as a bar graph and a heat map. (c) Model simulations corresponding to **Fig. 8d** are displayed as heatmaps. These simulations depict different types of AND gates broadly (**Online Methods**) and are not parameterized to data. The hybrid model (i/ii) depicting a transition from x3-S along the perimeter to x6-C in the interior of the landscape captures COMET AND gate behavior better than standard non-hybrid models. As discussed in **Supplementary Fig. 8**, this compact model does not encompass mechanisms that could lead to squelching, which is why experimentally observed diminishment in reporter output at high ZFa doses is not captured. (d) Three-input promoter with one repeat of the three-site motif. Cells were transfected with the reporter (containing one site each for ZF1, ZF2, and ZF3) and combinations of the ZFa plasmids. The difference between any two ZFa and all three ZFa was modest; this promoter is not ideal for implementing Boolean logic, since the *effective architecture* switches only from x2-S or x2-C to x3-C (when both ZFa inputs are present, rather than one or the other). AND gate activation is considered statistically significant if reporter expression with all three ZFa present is greater than the sum of reporter expression with the three ZFa expressed individually, and greater than the sum of reporter expression with two ZFa co-expressed and the other expressed individually. This criterion was met for all conditions tested (one-tailed Welch's *t*-test, $p < 0.05$) except for (ZF3a)+(ZF1a+ZF2a) vs. (ZF1a+ZF2a+ZF3a) ($p = 0.17$). (e, f) Inhibiting the three-repeat AND gate using ZFi-DsRed. The promoter has alternating sites for ZF2 and ZF3 and employs the strategy shown in **Supplementary Fig. 7a**, in which a ZF2 site overlaps with a ZF1 site. Plasmids for ZF2a, ZF3a, the reporter, and ZFi-DsRed (dose response) were transfected into cells. Although high doses of ZF1i-DsRed were necessary for inhibition, ZF2i-DsRed and ZF3i-DsRed had more potent effects, and ZF2i-DsRed and ZF3i-DsRed in combination conferred strong inhibition even at low plasmid doses. Results from this experiment informed the dose choices for the experiment in **Fig. 8f**, which uses 50 ng of each ZFa plasmid and 100 ng of each ZFi-DsRed plasmid. Error bars depict S.E.M.

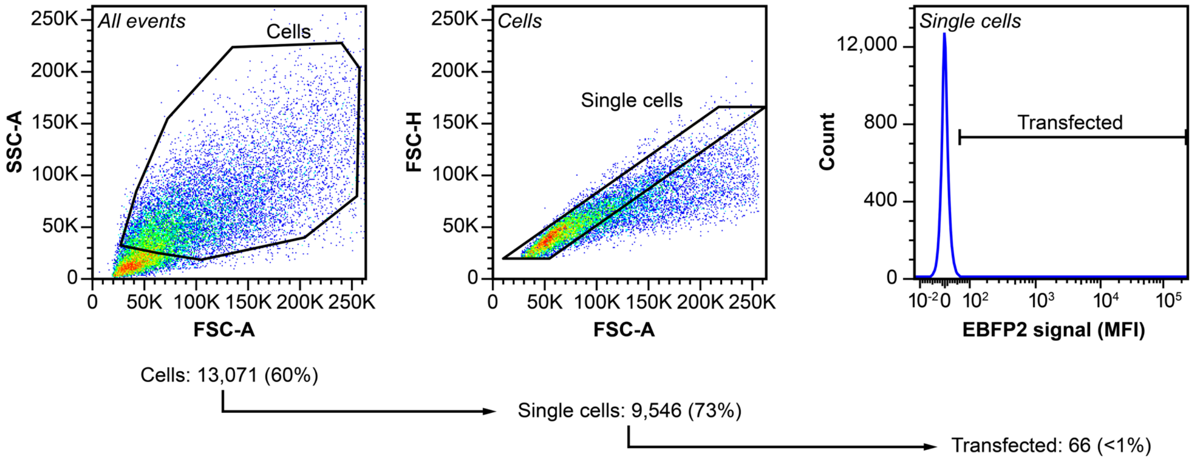


**Supplementary Fig. 15 Flow cytometry gating.** The plots show a sample of cells transfected with the modified pcDNA vector used in this study without any coding sequence in the multiple cloning site. These cells were *not* transfected with an EBFP2 expression plasmid. In the gating procedure, HEK293FT cells were identified based on the FSC-A vs SSC-A profile. From this population, single cells were identified based on the FSC-A vs FSC-H profile. The transfected population was defined as all single cells with a greater EBFP2 signal than the sample of single cells that was transfected with vector-only DNA (EBFP2 was used as a transfection control in this experiment and the vast majority of the experiments in this paper). This gate was drawn such that it did not encompass more than 1% of this non-fluorescent population of cells.

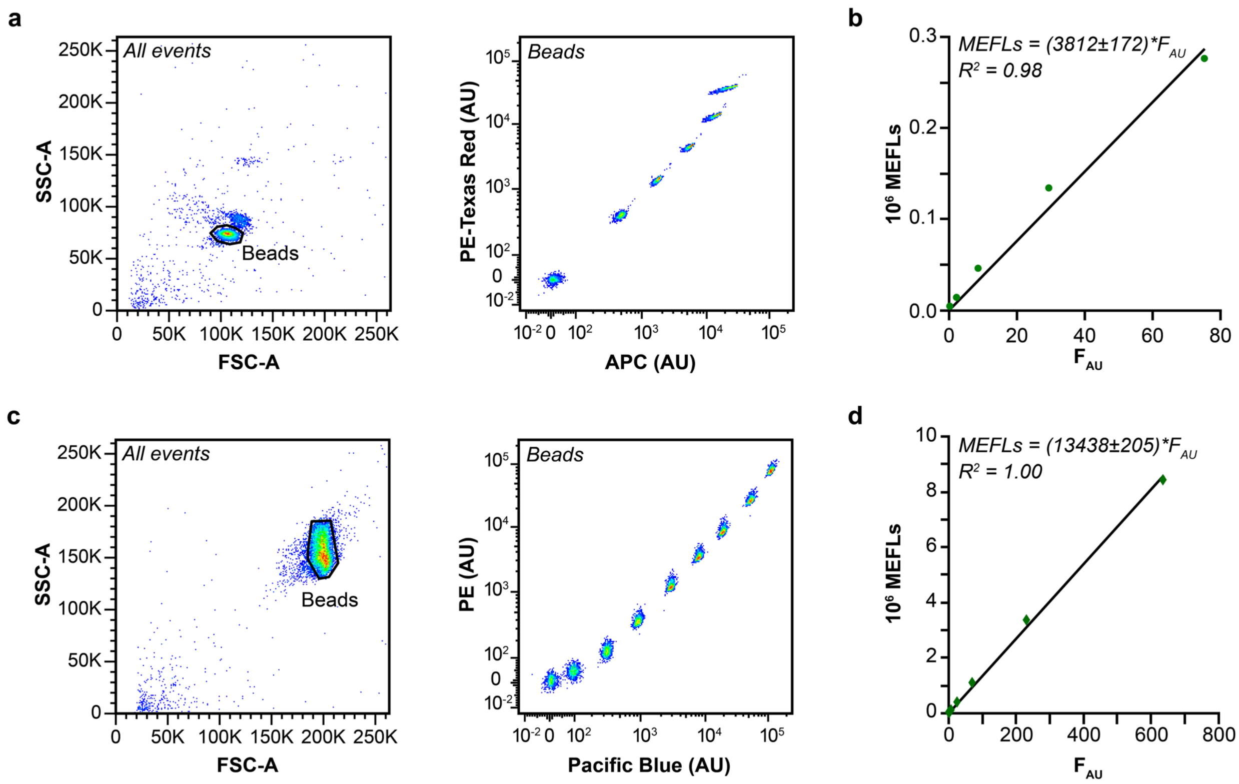


**Supplementary Fig. 16 Profile of fluorescent calibration beads.** Rainbow Calibration Particles (RCP; a, b) have six fluorescent bead populations, while UltraRainbow Calibration Particles (URCP; c, d) have nine fluorescent bead populations and are brighter than RCP. (a, c) Beads were identified based on the FSC-A vs. SSC-A profile. The beads are fluorescent in the majority of fluorescent channels on the flow cytometer. For each experiment, two channels were used to identify the bead populations. (b, d) The mean intensity of each population in the FITC (EYFP) channel in arbitrary units ($F_{AU}$) was recorded and plotted against the manufacturer-supplied number of fluorophores on the beads for each population (MEFLs). To generate the calibration curve, a linear regression was performed with the constraint that the y-intercept equals zero. In each experiment, MFI are converted to MEFLs by using the multiplier on $F_{AU}$ obtained from the regression. The magnitude of the multiplier, along with corresponding uncertainty (+/- one standard error), is reported in parentheses on each plot (b, d).

**Supplementary Table 9.** ZFa and fitted parameters for x6-C promoters.

<table>
<tr><th>Zinc finger[1]</th><th>Reference number[2]</th><th>Binding site[3]</th><th>b[5]</th><th>m[5]</th><th>w[5]</th></tr>
<tr><td>ZF1</td><td>43-8</td><td>a GAG TGA GGA c</td><td>0.08</td><td>33</td><td>0.036</td></tr>
<tr><td>ZF2</td><td>37-12</td><td>t GAG GAC GTG t</td><td>0.25</td><td>54</td><td>0.018</td></tr>
<tr><td>ZF3</td><td>158-2</td><td>t GTA GAT GGA g</td><td colspan="3" rowspan="3">n.d.</td></tr>
<tr><td>ZF4</td><td>97-4</td><td>a TTA TGG GAG a</td></tr>
<tr><td>ZF5</td><td>92-1</td><td>a GAT GTA GCC t</td></tr>
<tr><td>ZF6</td><td>150-4</td><td>g GTG TAG GGG t</td><td>0.02</td><td>58</td><td>0.043</td></tr>
<tr><td>ZF7</td><td>172-5</td><td>a GGA GGG GCT c</td><td>0.11</td><td>46</td><td>0.025</td></tr>
<tr><td>ZF8[4]</td><td>173-3</td><td>a GAT GAA GCT g</td><td>0.07</td><td>43</td><td>0.041</td></tr>
<tr><td>ZF9[4]</td><td>42-10</td><td>a GAC GCT GCT c</td><td>0.46</td><td>33</td><td>0.096</td></tr>
<tr><td>ZF10</td><td>13-6</td><td>a GAA GAT GGT g</td><td>0.01</td><td>31</td><td>0.037</td></tr>
<tr><td>ZF11</td><td>36-4</td><td>c GAA GAC GCT g</td><td>0.08</td><td>32</td><td>0.025</td></tr>
<tr><td>ZF12[4]</td><td>62-1</td><td>g GCC GAA GAT a</td><td>0.15</td><td>33</td><td>0.065</td></tr>
<tr><td>ZF13</td><td>21-16</td><td>a TTA GAA GTG a</td><td>0.04</td><td>41</td><td>0.012</td></tr>
<tr><td>ZF14[4]</td><td>14-3</td><td>g GAC GAC GGC a</td><td>0.20</td><td>30</td><td>0.069</td></tr>
<tr><td>ZF15[4]</td><td>129-3</td><td>c GGG GAC GTC a</td><td>0.18</td><td>33</td><td>0.007</td></tr>
<tr><td>ZF16</td><td>54-8</td><td>a TGG GTG GCA t</td><td colspan="3" rowspan="4">n.d.</td></tr>
<tr><td>ZF17</td><td>55-1</td><td>c TGG GGT GCC c</td></tr>
<tr><td>ZF18</td><td>93-10</td><td>c TTT GTT GGC a</td></tr>
<tr><td>ZF19</td><td>151-1</td><td>t GCA GGA GGT g</td></tr>
</table>

[1]All ZFa are WT (RRRR) with the VP16 AD. n.d. = no data. ZFa are numbered by order of appearance in this work. This is the nomenclature used in another study[2]; we provide this information to facilitate cross-referencing.[3] Each ZF binds one nucleotide triplet indicated by upper-case letters. Lower-case letters indicate nucleotides flanking the binding site; we observed that these flanking residues may influence binding affinity (**Supplementary Fig. 5**). The flanking nucleotides listed here are known to confer strong binding affinity[2], however some constructs in this study contain other nucleotides (promoter sequences are provided in **Supplementary Table 2**).[4] These ZFa exhibited squelching at high ZFa plasmid doses. [5]**Supplementary Fig. 5** contains the response profiles that were used to fit these parameters.

**Supplementary Table 10.** Fitted parameters for modifications to promoter architecture and ZFa domains.

| Figure | ZF | AD | Promoter | *b* | *m* | *w* |
|---|---|---|---|---|---|---|
| 2a | ZF1 | VP16 | ZF1x1 | 0.08 | 1.0 | 0.036 |
| | | | ZF1x3-S | | 1.7 | |
| | | | ZF1x6-S | | 5.4 | |
| | | | ZF1x12-S | | 15 | |
| | | | ZF1x3-C | | 7.1 | |
| | | | ZF1x6-C v1 | | 33 | |
| | | | ZF1x12-C | | 41 | |
| S2d | ZF1 | VP16 | ZF1x6-C CMV_Min | 0.26 | 33 | 0.058 |
| | | | ZF1x6-C SV40_Min | 0.43 | 7.5 | 0.046 |
| 4b | ZF1(RARR) | VP16 | ZF1x6-C v1 | 0.08 | 26 | 0.018 |
| | ZF1(ARRR) | | | | 19 | 0.010 |
| | ZF1(AARR) | | | | 15 | 0.011 |
| | ZF1(RAAR) | | | | 13 | 0.0043 |
| | ZF1(RAAA) | | | | 13 | 0.0023 |
| | ZF1(AAAR) | | | | 7 | 0.0040 |
| | ZF1(AAAA) | | | | 7 | 0.0017 |
| 4f | ZF1(AAAA) | VP64 | ZF1x6-C v1 | 0.08 | 24 | 0.012 |
| | | VPR | | | 78 | 0.020 |

The following tables are included as Microsoft Excel workbook files:

**Supplementary Table 1.** List of vectors and reporter plasmid templates.

**Supplementary Table 2.** List of reporter plasmids.

**Supplementary Table 3.** List of plasmids for constitutive expression of ZFa.

**Supplementary Table 4.** List of plasmids for constitutive expression of ZFi.

**Supplementary Table 5.** List of plasmids for use in mMoClo.

**Supplementary Table 6.** Features of mMoClo plasmids.

**Supplementary Table 7.** List of plasmids for integration of COMET ZFa.

**Supplementary Table 8.** List of plasmids for constitutive expression of RaZFa components.

**Supplementary Table 11.** Plasmid doses used in experiments in Fig. 1 and related supplementary figures.

**Supplementary Table 12.** Plasmid doses used in experiments in Fig. 2 and related supplementary figures.

**Supplementary Table 13.** Plasmid doses used in experiments in Fig. 3 and related supplementary figures.

**Supplementary Table 14.** Plasmid doses used in experiments in Fig. 4 and related supplementary figures.

**Supplementary Table 15.** Plasmid doses used in experiments in Fig. 5 and related supplementary figures.

**Supplementary Table 16.** Plasmid doses used in experiments in Fig. 6 and related supplementary figures.

**Supplementary Table 17.** Plasmid doses used in experiments in Fig. 7 and related supplementary figures.

**Supplementary Table 18.** Plasmid doses used in experiments in Fig. 8 and related supplementary figures.

**Supplementary Note 1. Nomenclature used in this manuscript.**

The ZF sequences used here were previously described by Khalil and colleagues[2]. We refer to ZF domains as ZF1, ZF2, etc. by order of appearance in this work for clarity. **Supplementary Table 9** provides a summary to facilitate cross-referencing of this nomenclature with that which was previously used by Khalil and colleagues. Addgene pages associated with our plasmids refer to ZFs by the original nomenclature.[2]

For ZF fusion proteins (e.g., ZFi-DsRed or VP16-ZF), the order of domains in the name does not indicate whether the protein is an N-terminal or C-terminal fusion. DsRed-Express2, FKBP, and all ADs fused to a ZF domain were fused to the N-terminus of the ZF domain. For AD-FRB, the AD is fused to the N-terminus of the FRB domain. Unless explicitly stated otherwise, all ZF proteins and ZF fusion proteins begin with a 3X-FLAG tag and SV40 NLS. More detailed information on the sequence of these constructs is provided in **Supplementary Tables 3,4,8**. Throughout, we refer to DsRed-Express2 as “DsRed” for brevity; this abbreviation always refers to DsRed-Express2 and never to either the original DsRed or DsRed2.

**Supplementary Note 2. Code availability**

Code is provided in MATLAB format in the supplementary material. The COMET_README file contains a description of code, system requirements, installation, demo, and instructions for use.

**Supplementary Note 3. Plasmid maps**

Plasmid maps are provided as GenBank files.
